# Temporal analysis of tear fluid proteome reveals critical corneal repair events after photorefractive surgery

**DOI:** 10.1101/2025.07.11.664300

**Authors:** Nadege Feret, Solene ter Schiphorst, Jana Kindermans, Hannah Crowdy, Laura Fichter, Jerome Vialaret, Christophe Hirtz, Frederic Michon, Vincent Daien

## Abstract

Corneal epithelial wound healing is a complex and finely orchestrated process critical for restoring visual acuity following injury or surgery. Photorefractive keratectomy (PRK), a common refractive surgical procedure, provides an ideal clinical model to study this process. Here, we employed advanced proteomic analysis to comprehensively map the dynamic changes in the tear fluid proteome at distinct phases following PRK. Our findings revealed significant alterations in nearly 45% of the tear proteome, highlighting temporally distinct molecular signatures. Immediately post-injury, a robust but controlled anti-inflammatory response coincided with pronounced upregulation of protein synthesis and cellular stress-management pathways. Subsequently, at day three, the molecular landscape shifted toward sustained epithelial regeneration, extracellular matrix remodeling, and controlled inflammation resolution. By delineating these critical and temporally compartmentalized molecular events, this study identifies novel tear-based biomarkers indicative of corneal healing efficacy and provides essential insights into potential therapeutic targets for improving clinical outcomes in corneal wound healing and ocular surface disorders.

## Introduction

The cornea is a vital, transparent structure at the forefront of the eye, essential for focusing visual images onto the retina. Its outermost epithelial layer is particularly crucial, serving as a protective barrier against pathogens and injuries, maintaining hydration balance, and supporting optical clarity[1]. Any disruption or damage to the cornea initiates a complex wound healing process that is critical to restoring transparency and preserving visual acuity[2].

Defective corneal epithelial healing presents significant clinical challenges, potentially leading to prolonged visual impairment and chronic ocular discomfort[3]. Corneal wound healing is a highly coordinated physiological event encompassing several overlapping phases: initial inflammatory responses, cellular proliferation, migration, and tissue remodeling. Immediately after injury, cytokines and growth factors alter the corneal microenvironment, attracting inflammatory cells and initiating tissue repair[4]. Epithelial cells migrate swiftly to close wounds, while underlying keratocytes differentiate into activated fibroblasts or myofibroblasts, synthesizing extracellular matrix components crucial for tissue integrity restoration. Effective completion of this process is essential for restoring corneal optical properties and maintaining clear vision[5,6]. Various factors, however, may impede optimal healing, resulting clinically in corneal haze, scarring, recurrent erosions, or persistent ulcers[7,8]. Such complications significantly compromise visual acuity, potentially leading to chronic discomfort, diminished vision quality, or severe vision-threatening conditions[9,10].

In recent years, refractive surgery, particularly photorefractive keratectomy (PRK), has markedly increased due to growing demand for vision correction. Unlike LASIK, PRK involves the removal of the corneal epithelial layer, making efficient and complication-free wound healing essential for optimal visual outcomes. PRK employs advanced laser technology that enables precise epithelial removal and typically minimal recovery time, as supported by studies demonstrating favorable outcomes across diverse patient populations[11–13]. Despite its generally safe profile, delayed or impaired wound healing post-PRK poses significant risks. Approximately 5–10% of patients experience delayed epithelial healing, influenced by pre-existing ocular surface conditions, the extent of myopia corrected, and specific surgical techniques[14,15]. Additionally, the intraoperative use of mitomycin C, aimed at preventing scarring, can paradoxically delay epithelial recovery by impacting cell cycle regulation and basal lamina remodeling[16,17]. Persistent inflammation from defective healing further exacerbates tissue remodeling challenges, fostering myofibroblast development and subsequent corneal opacity and scarring[18,19]. With millions of PRK procedures conducted annually worldwide, optimizing postoperative recovery is increasingly crucial.

Corneal physiology and healing depend on the microenvironment, including the tear film[20], corneal secretome[21], and nerve-derived neurotrophic factors[22]. Understanding the complex biochemical interplay within the corneal microenvironment has become an essential area of ophthalmologic research. Recent advances in tear proteomics have unveiled the dynamic composition of tear fluid, identifying numerous proteins, lipids, and cytokines reflective of ocular surface health[23]. These proteomic analyses have highlighted critical roles for specific proteins and mediators during corneal epithelial repair processes. For instance, matrix metalloproteinase-9 (MMP-9) levels correlate with corneal wound healing efficacy and inflammatory responses, particularly in chronic epithelial defects[24,25]. Consequently, identifying biomarkers within the tear film indicative of epithelial repair requirements has become vital. Tear film proteomics is emerging as a valuable tool, providing insights into molecular changes postoperatively and facilitating the discovery of novel therapeutic targets aimed at enhancing healing outcomes[2,26].

Here, we employed advanced proteomic techniques to elucidate changes in tear film composition following PRK, identifying potential biomarkers that underpin successful corneal re-epithelialization and wound healing. Our analyses revealed that the ocular surface fluid molecular landscape is highly dynamics. While the early response to the PRK procedure tends to favor anti-inflammatory pathways, and a strong transcriptome remodelling, the late response is focused on the extracellular matrix deposition and epithelial achitechture.

Identifying specific biomarkers involved in corneal wound healing illuminates previously unrecognized molecular pathways and cellular requirements crucial for effective corneal epithelial regeneration. These biomarkers could substantially assist clinicians in tailoring postoperative care and developing targeted therapeutic strategies to promote healing, minimize discomfort, and address potential complications.

## Results

### Tear fluid proteome landscape is deeply modified after PRK procedure

The transition from basal to reflex tear production represents a pivotal phase in the corneal wound healing process, yet its molecular underpinnings remain incompletely characterized. To elucidate the proteomic shifts occurring during this transition, we performed a comprehensive analysis of the tear proteome in patients undergoing corneal epithelial injury induced by photorefractive keratectomy (PRK) (**Figure 1**).

**Figure 1.**
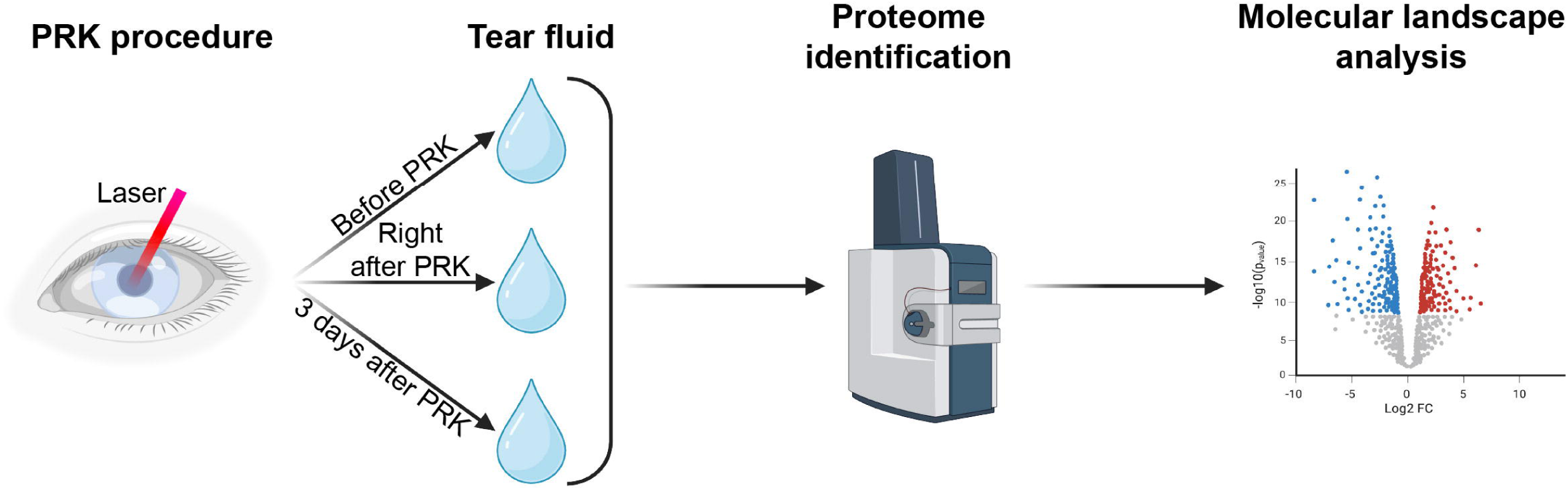
Experimental design. Tear samples were collected from both eyes of patients undergoing bilateral photorefractive keratectomy (PRK). The collection of tear fluid was performed at three distinct time points: immediately before surgery, immediately after surgery, and three days post-surgery. The tear samples were subsequently analyzed using mass spectrometry to characterize their protein composition. Finally, the molecular profiles obtained at these different time points were compared to elucidate changes associated with corneal wound healing.

Tear samples were collected from ten healthy adult patients (3 men and 7 women, aged 24–36 years; **Table 1**) without any history of ocular disease or prior treatment. Sampling was conducted at three time points: immediately before PRK (Pre), within one hour postoperatively (D0), and at a scheduled follow-up visit three days later (D3).

**Table 1.** Description of the patients. The demographic description of the patients.

| Patient | Age | Sex |
| --- | --- | --- |
| #1 | 29 | M |
| #2 | 34 | M |
| #3 | 24 | M |
| #4 | 36 | F |
| #5 | 34 | F |
| #6 | 28 | F |
| #7 | 29 | F |
| #8 | 24 | F |
| #9 | 31 | F |
| #10 | 31 | F |

Proteomic profiling revealed a complex and dynamic tear composition, with individual samples containing between 1,100 and 1,980 proteins. In total, 2,025 unique proteins were identified across all samples. The PCA clusterization of our samples has shown a close correlation of the Pre and D3 samples, compared to the D0 (**Figure 2A**). Notably, the protein concentration was unchanged at all timepoints (**Figure 2B**). Despite the steady protein concentration, we have seen a drastic change in protein composition during wound healing. This was further highlighted in the volcano plot representations (**Figure 2C**).

**Figure 2.**
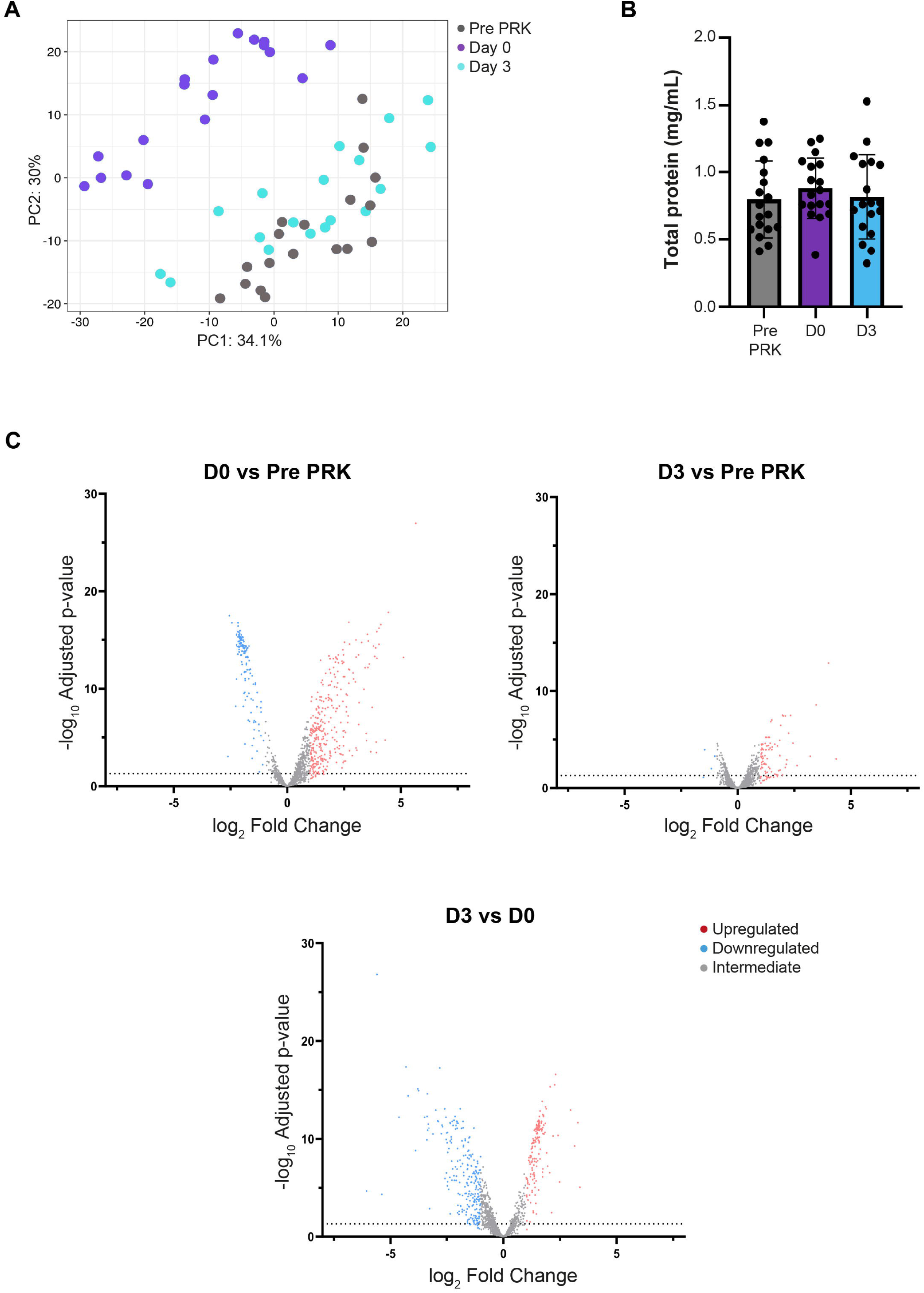
Drastic modulation of the tear film during corneal wound healing. **(A)** The PCA clustering of the analyzed samples demonstrates a strong correlation of the samples collected before (pre PRK) or 3 days (D3) after the PRK procedure. The samples taken right after the procedure (D0) were clustered away from the two other conditions. (B) The PRK procedure did not affect the protein concentration of the tear film. **(C)** All identified proteins are represented in the volcano plots and the limit of statistical significance is shown with the horizontal dotted line. Genes and proteins with a fold change lower than 0.5 are colored in blue. Genes and proteins with a fold change higher than 2 are colored in red and their number is indicated in the top right corner of each plot. The genes and proteins with an intermediate fold change are colored in grey.

Among the 2,025 identified proteins, 909 (44.9%) were significantly modulated during the healing period, constituting what we designate as the PRK response (**Table 2**). To examine the timing of this response, we stratified modulated proteins into six distinct categories: those upregulated or downregulated at D0, at D3, or across both time points.

**Table 2.**
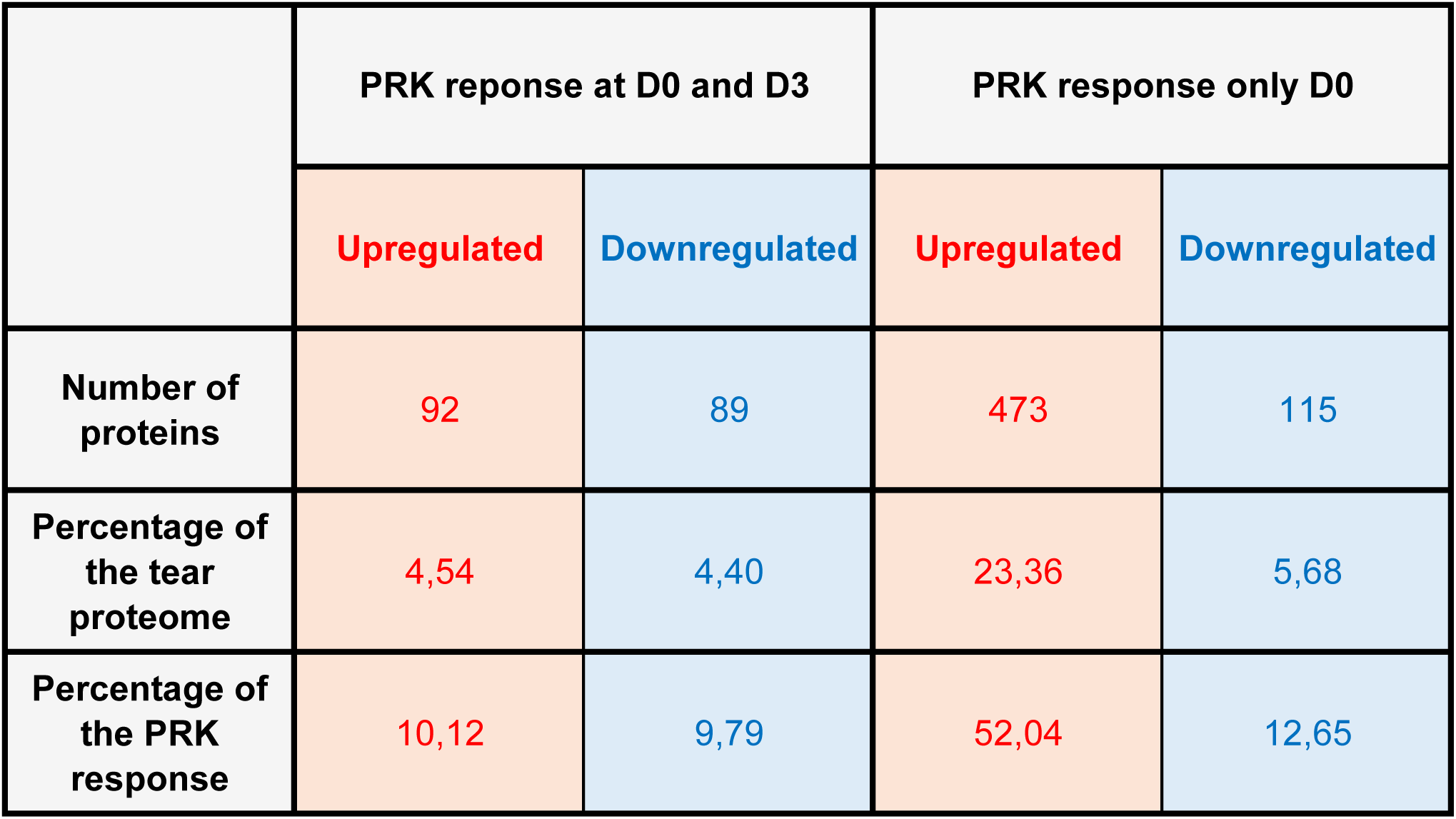
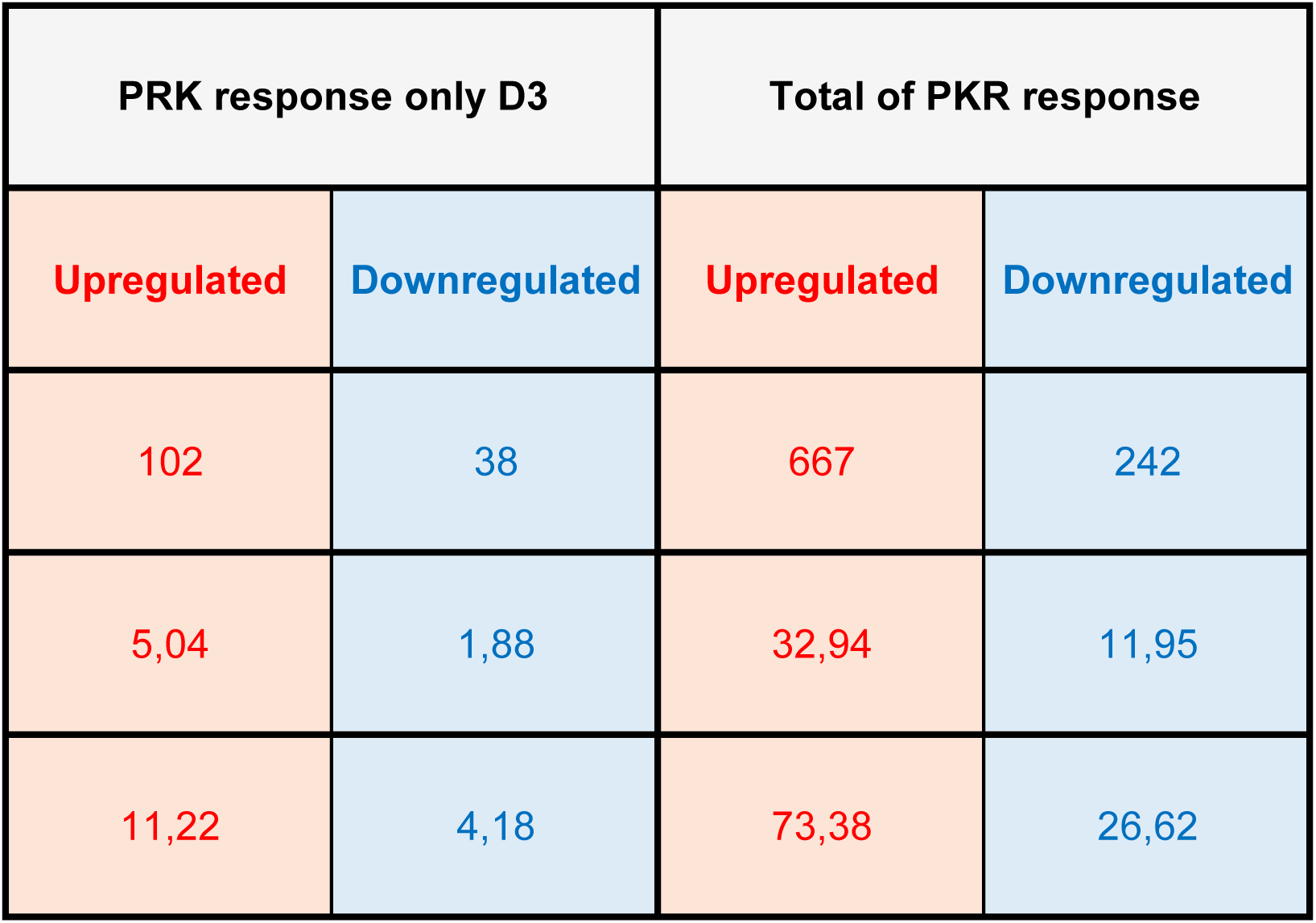
Overview of the tear fluid modifications upon PRK procedure. The molecular landscape modifications of the tear fluid are represented in terms of early, late and sustained modifications. These changes are compared to the whole tear film proteome, or PRK total response.

A major fraction of the PRK response—588 proteins (29.0% of the total proteome; 64.7% of the PRK response)—was transiently regulated within the first hour following surgery. An additional 181 proteins (8.9% of the total proteome; 19.9% of the PRK response) remained consistently modulated from D0 through D3, suggesting a sustained response integral to the wound healing cascade. The remaining 140 proteins (6.9% of the proteome; 15.4% of the PRK response) exhibited differential expression exclusively at D3, indicating delayed or secondary phase responses.

Collectively, these findings reveal that approximately 45% of the tear proteome is significantly altered after PRK. Because we have identified 909 proteins participating in the PRK response, we have deciphered the biological relevance of these important changes.

### The sustained PRK response

To delineate the molecular response to PRK, we have performed enrichment-analysis of the six protein categories (**Table 2**), according to three aspects: molecular pathways, GO biological process, and GO molecular function. First, we have analyzed the proteins which were modulated immediately after and 3 days after the PRK procedure. In this category, we have identified 181 proteins (19.91% of the total PRK response).

The main elements were reported in the **Table 3**. Throughout the corneal healing period, sustained activation was observed in molecular pathways predominantly associated with protein synthesis and cellular stress response, including “peptide chain elongation,” “eukaryotic translation termination and elongation,” and “nonsense-mediated decay.” Such consistent activation underscores a persistent demand for protein production to support ongoing tissue repair and regeneration (**Table 3, Table S1**). Similarly, the biological processes consistently upregulated included cytoplasmic translation, macromolecule biosynthesis, translation, and gene expression, reflecting continual cellular responses to sustain protein synthesis critical for cellular viability, repair, and structural integrity during tissue regeneration (**Table S2**). At the molecular function level, enduring activation of cadherin binding, GTPase activity, and ribonucleoside triphosphate phosphatase activity was prominent, indicating maintained cell adhesion, intercellular signaling, and nucleotide metabolism—functions critical for organized epithelial migration, cellular proliferation, and matrix organization essential in wound healing (**Table S3**).

**Table 3.**
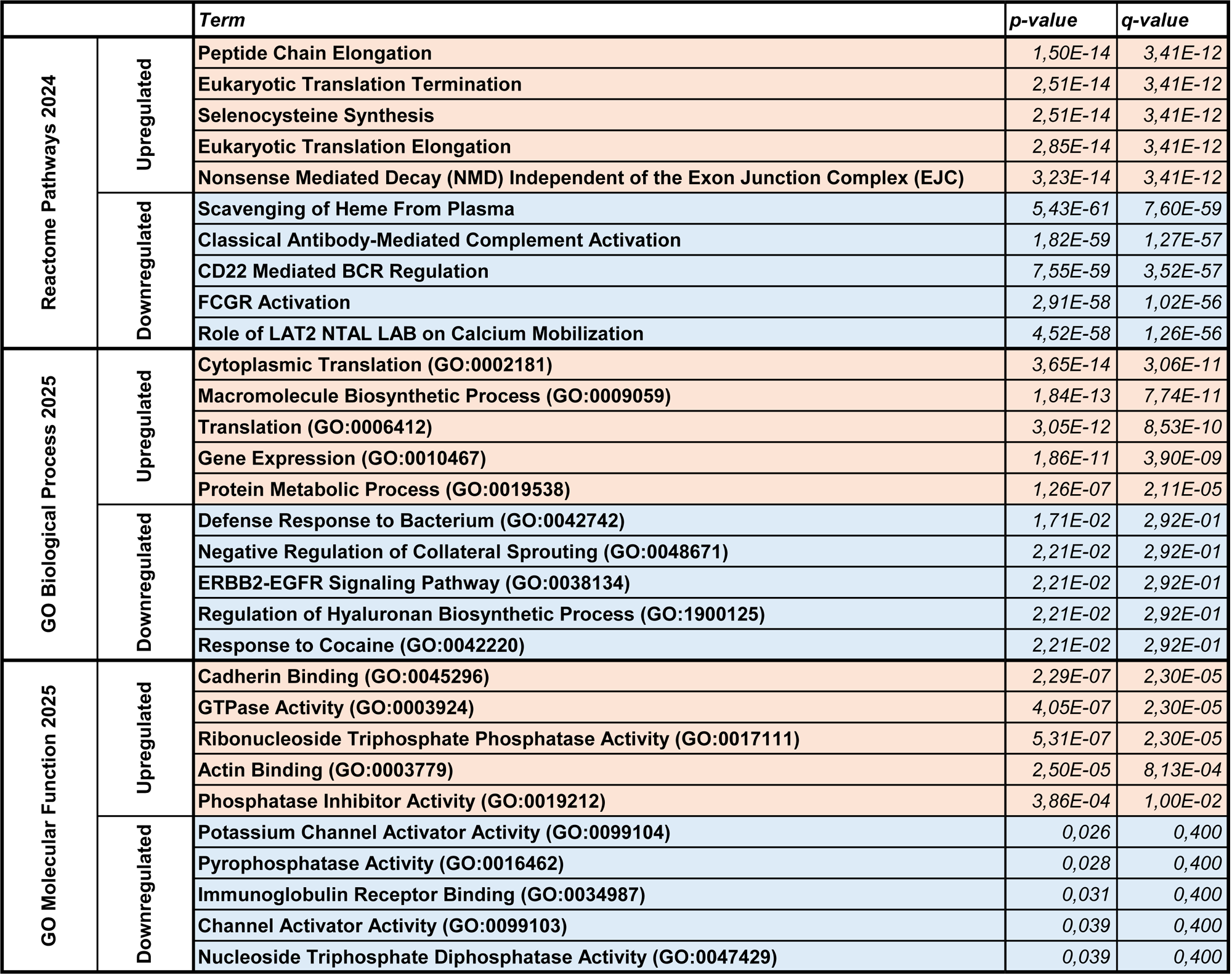
The sustained PRK response. The top 5 terms of each category (Reactome pathways 2024, GO Biological Process 2025, GO Molecular Function 2025) were displayed for the up and downregulated proteins.

Despite the relatively small set of upregulated proteins (92), notable functions included axon guidance (**Table S1**), epithelial development (**Table S2**), and IGF activity (**Table S3**)—all previously linked to corneal healing.

Concurrently, persistent suppression was observed in immune response-related pathways, notably involving scavenging of heme, classical antibody-mediated complement activation, FCGR activation, and LAT2 NTAL LAB-mediated calcium mobilization (**Table S4**). This prolonged downregulation indicates a deliberate, ongoing restraint of inflammatory responses essential to prevent prolonged or excessive inflammation, thus preserving corneal transparency and limiting fibrotic tissue development. Downregulated biological processes further reinforced immune modulation, specifically highlighting defense responses against bacteria, negative regulation of collateral neuronal sprouting, ERBB2-EGFR signaling, and hyaluronan biosynthetic processes (**Table S5**). Such sustained suppression might be critical in preventing aberrant nerve growth, excessive inflammation, and uncontrolled ECM production that could compromise healing outcomes. At the molecular function level, continuous downregulation of immunoglobulin receptor binding and potassium channel activator activities was identified, suggesting stringent control over immune cell signaling and ionic homeostasis necessary for cellular stability and controlled inflammatory responses throughout the healing period (**Table S6**). Finally, the suppression of negative regulation of axon regeneration was most likely necesseray for rapid corneal innervation regeneration.

### The core of the early PRK response

Then, we have analyzed the 588 proteins (64.69% of the total PRK response) which were modulated immediately after the PRK procedure (**Table 2**). The main elements were reported in the **Table 4**. Immediately after PRK, a robust activation of translation-associated pathways was observed (**Table S7**), notably involving “L13a-mediated translational silencing of ceruloplasmin,” “formation of free 40S subunits,” and “eukaryotic translation initiation”, among others. This translational activity was unveiled in the GO biological processes (**Table S8**) and GO molecular functions (**Table S9**), suggesting a prompt cellular effort to manage stress responses by controlling protein synthesis and turnover. Furthermore, biological processes predominantly feature cytoplasmic translation, macromolecule biosynthesis, and general protein metabolic activities, reflecting immediate priorities toward synthesizing proteins necessary for cell survival and repair initiation. Finally, cadherin binding and robust endopeptidase inhibitor activity indicate enhanced intercellular communication and protective responses against proteolysis, crucial for immediate tissue repair.

**Table 4.**
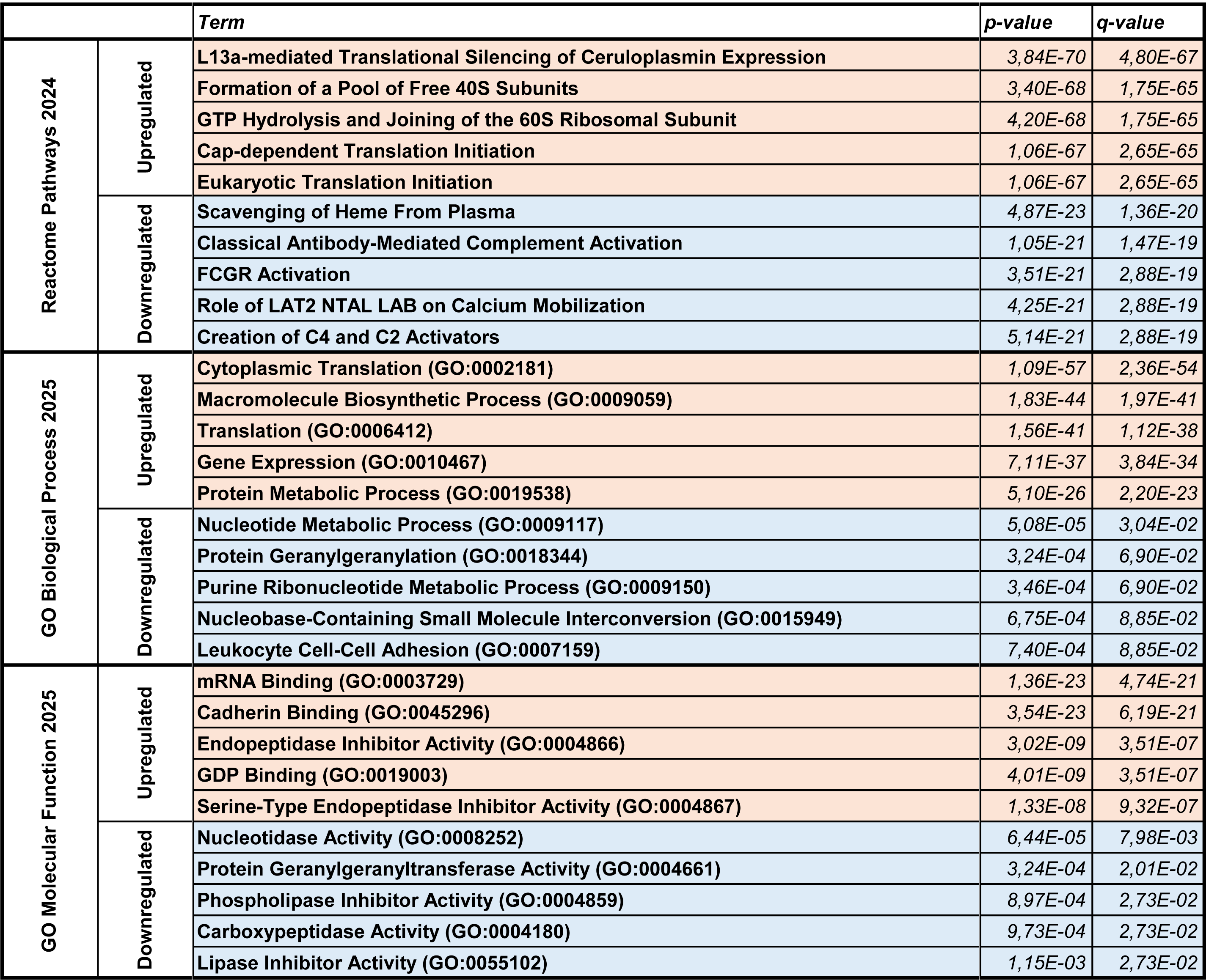
The early PRK response. The top 5 terms of each category (Reactome pathways 2024, GO Biological Process 2025, GO Molecular Function 2025) were displayed for the up and downregulated proteins.

Conversely, the analysis of the downregulated proteins has shown a significant suppression of immune-related molecular pathways, such as scavenging of heme from plasma, classical antibody-mediated complement activation, and FCGR activation, indicating a rapid modulation to limit inflammatory damage (**Table S10**). The suppression of LAT2 NTAL LAB calcium mobilization have underscored a deliberate inhibition of early inflammatory signaling, critical to minimizing initial cellular stress and maintaining corneal transparency. The GO biological processes have illustrated a significant downregulation in nucleotide metabolism, protein geranylgeranylation, and leukocyte cell-cell adhesion, reflecting a strategic reduction in metabolic demands and inflammation-related processes to mitigate potential excessive inflammatory responses and cellular stress (**Table S11**). At the molecular function level, nucleotidase activity and protein geranylgeranyltransferase activities were also notably downregulated, demonstrating a metabolic restraint that may protect cells by temporarily reducing metabolic burden and controlling cellular proliferation at this early injury phase (**Table S12**).

### The late PRK response

Finally, we have analyzed the 140 proteins (15.40% of the total PRK response) which were modulated only three days after the PRK procedure (**Table 2**). The main elements identified after the proteomic analysis were reported in the **Table 5**. This analysis at day 3 has revealed a significant upregulation in inflammatory and regenerative pathways. Notably, “neutrophil degranulation” and “post-translational protein phosphorylation” were highly active, signifying a controlled yet robust inflammatory response crucial for wound sterilization and initial tissue regeneration. Additionally, pathways regulating IGF transport and innate immunity further reflected a coordinated cellular proliferation and immune modulation (**Table S13**). The identified biological processes prominently included the positive regulation of cell adhesion, establishment and maintenance of epithelial cell polarity, and negative regulation of platelet aggregation (**Table S14**). These processes collectively facilitate precise epithelial migration and stable tissue formation, essential for optimal wound closure. The main molecular functions at day 3 emphasized an increased cadherin binding, endopeptidase inhibitor activity, and exopeptidase activity, underscoring mechanisms maintaining cellular cohesion, proteolytic regulation, and controlled matrix degradation necessary for effective epithelial resurfacing and stabilization (**Table S15**).

**Table 5.**
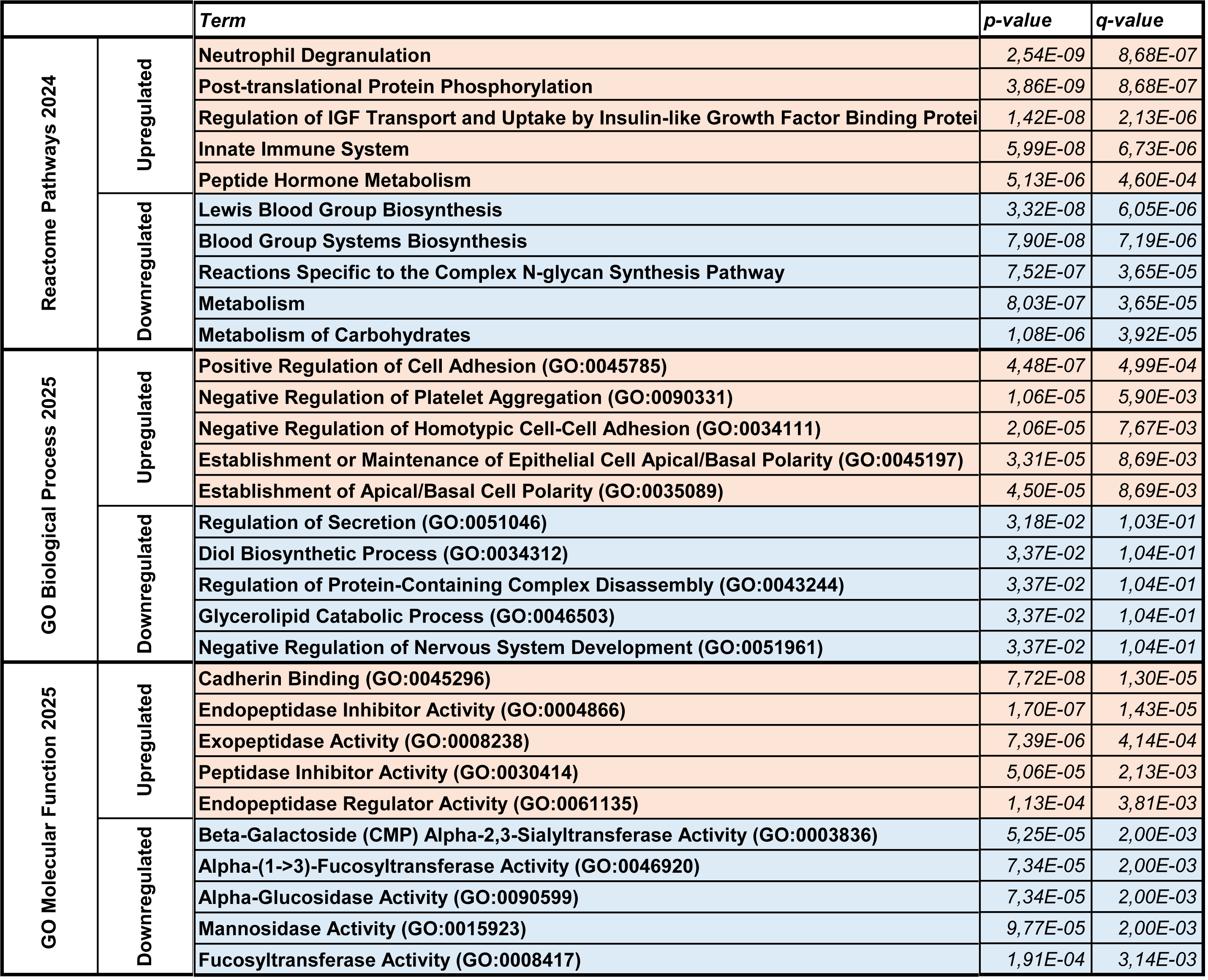
The late PRK response. The top 5 terms of each category (Reactome pathways 2024, GO Biological Process 2025, GO Molecular Function 2025) were displayed for the up and downregulated proteins.

Simultaneously, there was a notable suppression of molecular pathways related to glycosylation processes, including Lewis blood group biosynthesis and complex N-glycan synthesis, as well as general carbohydrate metabolism (**Table S16**). This downregulation reflects controlled glycoprotein synthesis, likely minimizing excessive ECM interactions to promote optimal cell migration and adhesion. The corresponding biological processes reinforced this theme, demonstrating suppressed protein glycosylation, oligosaccharide biosynthesis, and general carbohydrate metabolism (**Table S17**). These findings suggest that the corneal healing process at day 3 strategically minimizes unnecessary or aberrant glycosylation to facilitate effective epithelial repair and controlled ECM deposition. Finally, the molecular functions exhibited decreased activities of enzymes involved in carbohydrate processing and glycosylation, such as alpha-glucosidase, mannosidase, and fucosyltransferase (**Table S18**). Such downregulation points toward deliberate control over ECM structural composition and tissue remodeling.

## Discussion

Corneal abrasions and foreign bodies are common ocular injuries typically managed by primary care physicians. Although many corneal epithelial defects resolve spontaneously, a significant subset can progress to sight-threatening complications such as corneal ulcers or recurrent erosions. This underscores the critical need for a deeper mechanistic understanding of corneal wound healing to optimize clinical outcomes and prevent long-term visual impairment.

In this study, we have presented a non-invasive strategy to investigate corneal healing by profiling the tear film proteome—a dynamic and accessible window into the molecular events occurring at the ocular surface. By leveraging the PRK procedure in healthy individuals as a standardized and controlled model of epithelial injury, we created an opportunity to longitudinally examine the molecular landscape of corneal repair in a real-world, yet tightly regulated clinical setting.

Strikingly, PRK induced a profound and immediate remodeling of the tear film proteome, with over 900 proteins—representing nearly 45% of the detectable proteome—showing significant modulation. These changes occurred in a temporally dynamic manner, with the majority of molecular alterations observed immediately after surgery. Functional analysis of these differentially expressed proteins unveiled stage-specific biological processes associated with distinct phases of wound healing.

Immediately post-PRK, we observed a simultaneous upregulation of protein synthesis-related pathways and suppression of immune activation, reflecting a tightly orchestrated response aimed at stabilizing the corneal environment. This early phase prioritizes cellular repair, minimizes inflammatory damage, and conserves metabolic resources. Simultaneous modulation of adhesion molecules and inflammatory mediators emphasizes precise timing, facilitating wound closure and preventing excessive inflammation or scarring.

By day 3, the molecular profile shifted to support the resolution of inflammation and re-establishment of epithelial integrity. Upregulation of epithelial regeneration markers coincided with downregulation of glycosylation and carbohydrate metabolism, suggesting a strategic reprioritization toward tissue restoration. These complementary events are critical for optimal matrix remodeling and long-term visual recovery. Importantly, the continued activation of protein synthesis alongside suppression of inflammatory and extracellular matrix (ECM)-related pathways indicates a sustained reparative state. This phase is further characterized by enhanced regulation of cell adhesion and reestablishment of epithelial polarity—hallmarks of functional epithelial regeneration[5].

Collectively, our findings reveal a highly dynamic and temporally compartmentalized modulation of the ocular surface proteome following corneal injury. This study provides the first in-depth proteomic dissection of human tear film responses to PRK, delineating early, intermediate, and sustained phases of corneal wound healing. These insights offer a foundation for developing time-tailored therapeutic strategies that enhance wound healing while minimizing adverse outcomes such as fibrosis, delayed re-epithelialization, or vision loss.

Beyond the context of refractive surgery, our results have broader implications. The identification of tear-based biomarkers with temporal specificity offers a powerful tool to monitor healing status, stratify patients by risk, and guide personalized interventions in a range of ocular surface disorders—including persistent epithelial defects, corneal ulcers, and severe dry eye disease[19]. The clinical relevance of these findings is substantial: as refractive procedures grow in popularity, ensuring optimal healing trajectories is critical to maximizing safety, visual outcomes, and patient trust.

Finally, integrating tear film proteomics into clinical practice offers a transformative opportunity to monitor the healing process in a minimally invasive, repeatable manner. Persistent molecular signatures in the tear film could serve as valuable biomarkers for healing efficacy and therapeutic responsiveness [27,28]. Given the essential interplay of growth factors, inflammatory mediators, and ECM regulators in corneal regeneration, elucidating these molecular dynamics is key to unlocking new therapeutic avenues for vision preservation[5,10].

## Material and Methods

### Tear sampling

Tear samples were collected from each participant at three distinct time points: (1) prior to surgery (baseline), (2) approximately 30 minutes after corneal abrasion caused during the PRK procedure (acute post-injury phase), and (3) on postoperative day 3 (late healing phase).

Eligible participants were adults aged 18 years or older undergoing PRK for refractive indications. Patients with a history of ocular surgery, active ocular surface disease, systemic inflammatory or autoimmune disorders, or use of topical ocular medication (other than those required for surgery) were excluded. All participants provided written informed consent prior to inclusion. The study was approved by the relevant French institutional ethics committee and conducted in accordance with French regulations and the Declaration of Helsinki. The creation of a tear collection was approved by the French Ministry of Higher Education and Research (MESR) under number DC-2020-4151.

Tear fluid was collected using sterile Schirmer test strips following a standardized protocol. Bilateral Schirmer strips were carefully placed into the inferior conjunctival fornix of each eye without contacting the skin, ensuring aseptic technique throughout. Strips were left in place until at least 5 mm was wetted, typically taking seconds to minutes.

Tear-imbibed portions of the strips were transferred into pre-labelled sterile Eppendorf tubes and were immediately frozen at −20c°C and subsequently transported to the proteomics laboratory within 10 days for storage at −80c°C and further analysis.

### Proteomics analysis

The proteomic profile of tear samples was studied. The Schirmer strips were placed in a spin column (reference 69725, ThermoFisher Scientific) and incubated during 3 hours at 4°C with 150µL of 1% SDS 100mM ABC pH 8.5 solution. The spin columns were then transferred in LoBind® tubes (Eppendorf) and centrifuged at 13,000rpm during 2 minutes. The total protein concentration was quantified using the NanodropOne Microvolume UV-Vis Spectrophotometer (reference ND-ONE-W, ThermoFisher Scientific) at 280 nm. Tears proteomic preparation was carried out using the SP3 automated protocol on the LT Bravo Liquid handler (Agilent technologies) as described previously [29]. Briefly, 5µg of total protein from each sample were diluted in SDS to a final percentage of 1% and reduced with 5µL of 80mM DTT for 30 min at 60°C. The proteins were then alkylated with 5µL of 200mM IAA for 30 min at 30°C. Protein capture was performed by adding 5µL of Cytiva Sera-Mag at 100mg/mL (1:1 mix E3 and E7) and 35µL of acetonitrile. Beads were washed twice with 80% ethanol and once with acetonitrile. Digestion was performed by adding 250ng trypsin/lysC mix (Promega) in 35µL of 50mM ABC overnight at 37°C and 450 rpm, and was stopped with 10µL of 5% of formic acid. Peptide recovery was carried out using a magnetic rack prior to loading onto Evotips Pure™ according to manufacturer instructions. The peptide concentration was quantified using the NanodropOne Microvolume UV-Vis Spectrophotometer (reference ND-ONE-W, ThermoFisher Scientific) at 205 nm, a total of 400ng of peptides were desalted and peptide separation was performed using a EvoSep One liquid chromatography system (EvoSep) [30] with a PepSep C18 column, 15cm x 150µm, 1.5µm (Bruker Daltonics). Elution was carried out over 34-minutes gradient corresponding at 30 sample per day (SPD) using mobile phase A (0.1% FA in water; Biosolve) and mobile phase B (0.1% FA in acetontrile; Biosolve). Peptides were analyzed using a trapped ion mobility spectrometry quadrupole time-of-flight mass spectrometer (timsTOF HT, Bruker Daltonics) equipped with a nanoelectrospray ion source (Captive spray, Bruker Daltonics) in positive-ion mode. Data were acquired in a Data-Independent Acquisition (DIA) mode. MS1 and MS2 spectra were collected from 100 to 1700 m/z. Protein identification was then performed with DIA-NN software (version 1.9). The parameters used were the following: the digestion enzyme is trypsin, the number of missed cleavages was 1, the minimum peptide size was 7 amino acids, the precursor charge range was set from 1 to 4, the precursor m/z set from 300 to 1800 and a protein identification FDR was set at 1%. The Uniprot database of the human proteome was used as reference (Release_2024_03). Some modifications, induced by the sample preparation protocol, were studied: asparagine deamidation and methionine oxidation as variable modifications and cysteine carbamidomethylation as fixed modification. The maximum number of variable modifications was set to 2. LFQ intensities obtained from DIA-NN software (version 1.9) were processed using the DIA-Analyst platform (version 0.9.2). Proteins considered as contaminants and redundant were removed. The LFQ data for each protein were transformed by applying the log2(x) formula and grouped according to genotype (pre-op, post-op, J3). Missing values were not imputated. A cutoff of the adjusted p-value of 0.05 (t-statistic correction) along with a log2(fold change) of 1 has been applied to determine significantly regulated proteins in each pairwise comparison (paired test). Data were analysed using GraphPad Prism software (version 10.1.2, Prism, CA, USA).

### Availability of proteomics raw data

The raw data is available upon request and the datasets produced for the current study are publicly available. The mass spectrometry proteomics data is available in the PRIDE repository, via ProteomeXchange, with the identifier PXDxxxxxx.

## Functional and pathway enrichment analyses

We selected the proteins enriched or depleted at each timepoint based on a *p-value*<0.05, and a fold change >2 (enriched), or <0.5 (depleted). The resulting lists of proteins were analyzed using the Enrichr platforme (https://maayanlab.cloud/Enrichr/), followed by Appyter (https://appyters.maayanlab.cloud/) for enrichment analysis visualization. We gathered the analyses related to the Reactome Pathways 2024, GO Biological Pathways 2025, and GO Molecular Function 2025.

## Author contributions

Conceptualization: N.F., S.T.S., J.V., C.H., V.D., F.M.; Methodology: N.F., S.T.S., J.K., H.C., L.F., J.V., C.H., V.D., F.M.; Validation: J.V., C.H., V.D., F.M.; Formal analysis: N.F., S.T.S., J.K., H.C., L.F.; Data analysis: N.F., J.K., L.F., J.V., F.M.; Writing: N.F., J.V., C.H., V.D., F.M.; Supervision: V.D., F.M.; Project administration: V.D., F.M.; Funding acquisition: V.D., F.M.

## Acknowledgments

This research was supported by ATIP-Avenir program, Inserm, the CHU Montpellier (AOI FiLaPhyCo), the Région Occitanie, ANR (ANR-21-CE17-0061, TeFiCoPa), FRM (REP202110014140), Support for research: I-SITE 2024 - program of excellence of the University of Montpellier, CBS2 Doctoral School. Mass spectrometry experiments were carried out using the facilities of the Montpellier Proteomics Platform (PPM, BioCampus Montpellier), a member of the national Proteomics French Infrastructure (ProFI UAR 2048) supported by the French National Research Agency (ANR-24-INBS-0015, Investments for the future F2030).

## Supplemental Tables

**Table S1.** The Reactome Pathway analysis for the upregulated proteins identified at days 0 and 3.

Only the terms with a *p-value*<0.05 were displayed.

**Table S2.** The GO Biological Process analysis for the upregulated proteins identified at days 0 and 3.

Only the terms with a *p-value*<0.05 were displayed.

**Table S3.** The GO Molecular Function analysis for the upregulated proteins identified at days 0 and 3.

Only the terms with a *p-value*<0.05 were displayed.

**Table S4.** The Reactome Pathway analysis for the downregulated proteins identified at days 0 and 3.

Only the terms with a *p-value*<0.05 were displayed.

**Table S5.** The GO Biological Process analysis for the downregulated proteins identified at days 0 and 3.

Only the terms with a *p-value*<0.05 were displayed.

**Table S6.** The GO Molecular Function analysis for the downregulated proteins identified at days 0 and 3.

Only the terms with a *p-value*<0.05 were displayed.

**Table S7.** The Reactome Pathway analysis for the upregulated proteins identified at day 0.

Only the terms with a *p-value*<0.05 were displayed.

**Table S8.** The GO Biological Process analysis for the upregulated proteins identified at day 0.

Only the terms with a *p-value*<0.05 were displayed.

**Table S9.** The GO Molecular Function analysis for the upregulated proteins identified at day 0.

Only the terms with a *p-value*<0.05 were displayed.

**Table S10.** The Reactome Pathway analysis for the downregulated proteins identified at day 0.

Only the terms with a *p-value*<0.05 were displayed.

**Table S11.** The GO Biological Process analysis for the downregulated proteins identified at day 0. Only the terms with a *p-value*<0.05 were displayed.

**Table S12.** The GO Molecular Function analysis for the downregulated proteins identified at day 0.

Only the terms with a *p-value*<0.05 were displayed.

**Table S13.** The Reactome Pathway analysis for the upregulated proteins identified at day 3.

Only the terms with a *p-value*<0.05 were displayed.

**Table S14.** The GO Biological Process analysis for the upregulated proteins identified at day 3.

Only the terms with a *p-value*<0.05 were displayed.

**Table S15.** The GO Molecular Function analysis for the upregulated proteins identified at day 3.

Only the terms with a *p-value*<0.05 were displayed.

**Table S16.** The Reactome Pathway analysis for the downregulated proteins identified at day 3.

Only the terms with a *p-value*<0.05 were displayed.

**Table S17.** The GO Biological Process analysis for the downregulated proteins identified at day 3.

Only the terms with a *p-value*<0.05 were displayed.

**Table S18.** The GO Molecular Function analysis for the downregulated proteins identified at day 3.

Only the terms with a *p-value*<0.05 were displayed.

## References

[1] DelMonte DW, Kim T. Anatomy and physiology of the cornea. J Cataract Refract Surg 2011;37:588–98. 10.1016/j.jcrs.2010.12.037.

[2] Güneş İB, Öztürk H, Özen B. The association of pre-photorefractive keratectomy Schirmer-1 test value with postoperative corneal epithelial thickness, ocular surface discomfort, and visual acuity. Arq Bras Oftalmol 2024;87:e20230049. 10.5935/0004-2749.2023-0049.

[3] Barrientez B, Nicholas SE, Whelchel A, Sharif R, Hjortdal J, Karamichos D. Corneal injury: Clinical and molecular aspects. Exp Eye Res 2019;186:107709. 10.1016/j.exer.2019.107709.

[4] Wilson SE. Corneal wound healing. Exp Eye Res 2020;197:108089. 10.1016/j.exer.2020.108089.

[5] Ljubimov AV, Saghizadeh M. Progress in corneal wound healing. Prog Retin Eye Res 2015;49:17–45. 10.1016/j.preteyeres.2015.07.002.

[6] Tomás-Juan J, Murueta-Goyena Larrañaga A, Hanneken L. Corneal Regeneration After Photorefractive Keratectomy: A Review. J Optom 2015;8:149–69. 10.1016/j.optom.2014.09.001.

[7] Xu K, Yu F-SX. Impaired epithelial wound healing and EGFR signaling pathways in the corneas of diabetic rats. Invest Ophthalmol Vis Sci 2011;52:3301–8. 10.1167/iovs.10-5670.

[8] Chung B, Lee H, Choi BJ, Seo KR, Kim EK, Kim DY, et al. Clinical Outcomes of an Optimized Prolate Ablation Procedure for Correcting Residual Refractive Errors Following Laser Surgery. Korean J Ophthalmol KJO 2017;31:16–24. 10.3341/kjo.2017.31.1.16.

[9] Wilson DH, Rissin DM, Kan CW, Fournier DR, Piech T, Campbell TG, et al. The Simoa HD-1 Analyzer: A Novel Fully Automated Digital Immunoassay Analyzer with Single-Molecule Sensitivity and Multiplexing. J Lab Autom 2016;21:533–47. 10.1177/2211068215589580.

[10] Yan C, Jin L, Zhang Q, Liu X, Yu T, Zhao F, et al. Management of delayed corneal epithelial healing after refractive surgery: five case reports. Front Med 2025;12:1517403. 10.3389/fmed.2025.1517403.

[11] Maraghechi G, Ojaghi H, Amani F, Najafi A. A comparative study of Pentacam indices in various types and severities of refractive error in candidates for photorefractive keratectomy (PRK) surgery. J Med Life 2022;15:810–8. 10.25122/jml-2021-0027.

[12] Abing AA, Oh A, Ong LF, Marvasti AH, Tran DB, Lee JK. Surgical options and clinical outcomes for high myopia. Curr Opin Ophthalmol 2024;35:284–91. 10.1097/ICU.0000000000001053.

[13] Saad A, Saad A, Frings A. Refractive results of photorefractive keratectomy comparing trans-PRK and PTK-PRK for correction of myopia and myopic astigmatism. Int Ophthalmol 2024;44:111. 10.1007/s10792-024-02999-w.

[14] Wen D, McAlinden C, Flitcroft I, Tu R, Wang Q, Alió J, et al. Postoperative Efficacy, Predictability, Safety, and Visual Quality of Laser Corneal Refractive Surgery: A Network Meta-analysis. Am J Ophthalmol 2017;178:65–78. 10.1016/j.ajo.2017.03.013.

[15] Guedes J, Vilares-Morgado R, Brazuna R, Neto AC, Mora-Paez DJ, Salomão MQ, et al. Pressure-Induced Stromal Keratopathy after Surface Ablation Surgery. Case Rep Ophthalmol 2024;15:532–41. 10.1159/000539701.

[16] Kremer I, Ehrenberg M, Levinger S. Delayed epithelial healing following photorefractive keratectomy with mitomycin C treatment. Acta Ophthalmol (Copenh) 2012;90:271–6. 10.1111/j.1755-3768.2010.01894.x.

[17] Arranz-Marquez E, Katsanos A, Kozobolis VP, Konstas AGP, Teus MA. A Critical Overview of the Biological Effects of Mitomycin C Application on the Cornea Following Refractive Surgery. Adv Ther 2019;36:786–97. 10.1007/s12325-019-00905-w.

[18] Wilson SE, Marino GK, Medeiros CS, Santhiago MR. Phototherapeutic Keratectomy: Science and Art. J Refract Surg Thorofare NJ 1995 2017;33:203–10. 10.3928/1081597X-20161123-01.

[19] Wilson SE, Medeiros CS, Santhiago MR. Pathophysiology of Corneal Scarring in Persistent Epithelial Defects After PRK and Other Corneal Injuries. J Refract Surg Thorofare NJ 1995 2018;34:59–64. 10.3928/1081597X-20171128-01.

[20] Klenkler B, Sheardown H, Jones L. Growth Factors in the Tear Film: Role in Tissue Maintenance, Wound Healing, and Ocular Pathology. Ocul Surf 2007;5:228–39. 10.1016/S1542-0124(12)70613-4.

[21] Wilson SE, Mohan RR, Mohan RR, Ambrósio R, Hong J, Lee J. The corneal wound healing response: cytokine-mediated interaction of the epithelium, stroma, and inflammatory cells. Prog Retin Eye Res 2001;20:625–37. 10.1016/s1350-9462(01)00008-8.

[22] Labetoulle M, Baudouin C, Calonge M, Merayo-Lloves J, Boboridis KG, Akova YA, et al. Role of corneal nerves in ocular surface homeostasis and disease. Acta Ophthalmol (Copenh) 2019;97:137–45. 10.1111/aos.13844.

[23] Winiarczyk M, Biela K, Michalak K, Winiarczyk D, Mackiewicz J. Changes in Tear Proteomic Profile in Ocular Diseases. Int J Environ Res Public Health 2022;19:13341. 10.3390/ijerph192013341.

[24] Fournié PR, Gordon GM, Dawson DG, Malecaze FJ, Edelhauser HF, Fini ME. Correlation between epithelial ingrowth and basement membrane remodeling in human corneas after laser-assisted in situ keratomileusis. Arch Ophthalmol Chic Ill 1960 2010;128:426–36. 10.1001/archophthalmol.2010.23.

[25] Nishtala K, Pahuja N, Shetty R, Nuijts RMMA, Ghosh A. Tear biomarkers for keratoconus. Eye Vis Lond Engl 2016;3:19. 10.1186/s40662-016-0051-9.

[26] Zhong Y, Fang X, Wang X, Lin Y-A, Wu H, Li C. Effects of Sodium Hyaluronate Eye Drops With or Without Preservatives on Ocular Surface Bacterial Microbiota. Front Med 2022;9:793565. 10.3389/fmed.2022.793565.

[27] Jacob JT, Ham B. Compositional profiling and biomarker identification of the tear film. Ocul Surf 2008;6:175–85. 10.1016/s1542-0124(12)70178-7.

[28] Ambaw YA, Timbadia DP, Raida M, Torta F, Wenk MR, Tong L. Profile of tear lipid mediator as a biomarker of inflammation for meibomian gland dysfunction and ocular surface diseases: Standard operating procedures. Ocul Surf 2022;26:318–27. 10.1016/j.jtos.2020.09.008.

[29] Müller T, Kalxdorf M, Longuespée R, Kazdal DN, Stenzinger A, Krijgsveld J. Automated sample preparation with SP3 for low-input clinical proteomics. Mol Syst Biol 2020;16:e9111. 10.15252/msb.20199111.

[30] Bache N, Geyer PE, Bekker-Jensen DB, Hoerning O, Falkenby L, Treit PV, et al. A Novel LC System Embeds Analytes in Pre-formed Gradients for Rapid, Ultra-robust Proteomics. Mol Cell Proteomics MCP 2018;17:2284–96. 10.1074/mcp.TIR118.000853.

