## Supplemental Table S1 for "Temporal analysis of tear fluid proteome reveals critical corneal repair events after photorefractive surgery"

| term |
| --- |
| Peptide Chain Elongation |
| Eukaryotic Translation Termination |
| Selenocysteine Synthesis |
| Eukaryotic Translation Elongation |
| Nonsense Mediated Decay (NMD) Independent of the Exon Junction Complex (EJC) |
| Viral mRNA Translation |
| Response of EIF2AK4 (GCN2) to Amino Acid Deficiency |
| Formation of a Pool of Free 40S Subunits |
| L13a-mediated Translational Silencing of Ceruloplasmin Expression |
| GTP Hydrolysis and Joining of the 60S Ribosomal Subunit |
| SRP-dependent Cotranslational Protein Targeting to Membrane |
| Nonsense Mediated Decay (NMD) Enhanced by the Exon Junction Complex (EJC) |
| Nonsense-Mediated Decay (NMD) |
| Selenoamino Acid Metabolism |
| Eukaryotic Translation Initiation |
| Cap-dependent Translation Initiation |
| Regulation of Expression of SLITs and ROBOs |
| Influenza Viral RNA Transcription and Replication |
| Axon Guidance |
| Nervous System Development |
| Cellular Response to Starvation |
| Signaling by ROBO Receptors |
| Influenza Infection |
| Major Pathway of rRNA Processing in the Nucleolus and Cytosol |
| rRNA Processing in the Nucleus and Cytosol |
| Regulation of Complement Cascade |
| Complement Cascade |
| rRNA Processing |
| Metabolism of Amino Acids and Derivatives |
| Translation |
| Metabolism of RNA |
| Innate Immune System |
| Immune System |
| Platelet Activation Signaling and Aggregation |
| Infectious Disease |
| Cellular Responses to Stimuli |
| Disease |
| Developmental Biology |
| Cellular Responses to Stress |
| Activation of C3 and C5 |
| Opioid Signalling |
| Viral Infection Pathways |
| Hemostasis |
| Metabolism of Proteins |
| Signal Amplification |

| |
|---|
| Platelet Degranulation |
| Regulation of Insulin Secretion |
| Response to Elevated Platelet Cytosolic Ca2+ |
| Beta-catenin Independent WNT Signaling |
| Recycling Pathway of L1 |
| Protein Hydroxylation |
| G Beta Gamma Signalling Through PLC Beta |
| Metabolism |
| Presynaptic Function of Kainate Receptors |
| ADP Signalling Through P2Y Purinoceptor 12 |
| Integration of Energy Metabolism |
| Intrinsic Pathway of Fibrin Clot Formation |
| G-protein Activation |
| Thromboxane Signalling Through TP Receptor |
| ADP Signalling Through P2Y Purinoceptor 1 |
| Ca2+ Pathway |
| L1CAM Interactions |
| Activation of Kainate Receptors Upon Glutamate Binding |
| Neutrophil Degranulation |
| Neurotransmitter Receptors and Postsynaptic Signal Transmission |
| Thrombin Signalling Through Proteinase Activated Receptors (PARs) |
| G-protein Beta Gamma Signalling |
| Fatty Acids Bound to GPR40 (FFAR1) Regulate Insulin Secretion |
| Terminal Pathway of Complement |
| Cooperation of PDCL (PhLP1) and TRiC CCT in G-protein Beta Folding |
| Formation of Fibrin Clot (Clotting Cascade) |
| Chaperonin-mediated Protein Folding |
| Acetylcholine Regulates Insulin Secretion |
| HuR (ELAVL1) Binds and Stabilizes mRNA |
| MET Receptor Recycling |
| Glucagon-like Peptide-1 (GLP1) Regulates Insulin Secretion |
| ADORA2B Mediated Anti-Inflammatory Cytokines Production |
| Activation of GABAB Receptors |
| GABA B Receptor Activation |
| Protein Folding |
| Free Fatty Acids Regulate Insulin Secretion |
| Interferon Gamma Signaling |
| GPER1 Signaling |
| Initial Triggering of Complement |
| G Alpha (Z) Signalling Events |
| Transmission Across Chemical Synapses |
| Interferon Signaling |
| G-protein Mediated Events |
| Sensory Processing of Sound by Outer Hair Cells of the Cochlea |
| Signaling by WNT |
| Defects of Contact Activation System (CAS) and Kallikrein Kinin System (KKS) |

| |
|---|
| GABA Receptor Activation |
| MHC Class II Antigen Presentation |
| Gamma Carboxylation Hypusinylation Hydroxylation and Arylsulfatase Activation |
| Neuronal System |
| G Beta Gamma Signalling Through BTK |
| G Alpha (I) Signalling Events |
| Prostacyclin Signalling Through Prostacyclin Receptor |
| Diseases of Hemostasis |
| Plasma Lipoprotein Assembly |
| G Alpha (Q) Signalling Events |
| G Beta Gamma Signalling Through CDC42 |
| Sensory Processing of Sound by Inner Hair Cells of the Cochlea |
| High Laminar Flow Shear Stress Activates Signaling by PIEZO1 and PECAM1 CDH5 KDR in Endothelial [illegible] |
| Translocation of SLC2A4 (GLUT4) to the Plasma Membrane |
| Plasma Lipoprotein Assembly Remodeling and Clearance |
| Extra-nuclear Estrogen Signaling |
| Regulation of mRNA Stability by Proteins That Bind AU-rich Elements |
| Sensory Processing of Sound |
| Parasitic Infection Pathways |
| Leishmania Infection |
| Signaling by MET |
| G Beta Gamma Signalling Through PI3Kgamma |
| Syndecan Interactions |
| Adrenaline noradrenaline Inhibits Insulin Secretion |
| Vesicle-mediated Transport |
| Activation of G Protein Gated Potassium Channels |
| G Protein Gated Potassium Channels |
| Inhibition of Voltage Gated Ca2+ Channels via Gbeta Gamma Subunits |
| Cellular Responses to Mechanical Stimuli |
| Response of Endothelial Cells to Shear Stress |
| Signaling by ALK Fusions and Activated Point Mutants |
| Signaling by ALK in Cancer |
| Cytokine Signaling in Immune System |
| Glucagon Signaling in Metabolic Regulation |
| Glucagon-type Ligand Receptors |
| Inwardly Rectifying K+ Channels |
| Plasma Lipoprotein Clearance |
| SARS-CoV-1 Modulates Host Translation Machinery |
| Vasopressin Regulates Renal Water Homeostasis via Aquaporins |
| TBC RABGAPs |
| Binding and Uptake of Ligands by Scavenger Receptors |
| Regulation of IGF Transport and Uptake by Insulin-like Growth Factor Binding Proteins (IGFBPs) |
| PLC Beta Mediated Events |
| Alternative Complement Activation |
| Extrinsic Pathway of Fibrin Clot Formation |
| VLDL Assembly |

| |
|---|
| Aquaporin-mediated Transport |
| Formation of the Ternary Complex and Subsequently the 43S Complex |
| SARS-CoV-2 Modulates Host Translation Machinery |
| GPCR Downstream Signalling |
| Defective F9 Activation |
| VLDL Clearance |
| Signal Transduction |
| Clathrin-mediated Endocytosis |
| Non-integrin membrane-ECM Interactions |
| Translation Initiation Complex Formation |
| Ribosomal Scanning and Start Codon Recognition |
| ARMS-mediated Activation |
| Defective Factor VIII Causes Hemophilia A |
| mRNA Activation Upon Binding of the Cap-Binding Complex and eIFs and Subsequent Binding to 43S |
| Signaling by Nuclear Receptors |
| The Role of GTSE1 in G2 M Progression After G2 Checkpoint |
| Nef Mediated CD8 Down-regulation |
| Noncanonical Activation of NOTCH3 |
| Regulation of Cytoskeletal Remodeling and Cell Spreading by IPP Complex Components |
| HDL Assembly |
| Insulin-like Growth Factor-2 mRNA Binding Proteins (IGF2BPs IMPs VICKZs) Bind RNA |
| G Alpha (S) Signalling Events |
| Anti-inflammatory Response Favouring Leishmania Parasite Infection |
| Leishmania Parasite Growth and Survival |
| Downstream Signaling Events of B Cell Receptor (BCR) |
| Gamma-carboxylated Protein Precursors Transport From the Endoplasmic Reticulum to Golgi Appara |
| Serine Biosynthesis |
| Defective Factor IX Causes Hemophilia B |
| Calcineurin Activates NFAT |
| Signaling by GPCR |
| ATF6 (ATF6-alpha) Activates Chaperone Genes |
| Nef Mediated CD4 Down-regulation |
| Removal of Aminoterminal Propeptides From Gamma-Carboxylated Proteins |
| Gamma-carboxylation of Protein Precursors |
| MET Activates RAP1 and RAC1 |
| Gamma-carboxylation Transport and Amino-Terminal Cleavage of Proteins |
| Purine Ribonucleoside Monophosphate Biosynthesis |
| CLEC7A (Dectin-1) Induces NFAT Activation |
| Extracellular Matrix Organization |

| p-value | q-value |
|---|---|
| 1.499578703186952e-14 | 3.413120489294446e-12 |
| 2.513339124712494e-14 | 3.413120489294446e-12 |
| 2.513339124712494e-14 | 3.413120489294446e-12 |
| 2.8494291313650456e-14 | 3.413120489294446e-12 |
| 3.2260118046261306e-14 | 3.413120489294446e-12 |
| 4.118473989002713e-14 | 3.631121233637392e-12 |
| 6.610343226332369e-14 | 4.371089458412279e-12 |
| 6.610343226332369e-14 | 4.371089458412279e-12 |
| 1.9907276411561658e-13 | 1.1691059594598851e-11 |
| 2.210030169111314e-13 | 1.1691059594598851e-11 |
| 2.715919881116268e-13 | 1.2234223173512893e-11 |
| 3.0065198725078940e-13 | 1.2234223173512893e-11 |
| 3.0065198725078940e-13 | 1.2234223173512893e-11 |
| 3.674254273574463e-13 | 1.3883432219434934e-11 |
| 4.4743907429176157e-13 | 1.4793454393771367e-11 |
| 4.4743907429176157e-13 | 1.4793454393771367e-11 |
| 5.974696239382616e-13 | 1.8591848886078846e-11 |
| 4.062192731998468e-12 | 1.1938333084595498e-10 |
| 4.355629123675215e-12 | 1.212698845486415e-10 |
| 9.834580216866983e-12 | 2.566953213973174e-10 |
| 1.0190173439212979e-11 | 2.566953213973174e-10 |
| 1.3990467973538139e-11 | 3.364071617273489e-10 |
| 1.8065607548866845e-11 | 4.155089736239374e-10 |
| 6.702701519185759e-11 | 1.4773871265205279e-09 |
| 1.2210808950462606e-10 | 2.5838071739178876e-09 |
| 3.7361896234927444e-10 | 7.6017088877987e-09 |
| 8.419687748420806e-10 | 1.649635118116521e-08 |
| 9.121741465937381e-10 | 1.7233575841003122e-08 |
| 9.6027538221639e-10 | 1.751674748939553e-08 |
| 1.2592370409978937e-09 | 2.220454648959619e-08 |
| 1.428364173117028e-09 | 2.3970732308129864e-08 |
| 1.4500253948207101e-09 | 2.3970732308129864e-08 |
| 1.3370215613830557e-08 | 2.1432860787019288e-07 |
| 3.3302604313857514e-08 | 5.18149343589136e-07 |
| 8.473641683289278e-08 | 1.2807304144171509e-06 |
| 9.83917634019836e-08 | 1.4458123011013702e-06 |
| 6.454824664881578e-07 | 9.228654723573932e-06 |
| 9.438516226158704e-07 | 1.313940811483672e-05 |
| 2.861158231443968e-06 | 3.8809043703432285e-05 |
| 3.2529312982147432e-06 | 4.3020016418889975e-05 |
| 3.6664514356480358e-06 | 4.730616608433685e-05 |
| 3.974510701257826e-06 | 5.005990859441405e-05 |
| 4.1137770073398665e-06 | 5.0609024113553245e-05 |
| 4.443006886226233e-06 | 5.341706006394721e-05 |
| 1.548917558984961e-05 | 0.0001820838641562321 |

| | |
|---|---|
| 2.777147172878048e-05 | 0.0003193719248809755 |
| 2.989505287122671e-05 | 0.0003364783610399772 |
| 3.4469543485755875e-05 | 0.0003798830938326012 |
| 3.595291543295465e-05 | 0.00038814474008230635 |
| 6.987324687650974e-05 | 0.0007392589519534731 |
| 0.00010145759537430765 | 0.001032135922173245 |
| 0.00010145759537430765 | 0.001032135922173245 |
| 0.00010426678254255495 | 0.0010407005276417277 |
| 0.00011797347851055985 | 0.001155703150594188 |
| 0.00013614657811150416 | 0.0013094825421997402 |
| 0.00014164253171419024 | 0.001338016058514404 |
| 0.00015604795273537798 | 0.0014482345087195607 |
| 0.0001777476099679932 | 0.0015937031470011594 |
| 0.0001777476099679932 | 0.0015937031470011594 |
| 0.00020131451592755997 | 0.0017599358091326014 |
| 0.00020294155833098048 | 0.0017599358091326014 |
| 0.00023190986738710773 | 0.001978714836254516 |
| 0.00034949988210486357 | 0.0029279045089775713 |
| 0.0003542266324660956 | 0.0029279045089775713 |
| 0.0003600734735136159 | 0.002930444115210812 |
| 0.0004241446308301244 | 0.0033488434284945643 |
| 0.0004241446308301244 | 0.0033488434284945643 |
| 0.0005755910882831092 | 0.004412865010170504 |
| 0.0005755910882831092 | 0.004412865010170504 |
| 0.0007071459809171628 | 0.005344003198645416 |
| 0.0007635347680054731 | 0.005688871722181623 |
| 0.0008256107047770111 | 0.006065945317042206 |
| 0.0009195296371531131 | 0.0064857490407199585 |
| 0.0009195296371531131 | 0.0064857490407199585 |
| 0.0009195296371531131 | 0.0064857490407199585 |
| 0.0009496127719297549 | 0.006609804688826847 |
| 0.001017451521498842 | 0.006813061454087183 |
| 0.001017451521498842 | 0.006813061454087183 |
| 0.001017451521498842 | 0.006813061454087183 |
| 0.0010481312895977365 | 0.006930768152465032 |
| 0.0011205081876088584 | 0.007295184270248407 |
| 0.0011308225144808495 | 0.007295184270248407 |
| 0.0011621548884364816 | 0.007406987180516853 |
| 0.0013574000601219033 | 0.008548388473862939 |
| 0.0014025124056790167 | 0.008728577207108233 |
| 0.0015194992010309533 | 0.009346686945876445 |
| 0.0018606228064929507 | 0.011313442122238747 |
| 0.0019717371905147415 | 0.011852829247526116 |
| 0.002078597445497338 | 0.012354809535596535 |
| 0.0021061199077987577 | 0.01237930479139492 |
| 0.0024084361889752624 | 0.0140006894941529 |

| | |
|---|---|
| 0.002666757887047368 | 0.015333857850522365 |
| 0.002736844387612744 | 0.015567641731689695 |
| 0.00279543322391944 | 0.0157317465473764 |
| 0.0028795807002267946 | 0.016034717793894468 |
| 0.0030524520685432928 | 0.016820282752702104 |
| 0.003392431208613783 | 0.01790870254792713 |
| 0.0034013861191127576 | 0.01790870254792713 |
| 0.0034013861191127576 | 0.01790870254792713 |
| 0.0034013861191127576 | 0.01790870254792713 |
| 0.003419241885332023 | 0.01790870254792713 |
| 0.0037680465442261175 | 0.019542123744074668 |
| 0.00396340569561867 | 0.020160015509444965 |
| 0.00396340569561867 | 0.020160015509444965 |
| 0.004295106750729711 | 0.021639156867962066 |
| 0.0050078725723263166 | 0.0247585475772021 |
| 0.0050078725723263166 | 0.02475854757720207 |
| 0.005196512221021259 | 0.025453286712224 5 |
| 0.005389384843798236 | 0.026149551446044993 |
| 0.005486956919680519 | 0.026149551446044993 |
| 0.005486956919680519 | 0.026149551446044993 |
| 0.005787931380304514 | 0.027337640180188284 |
| 0.005861443927181807 | 0.02743985696884227 |
| 0.006817134117284325 | 0.03163389428108253 |
| 0.00731969302231779 | 0.033670587902661836 |
| 0.007719499133596217 | 0.034845171244631525 |
| 0.007838516782818812 | 0.034845171244631525 |
| 0.007838516782818812 | 0.034845171244631525 |
| 0.007838516782818812 | 0.034845171244631525 |
| 0.009070200863679196 | 0.03900923786086419 |
| 0.009070200863679196 | 0.03900923786086419 |
| 0.009070200863679196 | 0.03900923786086419 |
| 0.009070200863679196 | 0.03900923786086419 |
| 0.009379150211011182 | 0.04001266501310416 |
| 0.010073338153561873 | 0.042292030819319294 |
| 0.010073338153561873 | 0.042292030819319294 |
| 0.011284308834115625 | 0.04700314467123752 |
| 0.012556035536271243 | 0.051891740614745996 |
| 0.015277004383447555 | 0.06264756061119192 |
| 0.016723910895674 | 0.06805345279855034 |
| 0.018226901785548283 | 0.0736032904164507 |
| 0.01876606947767571 | 0.07520644510371555 |
| 0.01958382008519542 | 0.0778935400381081 |
| 0.021396640282656202 | 0.08446882619048605 |
| 0.022791467529210928 | 0.08800500965658818 |
| 0.022791467529210928 | 0.08800500965658818 |
| 0.022791467529210928 | 0.08800500965658818 |

| | |
|---|---|
| 0.0239128890269455 | 0.0916660746032911 |
| 0.024777381825604797 | 0.0942966545737046 |
| 0.0265441835896253 | 0.10011871452741951 |
| 0.02668570651865057 | 0.10011871452741951 |
| 0.027287734343433324 | 0.1009455347389946 |
| 0.027287734343433324 | 0.1009455347389946 |
| 0.027488493918926157 | 0.10098203668827734 |
| 0.029347561301612073 | 0.10706799950726059 |
| 0.030225389407192507 | 0.1095152807972934 |
| 0.03117568899624756 | 0.11143202350685782 |
| 0.03117568899624756 | 0.11143202350685782 |
| 0.03176353861225931 | 0.1120194128392345 |
| 0.03176353861225931 | 0.1120194128392345 |
| 0.03213773350942613 | 0.11258848361911537 |
| 0.03605651862165359 | 0.12126478639776911 |
| 0.03610086196437731 | 0.12126478639776911 |
| 0.036218972118804385 | 0.12126478639776911 |
| 0.036218972118804385 | 0.12126478639776911 |
| 0.036218972118804385 | 0.12126478639776911 |
| 0.036218972118804385 | 0.12126478639776911 |
| 0.036218972118804385 | 0.12126478639776911 |
| 0.03646414745332675 | 0.1213178239170431 |
| 0.037045161097470636 | 0.12171981503454637 |
| 0.037045161097470636 | 0.12171981503454637 |
| 0.03919046306536466 | 0.12797379605912287 |
| 0.040654126315732 | 0.12955441458447126 |
| 0.040654126315732 | 0.12955441458447126 |
| 0.040654126315732 | 0.12955441458447126 |
| 0.040654126315732 | 0.12955441458447126 |
| 0.04406817876370406 | 0.1394242680713578 |
| 0.04506909232552398 | 0.1394242680713578 |
| 0.04506909232552398 | 0.1394242680713578 |
| 0.04506909232552398 | 0.1394242680713578 |
| 0.04506909232552398 | 0.1394242680713578 |
| 0.04946396091014192 | 0.14948889729840087 |
| 0.04946396091014192 | 0.149488897298401 |
| 0.04946396091014192 | 0.14948889729840087 |
| 0.04946396091014192 | 0.14948889729840087 |
| 0.04973543653028082 | 0.14948889729840087 |
