## Supplemental Table S2 for "Temporal analysis of tear fluid proteome reveals critical corneal repair events after photorefractive surgery"

| term |
| --- |
| Cytoplasmic Translation (GO:0002181) |
| Macromolecule Biosynthetic Process (GO:0009059) |
| Translation (GO:0006412) |
| Gene Expression (GO:0010467) |
| Protein Metabolic Process (GO:0019538) |
| Establishment of Endothelial Barrier (GO:0061028) |
| Negative Regulation of Complement Activation (GO:0045916) |
| Positive Regulation of Cytoplasmic Translation (GO:2000767) |
| Endothelial Cell Development (GO:0001885) |
| Regulation of Non-Motile Cilium Assembly (GO:1902855) |
| Regulation of Cytoplasmic Translation (GO:2000765) |
| Positive Regulation of Apoptotic Cell Clearance (GO:2000427) |
| Regulation of Apoptotic Cell Clearance (GO:2000425) |
| Positive Regulation of Early Endosome to Late Endosome Transport (GO:2000643) |
| Cytoskeleton-Dependent Cytokinesis (GO:0061640) |
| Regulation of Cellular Component Biogenesis (GO:0044087) |
| Regulation of Protein Localization to Early Endosome (GO:1902965) |
| Positive Regulation of Protein Localization to Early Endosome (GO:1902966) |
| Positive Regulation of Protein Localization to Endosome (GO:1905668) |
| CRD-mediated mRNA Stabilization (GO:0070934) |
| Regulation of Receptor-Mediated Endocytosis (GO:0048259) |
| Regulation of Organelle Organization (GO:0033043) |
| - Reg of Nuclear-Transcribed mRNA Cat Proc, Deadenylation-Dep Decay (GO:1900152) |
| Positive Regulation of Intracellular Transport (GO:0032388) |
| Positive Regulation of Translation (GO:0045727) |
| Positive Regulation of Heart Rate (GO:0010460) |
| Epiboly Involved in Wound Healing (GO:0090505) |
| Regulation of Immune Effector Process (GO:0002697) |
| GTP Metabolic Process (GO:0046039) |
| Protein Processing (GO:0016485) |
| Ribonucleoprotein Complex Biogenesis (GO:0022613) |
| mRNA Stabilization (GO:0048255) |
| Ruffle Organization (GO:0031529) |
| Regulation of DNA Strand Elongation (GO:0060382) |
| Positive Regulation of Regulatory T Cell Differentiation (GO:0045591) |
| Lymphocyte Mediated Immunity (GO:0002449) |
| Regulation of Early Endosome to Late Endosome Transport (GO:2000641) |
| Protein Localization to Plasma Membrane (GO:0072659) |
| Positive Regulation of T Cell Proliferation (GO:0042102) |
| Positive Regulation of Activated T Cell Proliferation (GO:0042104) |
| Protein Localization to Cell Periphery (GO:1990778) |
| T Cell Mediated Immunity (GO:0002456) |
| Regulation of Nuclear-Transcribed mRNA Catabolic Process, Deadenylation-Dependent Decay (GO:19 |
| Positive Regulation of Blood Coagulation (GO:0030194) |
| Negative Regulation of Peptidase Activity (GO:0010466) |

| |
|---|
| Adaptive Imm Resp Based on Som Recomb of Imm Rcptrs Built Frm IgSF Domains (GO:0002460) |
| Negative Regulation of Receptor-Mediated Endocytosis (GO:0048261) |
| Regulation of Gene Expression (GO:0010468) |
| Antigen Processing and Presentation of Exogenous Peptide Antigen via MHC Class II (GO:0019886) |
| Regulation of Vascular Endothelial Growth Factor Signaling Pathway (GO:1900746) |
| Positive Regulation of Cilium Assembly (GO:0045724) |
| Positive Regulation of Macromolecule Biosynthetic Process (GO:0010557) |
| Regulation of Cell Size (GO:0008361) |
| Ribosome Biogenesis (GO:0042254) |
| Positive Regulation of Signal Transduction (GO:0009967) |
| Antigen Processing and Presentation of Peptide Antigen via MHC Class II (GO:0002495) |
| Wound Healing, Spreading of Cells (GO:0044319) |
| Zymogen Activation (GO:0031638) |
| Protein Localization (GO:0008104) |
| Positive Regulation of Protein Kinase Activity (GO:0045860) |
| Positive Regulation of Response to Stimulus (GO:0048584) |
| Antigen Processing and Presentation of Exogenous Peptide Antigen (GO:0002478) |
| Positive Regulation of ATP-dependent Activity (GO:0032781) |
| Positive Regulation of Immune Response (GO:0050778) |
| Regulation of Activated T Cell Proliferation (GO:0046006) |
| Regulation of calcineurin-NFAT Signaling Cascade (GO:0070884) |
| rRNA Metabolic Process (GO:0016072) |
| Negative Regulation of Endocytosis (GO:0045806) |
| Positive Regulation of Protein Catabolic Process (GO:0045732) |
| RNA Processing (GO:0006396) |
| Negative Regulation of Hydrolase Activity (GO:0051346) |
| Positive Regulation of Immune Effector Process (GO:0002699) |
| Regulation of Intracellular Protein Transport (GO:0033157) |
| Regulation of Potassium Ion Transport (GO:0043266) |
| Cell-Matrix Adhesion (GO:0007160) |
| Protein Localization to Membrane (GO:0072657) |
| Positive Regulation of Protein Metabolic Process (GO:0051247) |
| rRNA Processing (GO:0006364) |
| Regulation of Potassium Ion Transmembrane Transport (GO:1901379) |
| Positive Regulation of Immune System Process (GO:0002684) |
| RNA Stabilization (GO:0043489) |
| Positive Regulation of Gene Expression (GO:0010628) |
| T Cell Activation (GO:0042110) |
| Regulation of Apoptotic Process (GO:0042981) |
| Regulation of Cellular Component Size (GO:0032535) |
| Regulation of Chromosome Organization (GO:0033044) |
| Regulation of Signal Transduction (GO:0009966) |
| Regulation of DNA Metabolic Process (GO:0051052) |
| Negative Regulation of mRNA Catabolic Process (GO:1902373) |
| Positive Regulation of Receptor-Mediated Endocytosis (GO:0048260) |
| Regulation of Monoatomic Cation Transmembrane Transport (GO:1904062) |

| |
|---|
| Positive Regulation of Chemotaxis (GO:0050921) |
| Regulation of ERK1 and ERK2 Cascade (GO:0070372) |
| Ribosomal Large Subunit Biogenesis (GO:0042273) |
| Blood Circulation (GO:0008015) |
| Positive Regulation of Kinase Activity (GO:0033674) |
| Positive Regulation of Leukocyte Cell-Cell Adhesion (GO:1903039) |
| Positive Regulation of Cell Migration (GO:0030335) |
| Regulation of Protein Catabolic Process (GO:0042176) |
| Negative Regulation of Apoptotic Process (GO:0043066) |
| tRNA Transport (GO:0051031) |
| Positive Regulation of Kidney Development (GO:0090184) |
| Embryonic Brain Development (GO:1990403) |
| Kidney Morphogenesis (GO:0060993) |
| Negative Regulation of Cell Size (GO:0045792) |
| Positive Regulation of Osteoblast Proliferation (GO:0033690) |
| Ruffle Assembly (GO:0097178) |
| Peptidyl-Serine Dephosphorylation (GO:0070262) |
| + Reg of CD4-+, CD25-+, Alpha-Beta Regulatory T Cell Differentiation (GO:0032831) |
| Regulation of T-Circle Formation (GO:1904429) |
| Response to Testosterone (GO:0033574) |
| Cell Junction Disassembly (GO:0150146) |
| RNA Decapping (GO:0110154) |
| Negative Regulation of Macrophage Differentiation (GO:0045650) |
| Regulation of Endoplasmic Reticulum Tubular Network Organization (GO:1903371) |
| Regulation of Lipid Metabolic Process (GO:0019216) |
| Cellular Response to Ionizing Radiation (GO:0071479) |
| Positive Regulation of MAPK Cascade (GO:0043410) |
| - Reg of Blood Vessel Endothelial Cell Proliferation Inv in Sprouting Angiogenesis (GO:1903588) |
| Regulation of T-helper Cell Differentiation (GO:0045622) |
| Random Inactivation of X Chromosome (GO:0060816) |
| Humoral Immune Response Mediated by Circulating Immunoglobulin (GO:0002455) |
| Gland Morphogenesis (GO:0022612) |
| Vascular Associated Smooth Muscle Contraction (GO:0014829) |
| Telomerase RNA Localization to Cajal Body (GO:0090671) |
| Regulation of Blood-Brain Barrier Permeability (GO:1905603) |
| Positive Regulation of Calcium Ion Import Across Plasma Membrane (GO:1905665) |
| Phospholipase C-activating Dopamine Receptor Signaling Pathway (GO:0060158) |
| Regulation of Saliva Secretion (GO:0046877) |
| Negative Regulation of Lipase Activity (GO:0060192) |
| Negative Regulation of Granulocyte Differentiation (GO:0030853) |
| Regulation of Cortisol Biosynthetic Process (GO:2000064) |
| Telomerase RNA Localization (GO:0090672) |
| Negative Regulation of Complement Activation, Classical Pathway (GO:0045959) |
| Positive Regulation of Leukocyte Chemotaxis (GO:0002690) |
| Regulation of Cell Adhesion (GO:0030155) |
| Positive Regulation of Cell Motility (GO:2000147) |

| |
|---|
| Regulation of Embryonic Development (GO:0045995) |
| Regulation of Bone Mineralization (GO:0030500) |
| Negative Regulation of Programmed Cell Death (GO:0043069) |
| Protein Localization to Cytoplasmic Stress Granule (GO:1903608) |
| Regulation of CD4-positive, Alpha-Beta T Cell Activation (GO:2000514) |
| Leukocyte Aggregation (GO:0070486) |
| Immunological Synapse Formation (GO:0001771) |
| Establishment of Golgi Localization (GO:0051683) |
| Deadenylation-Dependent Decapping of Nuclear-Transcribed mRNA (GO:0000290) |
| Positive Regulation of Attachment of Mitotic Spindle Microtubules to Kinetochore (GO:1902425) |
| Negative Regulation of Regulated Secretory Pathway (GO:1903306) |
| Cellular Response to Thyroid Hormone Stimulus (GO:0097067) |
| Positive Regulation of CD4-positive, Alpha-Beta T Cell Differentiation (GO:0043372) |
| Regulatory ncRNA-mediated Heterochromatin Formation (GO:0031048) |
| Regulation of Positive Chemotaxis (GO:0050926) |
| Antigen Processing and Presentation of Exogenous Peptide Antigen via MHC Class I (GO:0042590) |
| Regulation of Complement Activation, Classical Pathway (GO:0030450) |
| Positive Regulation of IRE1-mediated Unfolded Protein Response (GO:1903896) |
| Telomere Organization (GO:0032200) |
| Positive Regulation of ERK1 and ERK2 Cascade (GO:0070374) |
| Positive Regulation of Phagocytosis (GO:0050766) |
| Positive Regulation of T Cell Differentiation (GO:0045582) |
| Regulation of Small GTPase Mediated Signal Transduction (GO:0051056) |
| Positive Regulation of Memory T Cell Differentiation (GO:0043382) |
| Positive Regulation of Transforming Growth Factor Beta Production (GO:0071636) |
| Establishment of Protein Localization to Chromatin (GO:0071169) |
| Synapse Pruning (GO:0098883) |
| Endosome to Melanosome Transport (GO:0035646) |
| Detection of Visible Light (GO:0009584) |
| Endosome to Pigment Granule Transport (GO:0043485) |
| Positive Regulation by Host of Viral Genome Replication (GO:0044829) |
| Nucleic Acid Transport (GO:0050657) |
| Box C/D snoRNP Assembly (GO:0000492) |
| Negative Regulation of Ubiquitin Protein Ligase Activity (GO:1904667) |
| Negative Regulation of Humoral Immune Response Mediated by Circulating Immunoglobulin (GO:00[illegible] |
| Response to Thyroid Hormone (GO:0097066) |
| Epithelium Development (GO:0060429) |
| Regulation of Cellular Component Organization (GO:0051128) |
| Positive Regulation of MAP Kinase Activity (GO:0043406) |
| Plasma Membrane Bounded Cell Projection Assembly (GO:0120031) |
| Positive Regulation of Intracellular Protein Transport (GO:0090316) |
| Vasoconstriction (GO:0042310) |
| Positive Regulation of Vasculogenesis (GO:2001214) |
| Nerve Growth Factor Signaling Pathway (GO:0038180) |
| Regulation of Attachment of Mitotic Spindle Microtubules to Kinetochore (GO:1902423) |
| Regulation of Calcium Ion Import Across Plasma Membrane (GO:1905664) |

| |
|---|
| Positive Regulation of Keratinocyte Migration (GO:0051549) |
| Protein Localization to Microtubule (GO:0035372) |
| Sex-Chromosome Dosage Compensation (GO:0007549) |
| Rap Protein Signal Transduction (GO:0032486) |
| Phototransduction, Visible Light (GO:0007603) |
| Regulation of Store-Operated Calcium Entry (GO:2001256) |
| Regulation of Memory T Cell Differentiation (GO:0043380) |
| Regulation of Ketone Biosynthetic Process (GO:0010566) |
| Serine Family Amino Acid Biosynthetic Process (GO:0009070) |
| Negative Regulation of Necroptotic Process (GO:0060546) |
| Negative Regulation of Epidermal Growth Factor-Activated Receptor Activity (GO:0007175) |
| Positive Regulation of Endothelial Cell Proliferation (GO:0001938) |
| Regulation of Actin Filament Polymerization (GO:0030833) |
| Telomere Maintenance (GO:0000723) |
| Regulation of Macromolecule Biosynthetic Process (GO:0010556) |
| Positive Regulation of Protein Phosphorylation (GO:0001934) |
| Positive Regulation of Protein Transport (GO:0051222) |
| Regulation of DNA Replication (GO:0006275) |
| Regulation of MAP Kinase Activity (GO:0043405) |
| Establishment of Endothelial Intestinal Barrier (GO:0090557) |
| Establishment of Epithelial Cell Apical/Basal Polarity (GO:0045198) |
| Small Nucleolar Ribonucleoprotein Complex Assembly (GO:0000491) |
| Homologous Recombination (GO:0035825) |
| Negative Regulation of Calcium Ion Import (GO:0090281) |
| Very-Low-Density Lipoprotein Particle Assembly (GO:0034379) |
| Response to X-ray (GO:0010165) |
| Positive Regulation of Attachment of Spindle Microtubules to Kinetochore (GO:0051987) |
| Regulation of Mononuclear Cell Migration (GO:0071675) |
| Regulation of Muscle System Process (GO:0090257) |
| Negative Regulation of Programmed Necrotic Cell Death (GO:0062099) |
| Regulation of Lamellipodium Morphogenesis (GO:2000392) |
| RNA Localization to Cajal Body (GO:0090670) |
| L-serine Metabolic Process (GO:0006563) |
| Regulation of Granulocyte Differentiation (GO:0030852) |
| Negative Regulation of Cholesterol Transport (GO:0032375) |
| Positive Regulation of Protein Serine/Threonine Kinase Activity (GO:0071902) |
| Positive Regulation of Secretion by Cell (GO:1903532) |
| Regulation of Plasma Membrane Bounded Cell Projection Assembly (GO:0120032) |
| Regulation of Notch Signaling Pathway (GO:0008593) |
| Positive Regulation of Vascular Endothelial Growth Factor Receptor Signaling Pathway (GO:0030949) |
| Regulation of Chromatin Organization (GO:1902275) |
| Negative Regulation of Calcineurin-Mediated Signaling (GO:0106057) |
| Negative Regulation of calcineurin-NFAT Signaling Cascade (GO:0070885) |
| Negative Regulation of Adenylate Cyclase Activity (GO:0007194) |
| Purine Ribonucleoside Triphosphate Biosynthetic Process (GO:0009206) |
| Macrophage Chemotaxis (GO:0048246) |

| |
|---|
| mRNA Methylguanosine-Cap Decapping (GO:0110156) |
| Positive Regulation of Vascular Endothelial Growth Factor Signaling Pathway (GO:1900748) |
| Positive Regulation of Endoplasmic Reticulum Unfolded Protein Response (GO:1900103) |
| Dosage Compensation by Inactivation of X Chromosome (GO:0009048) |
| Negative Regulation of Homotypic Cell-Cell Adhesion (GO:0034111) |
| Skeletal Muscle Fiber Development (GO:0048741) |
| Phasic Smooth Muscle Contraction (GO:0014821) |
| Cellular Hyperosmotic Response (GO:0071474) |
| Pentose-Phosphate Shunt (GO:0006098) |
| Negative Regulation of Ubiquitin-Protein Transferase Activity (GO:0051444) |
| Regulation of Keratinocyte Migration (GO:0051547) |
| Golgi Inheritance (GO:0048313) |
| Skeletal Muscle Tissue Regeneration (GO:0043403) |
| Regulation of Epidermal Growth Factor-Activated Receptor Activity (GO:0007176) |
| Negative Regulation of Fibrinolysis (GO:0051918) |

| p-value | q-value |
|---|---|
| 3,64744E-14 | 3,0602E-11 |
| 1,84435E-13 | 7,73706E-11 |
| 3,04938E-12 | 8,52811E-10 |
| 1,85727E-11 | 3,89562E-09 |
| 1,26032E-07 | 2,11482E-05 |
| 7,8877E-06 | 0,001102964 |
| 2,01088E-05 | 0,002410181 |
| 4,11746E-05 | 0,004318185 |
| 0,000177748 | 0,016570027 |
| 0,000207426 | 0,017299921 |
| 0,000226817 | 0,017299921 |
| 0,000432989 | 0,027944466 |
| 0,000432989 | 0,027944466 |
| 0,000575591 | 0,032581068 |
| 0,000610797 | 0,032581068 |
| 0,000653486 | 0,032581068 |
| 0,000737831 | 0,032581068 |
| 0,000737831 | 0,032581068 |
| 0,000737831 | 0,032581068 |
| 0,00091953 | 0,037939291 |
| 0,000949613 | 0,037939291 |
| 0,001048131 | 0,039971916 |
| 0,001120508 | 0,04087419 |
| 0,001489036 | 0,052054231 |
| 0,001560135 | 0,052358129 |
| 0,001837357 | 0,053156627 |
| 0,001837357 | 0,053156627 |
| 0,001837357 | 0,053156627 |
| 0,001837357 | 0,053156627 |
| 0,001964116 | 0,054929787 |
| 0,00229353 | 0,059431705 |
| 0,002302971 | 0,059431705 |
| 0,002408436 | 0,059431705 |
| 0,002408436 | 0,059431705 |
| 0,002721412 | 0,063424027 |
| 0,002721412 | 0,063424027 |
| 0,003052452 | 0,069216413 |
| 0,003235655 | 0,071344074 |
| 0,00334817 | 0,071344074 |
| 0,003401386 | 0,071344074 |
| 0,003600623 | 0,071849797 |
| 0,003768047 | 0,071849797 |
| 0,003768047 | 0,071849797 |
| 0,003768047 | 0,071849797 |
| 0,004972723 | 0,092713661 |

| | |
|---|---|
| 0,005408632 | 0,095020485 |
| 0,005408632 | 0,095020485 |
| 0,005436214 | 0,095020485 |
| 0,005861444 | 0,095981047 |
| 0,005861444 | 0,095981047 |
| 0,005861444 | 0,095981047 |
| 0,005948766 | 0,095981047 |
| 0,006330998 | 0,098613371 |
| 0,006529758 | 0,098613371 |
| 0,006583128 | 0,098613371 |
| 0,006817134 | 0,098613371 |
| 0,006817134 | 0,098613371 |
| 0,006817134 | 0,098613371 |
| 0,007325904 | 0,104176837 |
| 0,007797289 | 0,109032096 |
| 0,008806505 | 0,118849441 |
| 0,008924332 | 0,118849441 |
| 0,008924332 | 0,118849441 |
| 0,009611381 | 0,125999193 |
| 0,010073338 | 0,128053496 |
| 0,010073338 | 0,128053496 |
| 0,010457854 | 0,130705551 |
| 0,010671154 | 0,130705551 |
| 0,010749324 | 0,130705551 |
| 0,011082968 | 0,132837291 |
| 0,011912653 | 0,138815493 |
| 0,011912653 | 0,138815493 |
| 0,013214309 | 0,148229808 |
| 0,013214309 | 0,148229808 |
| 0,013250579 | 0,148229808 |
| 0,013463059 | 0,148625082 |
| 0,014152939 | 0,153461348 |
| 0,014266967 | 0,153461348 |
| 0,01457494 | 0,154789548 |
| 0,015277004 | 0,155904014 |
| 0,015277004 | 0,155904014 |
| 0,015343186 | 0,155904014 |
| 0,015689338 | 0,155904014 |
| 0,015775331 | 0,155904014 |
| 0,015993376 | 0,155904014 |
| 0,015993376 | 0,155904014 |
| 0,016356019 | 0,155904014 |
| 0,016429452 | 0,155904014 |
| 0,016723911 | 0,155904014 |
| 0,016723911 | 0,155904014 |
| 0,017468467 | 0,161055423 |

| | |
|---|---|
| 0,018999076 | 0,16627862 |
| 0,019612465 | 0,16627862 |
| 0,019784849 | 0,16627862 |
| 0,020584083 | 0,16627862 |
| 0,020584083 | 0,16627862 |
| 0,020584083 | 0,16627862 |
| 0,021316453 | 0,16627862 |
| 0,021713472 | 0,16627862 |
| 0,022348891 | 0,16627862 |
| 0,022791468 | 0,16627862 |
| 0,022791468 | 0,16627862 |
| 0,022791468 | 0,16627862 |
| 0,022791468 | 0,16627862 |
| 0,022791468 | 0,16627862 |
| 0,022791468 | 0,16627862 |
| 0,022791468 | 0,16627862 |
| 0,022791468 | 0,16627862 |
| 0,022791468 | 0,16627862 |
| 0,022791468 | 0,16627862 |
| 0,022791468 | 0,16627862 |
| 0,022791468 | 0,16627862 |
| 0,022791468 | 0,16627862 |
| 0,022791468 | 0,16627862 |
| 0,022791468 | 0,16627862 |
| 0,023061178 | 0,166545186 |
| 0,024777382 | 0,166545186 |
| 0,025851537 | 0,166545186 |
| 0,027287734 | 0,166545186 |
| 0,027287734 | 0,166545186 |
| 0,027287734 | 0,166545186 |
| 0,027287734 | 0,166545186 |
| 0,027287734 | 0,166545186 |
| 0,027287734 | 0,166545186 |
| 0,027287734 | 0,166545186 |
| 0,027287734 | 0,166545186 |
| 0,027287734 | 0,166545186 |
| 0,027287734 | 0,166545186 |
| 0,027287734 | 0,166545186 |
| 0,027287734 | 0,166545186 |
| 0,027287734 | 0,166545186 |
| 0,027287734 | 0,166545186 |
| 0,027287734 | 0,166545186 |
| 0,027287734 | 0,166545186 |
| 0,02744623 | 0,166545186 |
| 0,027821228 | 0,166545186 |
| 0,027967228 | 0,166545186 |

| | |
|---|---|
| 0,030225389 | 0,166545186 |
| 0,030225389 | 0,166545186 |
| 0,03156991 | 0,166545186 |
| 0,031763539 | 0,166545186 |
| 0,031763539 | 0,166545186 |
| 0,031763539 | 0,166545186 |
| 0,031763539 | 0,166545186 |
| 0,031763539 | 0,166545186 |
| 0,031763539 | 0,166545186 |
| 0,031763539 | 0,166545186 |
| 0,031763539 | 0,166545186 |
| 0,031763539 | 0,166545186 |
| 0,031763539 | 0,166545186 |
| 0,031763539 | 0,166545186 |
| 0,031763539 | 0,166545186 |
| 0,031763539 | 0,166545186 |
| 0,031763539 | 0,166545186 |
| 0,031763539 | 0,166545186 |
| 0,032137734 | 0,166545186 |
| 0,033630777 | 0,166545186 |
| 0,035093085 | 0,166545186 |
| 0,035093085 | 0,166545186 |
| 0,035887904 | 0,166545186 |
| 0,036218972 | 0,166545186 |
| 0,036218972 | 0,166545186 |
| 0,036218972 | 0,166545186 |
| 0,036218972 | 0,166545186 |
| 0,036218972 | 0,166545186 |
| 0,036218972 | 0,166545186 |
| 0,036218972 | 0,166545186 |
| 0,036218972 | 0,166545186 |
| 0,036218972 | 0,166545186 |
| 0,036218972 | 0,166545186 |
| 0,036218972 | 0,166545186 |
| 0,036218972 | 0,166545186 |
| 0,036218972 | 0,166545186 |
| 0,037045161 | 0,166545186 |
| 0,037045161 | 0,166545186 |
| 0,037119765 | 0,166545186 |
| 0,039024202 | 0,166545186 |
| 0,040242019 | 0,166545186 |
| 0,040654126 | 0,166545186 |
| 0,040654126 | 0,166545186 |
| 0,040654126 | 0,166545186 |
| 0,040654126 | 0,166545186 |
| 0,040654126 | 0,166545186 |

| | |
|---|---|
| 0,040654126 | 0,166545186 |
| 0,040654126 | 0,166545186 |
| 0,040654126 | 0,166545186 |
| 0,040654126 | 0,166545186 |
| 0,040654126 | 0,166545186 |
| 0,040654126 | 0,166545186 |
| 0,040654126 | 0,166545186 |
| 0,040654126 | 0,166545186 |
| 0,040654126 | 0,166545186 |
| 0,040654126 | 0,166545186 |
| 0,040654126 | 0,166545186 |
| 0,041304221 | 0,166545186 |
| 0,04237695 | 0,166545186 |
| 0,043460091 | 0,166545186 |
| 0,043748705 | 0,166545186 |
| 0,044386277 | 0,166545186 |
| 0,044553527 | 0,166545186 |
| 0,044553527 | 0,166545186 |
| 0,044553527 | 0,166545186 |
| 0,045069092 | 0,166545186 |
| 0,045069092 | 0,166545186 |
| 0,045069092 | 0,166545186 |
| 0,045069092 | 0,166545186 |
| 0,045069092 | 0,166545186 |
| 0,045069092 | 0,166545186 |
| 0,045069092 | 0,166545186 |
| 0,045069092 | 0,166545186 |
| 0,045069092 | 0,166545186 |
| 0,045069092 | 0,166545186 |
| 0,045069092 | 0,166545186 |
| 0,045069092 | 0,166545186 |
| 0,045069092 | 0,166545186 |
| 0,045069092 | 0,166545186 |
| 0,045069092 | 0,166545186 |
| 0,045069092 | 0,166545186 |
| 0,046770823 | 0,166545186 |
| 0,046770823 | 0,166545186 |
| 0,046770823 | 0,166545186 |
| 0,047894455 | 0,166545186 |
| 0,049463961 | 0,166545186 |
| 0,049463961 | 0,166545186 |
| 0,049463961 | 0,166545186 |
| 0,049463961 | 0,166545186 |
| 0,049463961 | 0,166545186 |
| 0,049463961 | 0,166545186 |
| 0,049463961 | 0,166545186 |

| | |
|---|---|
| 0,049463961 | 0,166545186 |
| 0,049463961 | 0,166545186 |
| 0,049463961 | 0,166545186 |
| 0,049463961 | 0,166545186 |
| 0,049463961 | 0,166545186 |
| 0,049463961 | 0,166545186 |
| 0,049463961 | 0,166545186 |
| 0,049463961 | 0,166545186 |
| 0,049463961 | 0,166545186 |
| 0,049463961 | 0,166545186 |
| 0,049463961 | 0,166545186 |
| 0,049463961 | 0,166545186 |
| 0,049463961 | 0,166545186 |
| 0,049463961 | 0,166545186 |
| 0,049463961 | 0,166545186 |
