## Supplemental Table S3 for "Temporal analysis of tear fluid proteome reveals critical corneal repair events after photorefractive surgery"

| term |
| --- |
| Cadherin Binding (GO:0045296) |
| GTPase Activity (GO:0003924) |
| Ribonucleoside Triphosphate Phosphatase Activity (GO:0017111) |
| Actin Binding (GO:0003779) |
| Phosphatase Inhibitor Activity (GO:0019212) |
| Protein Phosphatase Inhibitor Activity (GO:0004864) |
| Guanyl Ribonucleotide Binding (GO:0032561) |
| GTP Binding (GO:0005525) |
| GDP Binding (GO:0019003) |
| Serine-Type Endopeptidase Activity (GO:0004252) |
| Double-Stranded RNA Binding (GO:0003725) |
| Serine-Type Peptidase Activity (GO:0008236) |
| Protein Phosphatase Regulator Activity (GO:0019888) |
| Low-Density Lipoprotein Particle Receptor Binding (GO:0050750) |
| Lipoprotein Particle Receptor Binding (GO:0070325) |
| mRNA 3'-UTR Binding (GO:0003730) |
| Endopeptidase Activity (GO:0004175) |
| Regulatory RNA Binding (GO:0061980) |
| Endopeptidase Inhibitor Activity (GO:0004866) |
| Phosphatidylcholine-Sterol O-acyltransferase Activator Activity (GO:0060228) |
| Oxidoreductase Activity, Acting on a Sulfur Group of Donors, Oxygen as Acceptor (GO:0016670) |
| Calcium-Dependent Protein Serine/Threonine Phosphatase Activity (GO:0004723) |
| Insulin-Like Growth Factor II Binding (GO:0031995) |
| RNA Polymerase II C-terminal Domain Binding (GO:0099122) |
| Serine-Type Exopeptidase Activity (GO:0070008) |
| G Protein-Coupled Receptor Binding (GO:0001664) |
| Serine-Type Endopeptidase Inhibitor Activity (GO:0004867) |
| 5'-Deoxyribose-5-Phosphate Lyase Activity (GO:0051575) |
| G Protein-Coupled Glutamate Receptor Binding (GO:0035256) |
| poly(G) Binding (GO:0034046) |
| Double-Stranded Telomeric DNA Binding (GO:0003691) |
| MHC Class II Receptor Activity (GO:0032395) |
| Phospholipase Inhibitor Activity (GO:0004859) |
| 5S rRNA Binding (GO:0008097) |
| Anion Binding (GO:0043168) |
| Thioesterase Binding (GO:0031996) |
| Lipase Inhibitor Activity (GO:0055102) |
| U3 snoRNA Binding (GO:0034511) |
| Oxidoreductase Activity, Acting on the CH-OH Group of Donors, NAD or NADP as Acceptor (GO:0016 |
| Ubiquitin Ligase Inhibitor Activity (GO:1990948) |
| ATPase Binding (GO:0051117) |

| p-value | q-value |
|---|---|
| 2,28935E-07 | 2,2998E-05 |
| 4,04576E-07 | 2,2998E-05 |
| 5,30722E-07 | 2,2998E-05 |
| 2,50273E-05 | 0,000813386 |
| 0,000385661 | 0,010027197 |
| 0,000508319 | 0,011013575 |
| 0,00070947 | 0,013175871 |
| 0,002630144 | 0,042739842 |
| 0,003496037 | 0,046808103 |
| 0,003600623 | 0,046808103 |
| 0,004295107 | 0,050078726 |
| 0,004978206 | 0,050078726 |
| 0,005007873 | 0,050078726 |
| 0,005408632 | 0,05022301 |
| 0,007838517 | 0,067933812 |
| 0,012596853 | 0,102349429 |
| 0,014425294 | 0,110311069 |
| 0,016723911 | 0,120256305 |
| 0,017575921 | 0,120256305 |
| 0,022791468 | 0,137642001 |
| 0,027287734 | 0,137642001 |
| 0,027287734 | 0,137642001 |
| 0,027287734 | 0,137642001 |
| 0,027287734 | 0,137642001 |
| 0,027287734 | 0,137642001 |
| 0,02832515 | 0,137642001 |
| 0,031175689 | 0,137642001 |
| 0,031763539 | 0,137642001 |
| 0,031763539 | 0,137642001 |
| 0,031763539 | 0,137642001 |
| 0,036218972 | 0,137680735 |
| 0,036218972 | 0,137680735 |
| 0,036218972 | 0,137680735 |
| 0,036218972 | 0,137680735 |
| 0,03802782 | 0,137680735 |
| 0,040654126 | 0,137680735 |
| 0,040654126 | 0,137680735 |
| 0,040654126 | 0,137680735 |
| 0,041304221 | 0,137680735 |
| 0,045069092 | 0,144766549 |
| 0,045657142 | 0,144766549 |
