## Supplemental Table S4 for "Temporal analysis of tear fluid proteome reveals critical corneal repair events after photorefractive surgery"

| term |
| --- |
| Scavenging of Heme From Plasma |
| Classical Antibody-Mediated Complement Activation |
| CD22 Mediated BCR Regulation |
| FCGR Activation |
| Role of LAT2 NTAL LAB on Calcium Mobilization |
| Creation of C4 and C2 Activators |
| Initial Triggering of Complement |
| Role of Phospholipids in Phagocytosis |
| Binding and Uptake of Ligands by Scavenger Receptors |
| FCERI Mediated Ca+2 Mobilization |
| FCERI Mediated MAPK Activation |
| Regulation of Complement Cascade |
| FCGR3A-mediated IL10 Synthesis |
| Complement Cascade |
| Antigen Activates B Cell Receptor (BCR) Leading to Generation of Second Messengers |
| Parasite Infection |
| Leishmania Phagocytosis |
| FCGR3A-mediated Phagocytosis |
| Regulation of Actin Dynamics for Phagocytic Cup Formation |
| FCERI Mediated NF-kB Activation |
| Anti-inflammatory Response Favouring Leishmania Parasite Infection |
| Leishmania Parasite Growth and Survival |
| Cell Surface Interactions at the Vascular Wall |
| Fcgamma Receptor (FCGR) Dependent Phagocytosis |
| Fc Epsilon Receptor (FCERI) Signaling |
| Signaling by the B Cell Receptor (BCR) |
| Potential Therapeutics for SARS |
| Immunoregulatory Interactions Between a Lymphoid and a non-Lymphoid Cell |
| Leishmania Infection |
| Parasitic Infection Pathways |
| Hemostasis |
| Vesicle-mediated Transport |
| SARS-CoV Infections |
| Adaptive Immune System |
| Innate Immune System |
| Viral Infection Pathways |
| Infectious Disease |
| Immune System |
| Disease |
| Post-translational Modification Synthesis of GPI-anchored Proteins |
| Inhibition of Signaling by Overexpressed EGFR |
| Signaling by Overexpressed Wild-Type EGFR in Cancer |
| EGFR Interacts With Phospholipase C-gamma |
| Negative Regulation of Activity of TFAP2 (AP-2) Family Transcription Factors |
| Extra-nuclear Estrogen Signaling |

| |
|---|
| Ribavirin ADME |
| NFE2L2 Regulating Tumorigenic Genes |

| p-value | q-value |
|---|---|
| 5,42983E-61 | 7,60176E-59 |
| 1,81674E-59 | 1,27172E-57 |
| 7,55065E-59 | 3,52364E-57 |
| 2,90845E-58 | 1,01796E-56 |
| 4,51766E-58 | 1,26494E-56 |
| 6,97669E-58 | 1,62789E-56 |
| 1,86382E-56 | 3,72764E-55 |
| 5,90103E-56 | 1,03268E-54 |
| 7,66857E-56 | 1,19289E-54 |
| 1,79729E-55 | 2,51621E-54 |
| 2,58406E-55 | 3,2888E-54 |
| 5,3718E-55 | 6,2671E-54 |
| 7,95849E-54 | 8,57069E-53 |
| 1,46193E-53 | 1,46193E-52 |
| 1,60489E-53 | 1,4979E-52 |
| 3,98794E-51 | 3,10173E-50 |
| 3,98794E-51 | 3,10173E-50 |
| 3,98794E-51 | 3,10173E-50 |
| 5,21797E-51 | 3,84482E-50 |
| 1,49714E-50 | 1,048E-49 |
| 2,93158E-49 | 1,86555E-48 |
| 2,93158E-49 | 1,86555E-48 |
| 1,28632E-48 | 7,82975E-48 |
| 2,31767E-48 | 1,35197E-47 |
| 8,44761E-46 | 4,73066E-45 |
| 4,87052E-45 | 2,62259E-44 |
| 7,48786E-45 | 3,88259E-44 |
| 7,7383E-44 | 3,86915E-43 |
| 2,29539E-42 | 1,07118E-41 |
| 2,29539E-42 | 1,07118E-41 |
| 9,68362E-32 | 4,37325E-31 |
| 1,88239E-29 | 8,23545E-29 |
| 7,39638E-29 | 3,13786E-28 |
| 4,52663E-25 | 1,86391E-24 |
| 7,80737E-21 | 3,12295E-20 |
| 3,34576E-19 | 1,30113E-18 |
| 5,36621E-19 | 2,03046E-18 |
| 5,75039E-14 | 2,11856E-13 |
| 2,60944E-13 | 9,36722E-13 |
| 0,008284521 | 0,028995823 |
| 0,035056276 | 0,116854254 |
| 0,035056276 | 0,116854254 |
| 0,039351982 | 0,128122731 |
| 0,043628778 | 0,137098038 |
| 0,044067227 | 0,137098038 |

| | |
|---|---|
| 0,047886748 | 0,142641378 |
| 0,047886748 | 0,142641378 |
