## Supplemental Table S5 for "Temporal analysis of tear fluid proteome reveals critical corneal repair events after photorefractive surgery"

| term |
| --- |
| Defense Response to Bacterium (GO:0042742) |
| Negative Regulation of Collateral Sprouting (GO:0048671) |
| ERBB2-EGFR Signaling Pathway (GO:0038134) |
| Regulation of Hyaluronan Biosynthetic Process (GO:1900125) |
| Response to Cocaine (GO:0042220) |
| Antibacterial Humoral Response (GO:0019731) |
| Negative Regulation of Dendritic Spine Development (GO:0061000) |
| Positive Regulation of Mitochondrial Calcium Ion Concentration (GO:0051561) |
| Positive Regulation of Pyroptotic Inflammatory Response (GO:0140639) |
| Positive Regulation of Respiratory Burst (GO:0060267) |
| Negative Regulation of Axon Regeneration (GO:0048681) |
| Negative Regulation of Neuron Projection Regeneration (GO:0070571) |
| Regulation of Apoptotic Cell Clearance (GO:2000425) |
| Regulation of Collateral Sprouting (GO:0048670) |
| Cellular Response to Monoamine Stimulus (GO:0071868) |
| Nucleoside Triphosphate Metabolic Process (GO:0009141) |
| Positive Regulation of Potassium Ion Transmembrane Transporter Activity (GO:1901018) |
| Regulation of Cell Killing (GO:0031341) |
| Positive Regulation of Endopeptidase Activity (GO:0010950) |
| Positive Regulation of Epithelial Tube Formation (GO:1905278) |
| Regulation of Complement-Dependent Cytotoxicity (GO:1903659) |
| Regulation of Respiratory Burst (GO:0060263) |
| Granulocyte Migration (GO:0097530) |
| Positive Regulation of Protein Localization to Early Endosome (GO:1902966) |
| Positive Regulation of Protein Localization to Endosome (GO:1905668) |
| Regulation of Protein Localization to Early Endosome (GO:1902965) |
| Phagocytosis (GO:0006909) |
| Dopamine Transport (GO:0015872) |
| Glomerular Filtration (GO:0003094) |
| Regulation of Toll-Like Receptor 9 Signaling Pathway (GO:0034163) |
| Negative Regulation of Interferon-Beta Production (GO:0032688) |
| Regulation of Complement Activation (GO:0030449) |
| Regulation of Axon Regeneration (GO:0048679) |

| p-value | q-value |
|---|---|
| 0,017068545 | 0,291621421 |
| 0,022054881 | 0,291621421 |
| 0,022054881 | 0,291621421 |
| 0,022054881 | 0,291621421 |
| 0,022054881 | 0,291621421 |
| 0,023295961 | 0,291621421 |
| 0,026407809 | 0,291621421 |
| 0,026407809 | 0,291621421 |
| 0,026407809 | 0,291621421 |
| 0,026407809 | 0,291621421 |
| 0,026407809 | 0,291621421 |
| 0,030741579 | 0,291621421 |
| 0,030741579 | 0,291621421 |
| 0,030741579 | 0,291621421 |
| 0,030741579 | 0,291621421 |
| 0,035056276 | 0,291621421 |
| 0,035056276 | 0,291621421 |
| 0,035056276 | 0,291621421 |
| 0,035056276 | 0,291621421 |
| 0,035056276 | 0,291621421 |
| 0,039351982 | 0,291621421 |
| 0,039351982 | 0,291621421 |
| 0,039351982 | 0,291621421 |
| 0,039351982 | 0,291621421 |
| 0,039351982 | 0,291621421 |
| 0,039351982 | 0,291621421 |
| 0,039912151 | 0,291621421 |
| 0,043628778 | 0,291621421 |
| 0,043628778 | 0,291621421 |
| 0,043628778 | 0,291621421 |
| 0,043628778 | 0,291621421 |
| 0,043628778 | 0,291621421 |
| 0,047886748 | 0,291621421 |
