## Supplemental Table S6 for "Temporal analysis of tear fluid proteome reveals critical corneal repair events after photorefractive surgery"

| term | p-value | q-value |
|---|---|---|
| Potassium Channel Activator Activity (GO:0099104) | 0,026407809 | 0,400078479 |
| Pyrophosphatase Activity (GO:0016462) | 0,028434682 | 0,400078479 |
| Immunoglobulin Receptor Binding (GO:0034987) | 0,030741579 | 0,400078479 |
| Channel Activator Activity (GO:0099103) | 0,039351982 | 0,400078479 |
| Nucleoside Triphosphate Diphosphatase Activity (GO:0047429) | 0,039351982 | 0,400078479 |
| GDP Phosphatase Activity (GO:0004382) | 0,039351982 | 0,400078479 |
