## Supplemental Table S7 for "Temporal analysis of tear fluid proteome reveals critical corneal repair events after photorefractive surgery"

| term |
| --- |
| L13a-mediated Translational Silencing of Ceruloplasmin Expression |
| Formation of a Pool of Free 40S Subunits |
| GTP Hydrolysis and Joining of the 60S Ribosomal Subunit |
| Cap-dependent Translation Initiation |
| Eukaryotic Translation Initiation |
| Eukaryotic Translation Elongation |
| Peptide Chain Elongation |
| Nonsense Mediated Decay (NMD) Independent of the Exon Junction Complex (EJC) |
| Response of EIF2AK4 (GCN2) to Amino Acid Deficiency |
| Nonsense Mediated Decay (NMD) Enhanced by the Exon Junction Complex (EJC) |
| Nonsense-Mediated Decay (NMD) |
| Eukaryotic Translation Termination |
| Selenocysteine Synthesis |
| Selenoamino Acid Metabolism |
| Viral mRNA Translation |
| Metabolism of Proteins |
| SRP-dependent Cotranslational Protein Targeting to Membrane |
| Regulation of Expression of SLITs and ROBOs |
| Translation |
| Influenza Infection |
| Signaling by ROBO Receptors |
| Cellular Response to Starvation |
| Influenza Viral RNA Transcription and Replication |
| Formation of the Ternary Complex, and Subsequently, the 43S Complex |
| Translation Initiation Complex Formation |
| mRNA Activation Upon Binding of the Cap-Binding Complex and eIFs, and Subsequent Binding to 43S |
| Ribosomal Scanning and Start Codon Recognition |
| Major Pathway of rRNA Processing in the Nucleolus and Cytosol |
| Infectious Disease |
| Nervous System Development |
| Axon Guidance |
| rRNA Processing in the Nucleus and Cytosol |
| Metabolism of RNA |
| Cellular Responses to Stress |
| Cellular Responses to Stimuli |
| Viral Infection Pathways |
| Disease |
| rRNA Processing |
| SARS-CoV-2 Modulates Host Translation Machinery |
| Metabolism of Amino Acids and Derivatives |
| SARS-CoV-1 Modulates Host Translation Machinery |
| Developmental Biology |
| Innate Immune System |
| SARS-CoV-1-host Interactions |
| SARS-CoV-2-host Interactions |

| |
|---|
| SARS-CoV-2 Infection |
| Vesicle-mediated Transport |
| Neutrophil Degranulation |
| SARS-CoV-1 Infection |
| Immune System |
| Membrane Trafficking |
| Metabolism |
| SARS-CoV Infections |
| Platelet Degranulation |
| Post-translational Protein Modification |
| Response to Elevated Platelet Cytosolic Ca2+ |
| Regulation of IGF Transport and Uptake by Insulin-like Growth Factor Binding Proteins (IGFBPs) |
| mRNA Splicing - Major Pathway |
| mRNA Splicing |
| Post-translational Protein Phosphorylation |
| Hemostasis |
| RAB Geranylgeranylation |
| COPI-mediated Anterograde Transport |
| Platelet Activation, Signaling and Aggregation |
| Golgi Associated Vesicle Biogenesis |
| Processing of Capped Intron-Containing Pre-mRNA |
| Asparagine N-linked Glycosylation |
| Complement Cascade |
| ER to Golgi Anterograde Transport |
| Regulation of Complement Cascade |
| Transport to the Golgi and Subsequent Modification |
| Formation of Fibrin Clot (Clotting Cascade) |
| trans-Golgi Network Vesicle Budding |
| Gene and Protein Expression by JAK-STAT Signaling After Interleukin-12 Stimulation |
| Signaling by Rho GTPases |
| Apoptosis |
| Plasma Lipoprotein Assembly, Remodeling, and Clearance |
| Signaling by Rho GTPases, Miro GTPases and RHOBTB3 |
| Chylomicron Assembly |
| Chylomicron Remodeling |
| RHOBTB2 GTPase Cycle |
| Amyloid Fiber Formation |
| Interleukin-12 Family Signaling |
| Programmed Cell Death |
| RHOBTB GTPase Cycle |
| Interleukin-12 Signaling |
| Plasma Lipoprotein Assembly |
| Apoptotic Execution Phase |
| ER-Phagosome Pathway |
| Terminal Pathway of Complement |
| Formation of the Cornified Envelope |

| |
|---|
| RNA Polymerase II Transcription Termination |
| Chaperone Mediated Autophagy |
| Common Pathway of Fibrin Clot Formation |
| Intrinsic Pathway of Fibrin Clot Formation |
| Retinoid Metabolism and Transport |
| Golgi-to-ER Retrograde Transport |
| Metabolism of Fat-Soluble Vitamins |
| Clathrin-mediated Endocytosis |
| Plasma Lipoprotein Clearance |
| Antigen processing-Cross Presentation |
| RHO GTPase Cycle |
| Intra-Golgi and Retrograde Golgi-to-ER Traffic |
| Dissolution of Fibrin Clot |
| RAB GEFs Exchange GTP for GDP on RABs |
| Regulation of TLR by Endogenous Ligand |
| MHC Class II Antigen Presentation |
| M Phase |
| Signaling by Interleukins |
| Plasma Lipoprotein Remodeling |
| Lysosome Vesicle Biogenesis |
| Cellular Response to Chemical Stress |
| Binding and Uptake of Ligands by Scavenger Receptors |
| Rab Regulation of Trafficking |
| MAP2K and MAPK Activation |
| Cell Cycle, Mitotic |
| Signal Transduction |
| AUF1 (hnRNP D0) Binds and Destabilizes mRNA |
| HDL Remodeling |
| SLBP Independent Processing of Histone Pre-mRNAs |
| Scavenging by Class A Receptors |
| Regulation of mRNA Stability by Proteins That Bind AU-rich Elements |
| Antigen Presentation Folding, Assembly and Peptide Loading of Class I MHC |
| TBC RABGAPs |
| SLBP Dependent Processing of Replication-Dependent Histone Pre-mRNAs |
| Cytokine Signaling in Immune System |
| Visual Phototransduction |
| Nef-mediates Down Modulation of Cell Surface Receptors by Recruiting Them to Clathrin Adapters |
| COPI-dependent Golgi-to-ER Retrograde Traffic |
| Signaling by BRAF and RAF1 Fusions |
| N-glycan Trimming in the ER and Calnexin Calreticulin Cycle |
| RUNX1 Regulates Transcription of Genes Involved in Differentiation of HSCs |
| Signaling by High-Kinase Activity BRAF Mutants |
| Apoptosis Induced DNA Fragmentation |
| Host Interactions of HIV Factors |
| RHOV GTPase Cycle |
| Cytosolic tRNA Aminoacylation |

| |
|---|
| Keratinization |
| Apoptotic Cleavage of Cellular Proteins |
| Golgi Cisternae Pericentriolar Stack Reorganization |
| ABC-family Proteins Mediated Transport |
| Mitotic Prophase |
| Respiratory Syncytial Virus Infection Pathway |
| EPH-Ephrin Signaling |
| p130Cas Linkage to MAPK Signaling for Integrins |
| GRB2 SOS Provides Linkage to MAPK Signaling for Integrins |
| Signaling by ALK Fusions and Activated Point Mutants |
| Signaling by ALK in Cancer |
| Cell-Cell Communication |
| Signaling by RAF1 Mutants |
| Cell Cycle |
| mRNA 3'-End Processing |
| Regulation of CDH19 Expression and Function |
| Scavenging by Class F Receptors |
| Activation of C3 and C5 |
| CDH11 Homotypic and Heterotypic Interactions |
| VLDLR Internalisation and Degradation |
| Signaling by Receptor Tyrosine Kinases |
| Diseases of Signal Transduction by Growth Factor Receptors and Second Messengers |
| Diseases of Programmed Cell Death |
| Signaling by WNT |
| KEAP1-NFE2L2 Pathway |
| The Role of Nef in HIV-1 Replication and Disease Pathogenesis |
| Processing of Capped Intronless Pre-mRNA |
| Transport of Small Molecules |
| RHO GTPase Effectors |
| Paradoxical Activation of RAF Signaling by Kinase Inactive BRAF |
| Signaling by RAS Mutants |
| Signaling by Moderate Kinase Activity BRAF Mutants |
| Signaling Downstream of RAS Mutants |
| Respiratory Syncytial Virus (RSV) Genome Replication, Transcription and Translation |
| RHO GTPases Activate PKNs |
| UCH Proteinases |
| Oncogenic MAPK Signaling |
| Prevention of Phagosomal-Lysosomal Fusion |
| Initial Triggering of Complement |
| Respiratory Syncytial Virus (RSV) Attachment and Entry |
| LDL Clearance |
| MyD88 Deficiency (TLR2 4) |
| Nuclear Events Mediated by NFE2L2 |
| HIV Infection |
| IRAK4 Deficiency (TLR2 4) |
| Formation of ATP by Chemiosmotic Coupling |

| |
|---|
| Autophagy |
| InlA-mediated Entry of Listeria Monocytogenes Into Host Cells |
| G2 M Transition |
| RHO GTPases Activate WASPs and WAVEs |
| mRNA Splicing - Minor Pathway |
| Mitotic G2-G2 M Phases |
| Unfolded Protein Response (UPR) |
| Nef Mediated Downregulation of MHC Class I Complex Cell Surface Expression |
| Regulation of CDH11 Function |
| Apoptotic Cleavage of Cell Adhesion Proteins |
| Nephrin Family Interactions |
| RHOBTB1 GTPase Cycle |
| Suppression of Phagosomal Maturation |
| HSP90 Chaperone Cycle for Steroid Hormone Receptors (SHR) in the Presence of Ligand |
| Mitochondrial Protein Degradation |
| Regulation of Endogenous Retroelements by KRAB-ZFP Proteins |
| Platelet Aggregation (Plug Formation) |
| Regulation of Actin Dynamics for Phagocytic Cup Formation |
| Parasitic Infection Pathways |
| Leishmania Infection |
| Fcgamma Receptor (FCGR) Dependent Phagocytosis |
| Replacement of Protamines by Nucleosomes in the Male Pronucleus |
| VEGFA-VEGFR2 Pathway |
| ALK Mutants Bind TKIs |
| Josephin Domain DUBs |
| Vitamin D (Calciferol) Metabolism |
| Glycogen Metabolism |
| PCP CE Pathway |
| PRC2 Methylates Histones and DNA |
| Defective Pyroptosis |
| Assembly of the Pre-Replicative Complex |
| HDACs Deacetylate Histones |
| tRNA Aminoacylation |
| Calnexin Calreticulin Cycle |
| WNT5A-dependent Internalization of FZD2, FZD5 and ROR2 |
| p38MAPK Events |
| Advanced Glycosylation Endproduct Receptor Signaling |
| Hh Mutants Are Degraded by ERAD |
| Formation of Incision Complex in GG-NER |
| Aggrephagy |
| MAPK Family Signaling Cascades |
| Integrin Signaling |
| Smooth Muscle Contraction |
| Erythropoietin Activates RAS |
| Glycogen Breakdown (Glycogenolysis) |
| Retrograde Neurotrophin Signalling |

| |
|---|
| FGFR2 Alternative Splicing |
| ERCC6 (CSB) and EHMT2 (G9a) Positively Regulate rRNA Expression |
| Adaptive Immune System |
| Signaling by VEGF |
| ABC Transporter Disorders |
| The Role of GTSE1 in G2 M Progression After G2 Checkpoint |
| Maternal to Zygotic Transition (MZT) |
| RAF MAP Kinase Cascade |
| Hh Mutants Abrogate Ligand Secretion |
| Regulation of PLK1 Activity at G2 M Transition |
| VEGFR2 Mediated Vascular Permeability |
| Pentose Phosphate Pathway |
| mRNA Decay by 5' to 3' Exoribonuclease |
| WNT5A-dependent Internalization of FZD4 |
| RAC2 GTPase Cycle |
| Degradation of GLI1 by the Proteasome |
| Degradation of GLI2 by the Proteasome |
| GLI3 Is Processed to GLI3R by the Proteasome |
| Selective Autophagy |
| Alternative Complement Activation |
| Chylomicron Clearance |
| HDL Clearance |
| G2 M Checkpoints |
| MAPK1 MAPK3 Signaling |
| Defective CFTR Causes Cystic Fibrosis |
| COPII-mediated Vesicle Transport |
| Defects of Contact Activation System (CAS) and Kallikrein Kinin System (KKS) |
| RNA Polymerase I Promoter Opening |
| Signaling by NTRK1 (TRKA) |
| Cell-cell Junction Organization |
| Scavenging of Heme From Plasma |
| Translocation of SLC2A4 (GLUT4) to the Plasma Membrane |
| Parasite Infection |
| FCGR3A-mediated Phagocytosis |
| Leishmania Phagocytosis |
| RAC3 GTPase Cycle |
| Transcriptional and Post-Translational Regulation of MITF-M Expression and Activity |
| Protein Methylation |
| DNA Replication Pre-Initiation |
| DNA Methylation |
| Cristae Formation |
| PERK Regulates Gene Expression |
| Cell Junction Organization |
| COPI-independent Golgi-to-ER Retrograde Traffic |
| Hedgehog Ligand Biogenesis |
| L1CAM Interactions |

| |
|---|
| Assembly of Viral Components at the Budding Site |
| Scavenging by Class B Receptors |
| Uptake and Function of Diphtheria Toxin |
| RHOG GTPase Cycle |
| Transport of Mature mRNA Derived From an Intron-Containing Transcript |
| Signalling to ERKs |
| Late Endosomal Microautophagy |
| Deposition of New CENPA-containing Nucleosomes at the Centromere |
| Nucleosome Assembly |
| Signaling by FGFR2 |
| snRNP Assembly |
| Metabolism of Non-Coding RNA |
| Transcriptional Regulation by RUNX1 |
| Diseases of Immune System |
| Activated PKN1 Stimulates Transcription of AR (Androgen Receptor) Regulated Genes KLK2 and KLK3 |
| Diseases Associated With the TLR Signaling Cascade |
| Regulation of RAS by GAPs |
| Intrinsic Pathway for Apoptosis |
| Constitutive Signaling by Ligand-Responsive EGFR Cancer Variants |
| Diseases of Hemostasis |
| SARS-CoV-1 Targets Host Intracellular Signalling and Regulatory Pathways |
| Signaling by Ligand-Responsive EGFR Variants in Cancer |
| Detoxification of Reactive Oxygen Species |
| Striated Muscle Contraction |
| Cellular Response to Heat Stress |
| Regulation of PTEN Stability and Activity |
| Deadenylation-dependent mRNA Decay |
| Epithelial-Mesenchymal Transition (EMT) During Gastrulation |
| SUMO Is Transferred From E1 to E2 (UBE2I, UBC9) |
| Assembly of the ORC Complex at the Origin of Replication |
| Regulation of Activated PAK-2p34 by Proteasome Mediated Degradation |
| Signalling to RAS |
| Organelle Biogenesis and Maintenance |
| Processing of DNA Double-Strand Break Ends |
| MAPK6 MAPK4 Signaling |
| Regulation of Ornithine Decarboxylase (ODC) |
| DNA Damage Recognition in GG-NER |
| Cargo Recognition for Clathrin-Mediated Endocytosis |
| Adherens Junctions Interactions |
| Meiotic Synapsis |
| RHOQ GTPase Cycle |
| CD209 (DC-SIGN) Signaling |
| Factors Involved in Megakaryocyte Development and Platelet Production |
| Ubiquitin-dependent Degradation of Cyclin D |
| Ubiquitin Mediated Degradation of Phosphorylated Cdc25A |
| SIRT1 Negatively Regulates rRNA Expression |

| |
|---|
| Response of Mtb to Phagocytosis |
| Autodegradation of the E3 Ubiquitin Ligase COP1 |
| Chromatin Modifications During the Maternal to Zygotic Transition (MZT) |
| p53-Independent G1 S DNA Damage Checkpoint |
| p53-Independent DNA Damage Response |
| Regulation of Endogenous Retroelements by the Human Silencing Hub (HUSH) Complex |
| Formation of the Beta-Catenin TCF Transactivating Complex |
| DNA Damage Telomere Stress Induced Senescence |
| Recruitment and ATM-med Phosphorylation of Repair and Signaling Proteins at DNA Double Strand B |
| Transport of Mature Transcript to Cytoplasm |
| Defective B4GALT1 Causes B4GALT1-CDG (CDG-2d) |
| Defective CHST6 Causes MCDC1 |
| Defective ST3GAL3 Causes MCT12 and EIEE15 |
| Erythrocytes Take up Oxygen and Release Carbon Dioxide |
| HDL Assembly |
| NFE2L2 Regulates Pentose Phosphate Pathway Genes |
| Recycling of eIF2 GDP |
| Integration of Energy Metabolism |
| Vpu Mediated Degradation of CD4 |
| RHOU GTPase Cycle |
| Regulation of Apoptosis |
| Beta-catenin Independent WNT Signaling |
| Signaling by NTRKs |
| Regulation of Signaling by CBL |
| Deregulated CDK5 Triggers Multiple Neurodegenerative Pathways in Alzheimer's Disease Models |
| Listeria Monocytogenes Entry Into Host Cells |
| Neurodegenerative Diseases |
| DNA Double Strand Break Response |
| Global Genome Nucleotide Excision Repair (GG-NER) |
| TCF Dependent Signaling in Response to WNT |
| Integrin Cell Surface Interactions |
| Transcriptional Regulation by RUNX3 |
| Regulation of Endogenous Retroelements |
| Translation of Structural Proteins 9694635 |
| Vif-mediated Degradation of APOBEC3G |
| SCF-beta-TrCP Mediated Degradation of Emi1 |
| RND3 GTPase Cycle |
| GSK3B and BTRC CUL1-mediated-degradation of NFE2L2 |
| FBXL7 Down-Regulates AURKA During Mitotic Entry and in Early Mitosis |
| EPHB-mediated Forward Signaling |
| Degradation of AXIN |
| Condensation of Prophase Chromosomes |
| Cellular Response to Mitochondrial Stress |
| Malate-aspartate Shuttle |
| Myoclonic Epilepsy of Lafora |
| ROBO Receptors Bind AKAP5 |

| |
|---|
| Respiratory Syncytial Virus Genome Replication |
| Classical Antibody-Mediated Complement Activation |
| DARPP-32 Events |
| Meiosis |
| Metabolism of Vitamins and Cofactors |
| Regulation of Endogenous Retroelements by Piwi-interacting RNAs (piRNAs) |
| Degradation of DVL |
| Negative Regulation of NOTCH4 Signaling |
| Stabilization of P53 |
| Signaling by Erythropoietin |
| Signaling by EGFR in Cancer |
| Defective Intrinsic Pathway for Apoptosis |
| Bacterial Infection Pathways |
| DNA Replication |
| Macroautophagy |
| Signaling by FGFR |
| GPER1 Signaling |
| Regulation of RUNX3 Expression and Activity |
| Virus Assembly and Release |
| Processing and Activation of SUMO |
| Prednisone ADME |
| Nef Mediated CD4 Down-regulation |
| Entry of Influenza Virion Into Host Cell via Endocytosis |
| ATF6 (ATF6-alpha) Activates Chaperone Genes |
| Downstream Signaling Events of B Cell Receptor (BCR) |
| Downregulation of TGF-beta Receptor Signaling |
| WNT Ligand Biogenesis and Trafficking |
| Metabolism of Polyamines |
| NIK-->noncanonical NF-kB Signaling |
| Loss of Proteins Required for Interphase Microtubule Organization From the Centrosome |
| Loss of Nlp From Mitotic Centrosomes |
| Muscle Contraction |
| Oxidative Stress Induced Senescence |
| Heme Signaling |
| Inhibition of DNA Recombination at Telomere |
| SUMOylation of RNA Binding Proteins |
| Attenuation Phase |
| Deubiquitination |
| Creation of C4 and C2 Activators |
| RMTs Methylate Histone Arginines |
| Interleukin-3, Interleukin-5 and GM-CSF Signaling |
| Assembly and Release of Respiratory Syncytial Virus (RSV) Virions |
| Constitutive Signaling by Overexpressed ERBB2 |
| Formation of Annular Gap Junctions |
| Extracellular Matrix Organization |
| Signaling by NOTCH |

| |
|---|
| Anchoring of the Basal Body to the Plasma Membrane |
| Mitochondrial Biogenesis |
| FLT3 Signaling in Disease |
| AURKA Activation by TPX2 |
| Dectin-1 Mediated Noncanonical NF-kB Signaling |
| Cyclin E Associated Events During G1 S Transition |
| Hedgehog 'Off' State |
| SCF(Skp2)-mediated Degradation of P27 P21 |
| Regulation of Localization of FOXO Transcription Factors |
| O2 CO2 Exchange in Erythrocytes |
| Gap Junction Degradation |
| Erythrocytes Take up Carbon Dioxide and Release Oxygen |
| Endosomal Vacuolar Pathway |
| Caspase-mediated Cleavage of Cytoskeletal Proteins |
| CREB1 Phosphorylation Through the Activation of Adenylate Cyclase |
| ATF6 (ATF6-alpha) Activates Chaperones |
| Z-decay Degradation of Maternal mRNAs by Zygotically Expressed Factors |
| Asymmetric Localization of PCP Proteins |
| EPH-ephrin Mediated Repulsion of Cells |
| IRE1alpha Activates Chaperones |
| Nonhomologous End-Joining (NHEJ) |
| Transcriptional Regulation by Small RNAs |
| Positive Epigenetic Regulation of rRNA Expression |
| Cyclin A Cdk2-associated Events at S Phase Entry |
| Class I MHC Mediated Antigen Processing & Presentation |
| Activation of BH3-only Proteins |
| HSF1 Activation |
| Regulation of CDH11 Expression and Function |
| Signaling by CSF3 (G-CSF) |
| Interleukin-1 Signaling |
| G2 M DNA Damage Checkpoint |
| Turbulent Flow Shear Stress Activates Signaling by PIEZO1 and Integrins in Endothelial Cells |
| Signaling by CSF1 (M-CSF) in Myeloid Cells |
| Interaction Between L1 and Ankyrins |
| Metalloprotease DUBs |
| RHO GTPases Activate IQGAPs |
| C-type Lectin Receptors (CLRs) |
| p53-Dependent G1 DNA Damage Response |
| p53-Dependent G1 S DNA Damage Checkpoint |
| Oxygen-dependent Proline Hydroxylation of Hypoxia-inducible Factor Alpha |
| Retinoid Cycle Disease Events |
| Diseases of the Neuronal System |
| Diseases Associated With Visual Transduction |
| eNOS Activation |
| Chk1 Chk2(Cds1) Mediated Inactivation of Cyclin B Cdk1 Complex |
| Ub-specific Processing Proteases |

| |
|---|
| Interferon Alpha Beta Signaling |
| Regulation of Insulin Secretion |
| Mitotic Anaphase |
| Activation of NF-kappaB in B Cells |
| Autodegradation of Cdh1 by Cdh1 APC C |
| Mitotic Metaphase and Anaphase |
| RNA Polymerase I Promoter Clearance |
| Packaging Of Telomere Ends |
| Transcriptional Regulation by RUNX2 |
| APC C Cdc20 Mediated Degradation of Securin |
| G1 S DNA Damage Checkpoints |
| RHOJ GTPase Cycle |
| Late SARS-CoV-2 Infection Events |
| Glycogen Synthesis |
| Regulation of Expression and Function of Type II Classical Cadherins |
| Regulation of Homotypic Cell-Cell Adhesion |
| Activation of Matrix Metalloproteinases |
| Centrosome Maturation |
| RNA Polymerase I Transcription |
| Recruitment of Mitotic Centrosome Proteins and Complexes |
| Senescence-Associated Secretory Phenotype (SASP) |
| Signaling by Hedgehog |
| HATs Acetylate Histones |
| Nucleotide Excision Repair |
| Metabolism of Carbohydrates |
| Cellular Senescence |
| Meiotic Recombination |
| E3 Ubiquitin Ligases Ubiquitinate Target Proteins |
| Citric Acid Cycle (TCA Cycle) |
| Diseases of Carbohydrate Metabolism |
| Cell Cycle Checkpoints |
| SARS-CoV-2 Targets Host Intracellular Signalling and Regulatory Pathways |
| Signaling by EGFRvIII in Cancer |
| Keratan Sulfate Degradation |
| Response of EIF2AK1 (HRI) to Heme Deficiency |
| Constitutive Signaling by EGFRvIII |
| Activation of BAD and Translocation to Mitochondria |
| Depolymerization of the Nuclear Lamina |
| Orc1 Removal From Chromatin |
| Signaling by ERBB4 |
| Transcriptional Regulation of Granulopoiesis |
| Translation of Structural Proteins 9683701 |

| p-value | q-value |
|---|---|
| 3,8431E-70 | 4,80003E-67 |
| 3,39852E-68 | 1,74723E-65 |
| 4,1967E-68 | 1,74723E-65 |
| 1,06254E-67 | 2,65423E-65 |
| 1,06254E-67 | 2,65423E-65 |
| 9,30345E-62 | 1,93667E-59 |
| 3,52762E-58 | 6,29429E-56 |
| 5,40763E-58 | 8,44266E-56 |
| 6,93143E-58 | 9,61929E-56 |
| 2,44248E-57 | 2,77333E-55 |
| 2,44248E-57 | 2,77333E-55 |
| 6,57089E-57 | 6,31311E-55 |
| 6,57089E-57 | 6,31311E-55 |
| 7,82176E-57 | 6,97813E-55 |
| 1,02767E-55 | 8,55708E-54 |
| 1,66152E-54 | 1,29702E-52 |
| 2,01847E-54 | 1,48298E-52 |
| 1,83892E-52 | 1,276E-50 |
| 1,77107E-51 | 1,16425E-49 |
| 3,38902E-50 | 2,11644E-48 |
| 5,89088E-48 | 3,50367E-46 |
| 2,02671E-47 | 1,15062E-45 |
| 1,02148E-46 | 5,54707E-45 |
| 1,50166E-44 | 7,81489E-43 |
| 1,61648E-43 | 8,07595E-42 |
| 3,86046E-43 | 1,85451E-41 |
| 1,03799E-41 | 4,80168E-40 |
| 1,1322E-40 | 5,05044E-39 |
| 3,0064E-40 | 1,29483E-38 |
| 4,64453E-40 | 1,93367E-38 |
| 9,24645E-40 | 3,72542E-38 |
| 2,06101E-39 | 8,04437E-38 |
| 1,13199E-38 | 4,28441E-37 |
| 2,67069E-38 | 9,81085E-37 |
| 5,73211E-38 | 2,04554E-36 |
| 9,78225E-38 | 3,3939E-36 |
| 3,85505E-37 | 1,30134E-35 |
| 2,93242E-35 | 9,63841E-34 |
| 1,17172E-34 | 3,7525E-33 |
| 1,3907E-34 | 4,34246E-33 |
| 1,20782E-32 | 3,67944E-31 |
| 1,81975E-30 | 5,4116E-29 |
| 4,53398E-23 | 1,31696E-21 |
| 2,74064E-22 | 7,77969E-21 |
| 3,37766E-22 | 9,37487E-21 |

| | |
|---|---|
| 1,1996E-21 | 3,25716E-20 |
| 4,62587E-21 | 1,2293E-19 |
| 5,21584E-21 | 1,3572E-19 |
| 5,66332E-20 | 1,44357E-18 |
| 8,53093E-20 | 2,13103E-18 |
| 3,25723E-18 | 7,97702E-17 |
| 1,76028E-17 | 4,22806E-16 |
| 2,52824E-17 | 5,95805E-16 |
| 3,12994E-16 | 7,23944E-15 |
| 6,31408E-16 | 1,43387E-14 |
| 8,09171E-16 | 1,80474E-14 |
| 1,58764E-14 | 3,47889E-13 |
| 6,28146E-14 | 1,35268E-12 |
| 2,38585E-13 | 5,05073E-12 |
| 7,06271E-12 | 1,47022E-10 |
| 1,69768E-11 | 3,47606E-10 |
| 2,27952E-11 | 4,59213E-10 |
| 2,37825E-11 | 4,71498E-10 |
| 3,05451E-11 | 5,96106E-10 |
| 3,31226E-11 | 6,36463E-10 |
| 4,4775E-11 | 8,47333E-10 |
| 5,2087E-11 | 9,70995E-10 |
| 1,0633E-10 | 1,95304E-09 |
| 1,15164E-10 | 2,08463E-09 |
| 1,80518E-10 | 3,22096E-09 |
| 5,96892E-10 | 1,05002E-08 |
| 1,06327E-09 | 1,84448E-08 |
| 1,20687E-09 | 2,0649E-08 |
| 9,78422E-09 | 1,65142E-07 |
| 1,10073E-08 | 1,83308E-07 |
| 1,87619E-08 | 3,08338E-07 |
| 2,04747E-08 | 3,32116E-07 |
| 2,15361E-08 | 3,44854E-07 |
| 3,28498E-08 | 5,10527E-07 |
| 3,28498E-08 | 5,10527E-07 |
| 3,31086E-08 | 5,10527E-07 |
| 4,57728E-08 | 6,97198E-07 |
| 6,57416E-08 | 9,89292E-07 |
| 7,55799E-08 | 1,1238E-06 |
| 8,76916E-08 | 1,28855E-06 |
| 9,46269E-08 | 1,37429E-06 |
| 1,55943E-07 | 2,23877E-06 |
| 3,23458E-07 | 4,5909E-06 |
| 3,73813E-07 | 5,24598E-06 |
| 3,8252E-07 | 5,30853E-06 |
| 4,1185E-07 | 5,65276E-06 |

| | |
|---|---|
| 4,48932E-07 | 6,09474E-06 |
| 4,96294E-07 | 6,59438E-06 |
| 4,96294E-07 | 6,59438E-06 |
| 6,98941E-07 | 9,18923E-06 |
| 7,28893E-07 | 9,4832E-06 |
| 7,44096E-07 | 9,5812E-06 |
| 1,58562E-06 | 2,02086E-05 |
| 1,85571E-06 | 2,34119E-05 |
| 1,94885E-06 | 2,43411E-05 |
| 2,75405E-06 | 3,40575E-05 |
| 3,59921E-06 | 4,40727E-05 |
| 7,63094E-06 | 9,25345E-05 |
| 7,96898E-06 | 9,57044E-05 |
| 8,91822E-06 | 0,000106084 |
| 9,16559E-06 | 0,000107953 |
| 9,24818E-06 | 0,000107953 |
| 9,82138E-06 | 0,000113582 |
| 1,12709E-05 | 0,00012915 |
| 1,22447E-05 | 0,000139034 |
| 1,49964E-05 | 0,000168743 |
| 2,64209E-05 | 0,00029464 |
| 3,28907E-05 | 0,000363544 |
| 3,56796E-05 | 0,000390911 |
| 3,75428E-05 | 0,000407748 |
| 4,39538E-05 | 0,000473261 |
| 5,01701E-05 | 0,000535577 |
| 5,21558E-05 | 0,000552056 |
| 5,79252E-05 | 0,000602905 |
| 5,79252E-05 | 0,000602905 |
| 6,40182E-05 | 0,000660816 |
| 7,5034E-05 | 0,000766737 |
| 7,55073E-05 | 0,000766737 |
| 8,25288E-05 | 0,000831278 |
| 8,93255E-05 | 0,00089254 |
| 9,95911E-05 | 0,000987217 |
| 0,000118362 | 0,00116405 |
| 0,000136731 | 0,001334191 |
| 0,000139971 | 0,001355221 |
| 0,000142811 | 0,001372087 |
| 0,000153659 | 0,001465039 |
| 0,000179813 | 0,001697354 |
| 0,000180743 | 0,001697354 |
| 0,000186386 | 0,001737282 |
| 0,000203447 | 0,001882262 |
| 0,00021146 | 0,001934985 |
| 0,000212244 | 0,001934985 |

| | |
|---|---|
| 0,000215387 | 0,001949405 |
| 0,000246145 | 0,002211765 |
| 0,000256078 | 0,002284585 |
| 0,000278006 | 0,002462618 |
| 0,000304544 | 0,002678701 |
| 0,000326028 | 0,002845517 |
| 0,000328066 | 0,002845517 |
| 0,000342697 | 0,0029317 |
| 0,000342697 | 0,0029317 |
| 0,000355757 | 0,003002299 |
| 0,000355757 | 0,003002299 |
| 0,000366948 | 0,003075961 |
| 0,000377617 | 0,003144293 |
| 0,000399258 | 0,003302475 |
| 0,000419894 | 0,003431239 |
| 0,000428561 | 0,003431239 |
| 0,000428561 | 0,003431239 |
| 0,000428561 | 0,003431239 |
| 0,000428561 | 0,003431239 |
| 0,000448429 | 0,003567438 |
| 0,000472505 | 0,003735184 |
| 0,000501011 | 0,003935617 |
| 0,00050783 | 0,003964251 |
| 0,000516709 | 0,004008503 |
| 0,000531856 | 0,004097374 |
| 0,000538006 | 0,004097374 |
| 0,000538006 | 0,004097374 |
| 0,000558299 | 0,004226157 |
| 0,000604001 | 0,004544558 |
| 0,000631667 | 0,004599144 |
| 0,000631667 | 0,004599144 |
| 0,000631667 | 0,004599144 |
| 0,000631667 | 0,004599144 |
| 0,000633183 | 0,004599144 |
| 0,000633349 | 0,004599144 |
| 0,00071116 | 0,005104824 |
| 0,00071116 | 0,005104824 |
| 0,000726195 | 0,005182957 |
| 0,000809109 | 0,005741919 |
| 0,000860677 | 0,006073363 |
| 0,000902776 | 0,006299261 |
| 0,000902776 | 0,006299261 |
| 0,001052468 | 0,007302956 |
| 0,001068363 | 0,007372296 |
| 0,001107545 | 0,007559147 |
| 0,001107545 | 0,007559147 |

| | |
|---|---|
| 0,001373874 | 0,009325917 |
| 0,00139346 | 0,009407737 |
| 0,001486422 | 0,009959084 |
| 0,001491072 | 0,009959084 |
| 0,001518873 | 0,010090808 |
| 0,00162907 | 0,010765651 |
| 0,001859111 | 0,012182687 |
| 0,001882513 | 0,012182687 |
| 0,001882513 | 0,012182687 |
| 0,001882513 | 0,012182687 |
| 0,001913859 | 0,01219597 |
| 0,001913859 | 0,01219597 |
| 0,001913859 | 0,01219597 |
| 0,002024196 | 0,012833607 |
| 0,002122045 | 0,013320535 |
| 0,002122327 | 0,013320535 |
| 0,002148871 | 0,013419701 |
| 0,00216547 | 0,013456081 |
| 0,002187346 | 0,013458103 |
| 0,002187346 | 0,013458103 |
| 0,002236768 | 0,013694719 |
| 0,002254184 | 0,013734026 |
| 0,002413361 | 0,014632464 |
| 0,002466191 | 0,014738144 |
| 0,002466191 | 0,014738144 |
| 0,002466191 | 0,014738144 |
| 0,002633995 | 0,015665996 |
| 0,002649248 | 0,015682042 |
| 0,002690996 | 0,015779596 |
| 0,002690996 | 0,015779596 |
| 0,002735111 | 0,015963335 |
| 0,002883807 | 0,016752908 |
| 0,002996285 | 0,017325743 |
| 0,003055464 | 0,01758652 |
| 0,003150135 | 0,017884177 |
| 0,003150135 | 0,017884177 |
| 0,003150135 | 0,017884177 |
| 0,003325799 | 0,018627454 |
| 0,003325799 | 0,018627454 |
| 0,003325799 | 0,018627454 |
| 0,003468252 | 0,0193386 |
| 0,003520722 | 0,019543916 |
| 0,003680588 | 0,02034095 |
| 0,003939427 | 0,021486218 |
| 0,003939427 | 0,021486218 |
| 0,003939427 | 0,021486218 |

| | |
|---|---|
| 0,004031846 | 0,021894674 |
| 0,004061705 | 0,021907725 |
| 0,004069329 | 0,021907725 |
| 0,004131843 | 0,022148807 |
| 0,004310974 | 0,022912371 |
| 0,004310974 | 0,022912371 |
| 0,004369437 | 0,023124689 |
| 0,004422246 | 0,023305425 |
| 0,004470193 | 0,023459122 |
| 0,004548067 | 0,023767929 |
| 0,004590862 | 0,02389161 |
| 0,004838613 | 0,024791046 |
| 0,004838613 | 0,024791046 |
| 0,004838613 | 0,024791046 |
| 0,004843087 | 0,024791046 |
| 0,004907092 | 0,024813595 |
| 0,004907092 | 0,024813595 |
| 0,004907092 | 0,024813595 |
| 0,005152284 | 0,025948399 |
| 0,005323288 | 0,026422902 |
| 0,005323288 | 0,026422902 |
| 0,005323288 | 0,026422902 |
| 0,005346268 | 0,026422902 |
| 0,005371471 | 0,026422902 |
| 0,005373432 | 0,026422902 |
| 0,005782886 | 0,028324803 |
| 0,005851736 | 0,028480981 |
| 0,005860378 | 0,028480981 |
| 0,00602302 | 0,029157952 |
| 0,006169095 | 0,029635382 |
| 0,006169095 | 0,029635382 |
| 0,006642584 | 0,031787692 |
| 0,006727738 | 0,031829338 |
| 0,006727738 | 0,031829338 |
| 0,006727738 | 0,031829338 |
| 0,006925489 | 0,032641269 |
| 0,006959247 | 0,032662801 |
| 0,00698236 | 0,032662801 |
| 0,007009727 | 0,032668469 |
| 0,007344245 | 0,033816315 |
| 0,007344245 | 0,033816315 |
| 0,007344245 | 0,033816315 |
| 0,007364322 | 0,033816315 |
| 0,007553435 | 0,034431532 |
| 0,007553435 | 0,034431532 |
| 0,007732266 | 0,035118546 |

| | |
|---|---|
| 0,007860308 | 0,035314838 |
| 0,007860308 | 0,035314838 |
| 0,007860308 | 0,035314838 |
| 0,008100729 | 0,035983674 |
| 0,008100729 | 0,035983674 |
| 0,008170944 | 0,035983674 |
| 0,008170944 | 0,035983674 |
| 0,008182036 | 0,035983674 |
| 0,008182036 | 0,035983674 |
| 0,008634181 | 0,037838921 |
| 0,008846 | 0,038416843 |
| 0,008846 | 0,038416843 |
| 0,008858327 | 0,038416843 |
| 0,009056347 | 0,038870715 |
| 0,009056347 | 0,038870715 |
| 0,009056347 | 0,038870715 |
| 0,009546257 | 0,040405909 |
| 0,009546257 | 0,040405909 |
| 0,00960813 | 0,040405909 |
| 0,00960813 | 0,040405909 |
| 0,00960813 | 0,040405909 |
| 0,00960813 | 0,040405909 |
| 0,010002009 | 0,041780969 |
| 0,010002009 | 0,041780969 |
| 0,010120185 | 0,042133705 |
| 0,010283715 | 0,042530994 |
| 0,010283715 | 0,042530994 |
| 0,010833018 | 0,044508026 |
| 0,010833018 | 0,044508026 |
| 0,011009405 | 0,044937083 |
| 0,011009405 | 0,044937083 |
| 0,011108238 | 0,045192799 |
| 0,011539366 | 0,046794377 |
| 0,011683868 | 0,047074682 |
| 0,011683868 | 0,047074682 |
| 0,012079934 | 0,048358455 |
| 0,012079934 | 0,048358455 |
| 0,012367427 | 0,049351168 |
| 0,012728087 | 0,05017992 |
| 0,012728087 | 0,05017992 |
| 0,012728087 | 0,05017992 |
| 0,012735816 | 0,05017992 |
| 0,013028776 | 0,050475319 |
| 0,013214915 | 0,050475319 |
| 0,013214915 | 0,050475319 |
| 0,013214915 | 0,050475319 |

| | |
|---|---|
| 0,013214915 | 0,050475319 |
| 0,013214915 | 0,050475319 |
| 0,013214915 | 0,050475319 |
| 0,013214915 | 0,050475319 |
| 0,013214915 | 0,050475319 |
| 0,013214915 | 0,050475319 |
| 0,013623023 | 0,051561079 |
| 0,013623023 | 0,051561079 |
| 0,013623023 | 0,051561079 |
| 0,013840421 | 0,052014392 |
| 0,014219479 | 0,052014392 |
| 0,014219479 | 0,052014392 |
| 0,014219479 | 0,052014392 |
| 0,014219479 | 0,052014392 |
| 0,014219479 | 0,052014392 |
| 0,014219479 | 0,052014392 |
| 0,014219479 | 0,052014392 |
| 0,014273967 | 0,052014392 |
| 0,014415589 | 0,052014392 |
| 0,014415589 | 0,052014392 |
| 0,014415589 | 0,052014392 |
| 0,014454826 | 0,052014392 |
| 0,014454826 | 0,052014392 |
| 0,014492401 | 0,052014392 |
| 0,014492401 | 0,052014392 |
| 0,014492401 | 0,052014392 |
| 0,014492401 | 0,052014392 |
| 0,014559379 | 0,052105056 |
| 0,014617074 | 0,052162073 |
| 0,014752371 | 0,052494904 |
| 0,015423519 | 0,054727201 |
| 0,016260273 | 0,057532807 |
| 0,016381745 | 0,057798869 |
| 0,016559402 | 0,058261107 |
| 0,017018574 | 0,058557022 |
| 0,017018574 | 0,058557022 |
| 0,017018574 | 0,058557022 |
| 0,017018574 | 0,058557022 |
| 0,017018574 | 0,058557022 |
| 0,017018574 | 0,058557022 |
| 0,017018574 | 0,058557022 |
| 0,017018574 | 0,058557022 |
| 0,017998543 | 0,061017311 |
| 0,017998543 | 0,061017311 |
| 0,017998543 | 0,061017311 |
| 0,017998543 | 0,061017311 |

| | |
|---|---|
| 0,017998543 | 0,061017311 |
| 0,018026732 | 0,061017311 |
| 0,018397049 | 0,062102469 |
| 0,018957416 | 0,063821597 |
| 0,019753634 | 0,066094702 |
| 0,019888457 | 0,066094702 |
| 0,019897204 | 0,066094702 |
| 0,019897204 | 0,066094702 |
| 0,019897204 | 0,066094702 |
| 0,020546561 | 0,067259389 |
| 0,020546561 | 0,067259389 |
| 0,020546561 | 0,067259389 |
| 0,020570926 | 0,067259389 |
| 0,020570926 | 0,067259389 |
| 0,020570926 | 0,067259389 |
| 0,020916054 | 0,068209273 |
| 0,021442134 | 0,069561624 |
| 0,021442134 | 0,069561624 |
| 0,022149825 | 0,070754811 |
| 0,022149825 | 0,070754811 |
| 0,022149825 | 0,070754811 |
| 0,022149825 | 0,070754811 |
| 0,022149825 | 0,070754811 |
| 0,022149825 | 0,070754811 |
| 0,022334871 | 0,071163914 |
| 0,022828013 | 0,072365959 |
| 0,022828013 | 0,072365959 |
| 0,023058512 | 0,072727479 |
| 0,023058512 | 0,072727479 |
| 0,023627997 | 0,074149167 |
| 0,023627997 | 0,074149167 |
| 0,023714227 | 0,074233257 |
| 0,024103875 | 0,075264349 |
| 0,024747019 | 0,076697337 |
| 0,024747019 | 0,076697337 |
| 0,024747019 | 0,076697337 |
| 0,025241429 | 0,078036001 |
| 0,026334491 | 0,080998661 |
| 0,026400549 | 0,080998661 |
| 0,026508257 | 0,080998661 |
| 0,026508257 | 0,080998661 |
| 0,026653683 | 0,080998661 |
| 0,026653683 | 0,080998661 |
| 0,026653683 | 0,080998661 |
| 0,026952163 | 0,081706922 |
| 0,02742871 | 0,082950262 |

| | |
|---|---|
| 0,027601494 | 0,083070522 |
| 0,027601494 | 0,083070522 |
| 0,027786584 | 0,08324707 |
| 0,027793457 | 0,08324707 |
| 0,028342751 | 0,084689225 |
| 0,029278906 | 0,087277693 |
| 0,030110408 | 0,089542618 |
| 0,030250949 | 0,089746877 |
| 0,031491192 | 0,091470927 |
| 0,031491192 | 0,091470927 |
| 0,031491192 | 0,091470927 |
| 0,031491192 | 0,091470927 |
| 0,031491192 | 0,091470927 |
| 0,031491192 | 0,091470927 |
| 0,031491192 | 0,091470927 |
| 0,031491192 | 0,091470927 |
| 0,031491192 | 0,091470927 |
| 0,032233224 | 0,092596921 |
| 0,032233224 | 0,092596921 |
| 0,032233224 | 0,092596921 |
| 0,032233224 | 0,092596921 |
| 0,032397802 | 0,092596921 |
| 0,032397802 | 0,092596921 |
| 0,032397802 | 0,092596921 |
| 0,032796814 | 0,093523335 |
| 0,033270054 | 0,094014246 |
| 0,033270054 | 0,094014246 |
| 0,033270054 | 0,094014246 |
| 0,033270054 | 0,094014246 |
| 0,034146287 | 0,096272487 |
| 0,035716451 | 0,09993124 |
| 0,036206809 | 0,09993124 |
| 0,036206809 | 0,09993124 |
| 0,036206809 | 0,09993124 |
| 0,036206809 | 0,09993124 |
| 0,036206809 | 0,09993124 |
| 0,036272855 | 0,09993124 |
| 0,036421132 | 0,09993124 |
| 0,036421132 | 0,09993124 |
| 0,036421132 | 0,09993124 |
| 0,036644122 | 0,09993124 |
| 0,036644122 | 0,09993124 |
| 0,036644122 | 0,09993124 |
| 0,036644122 | 0,09993124 |
| 0,036644122 | 0,09993124 |
| 0,037285122 | 0,101457772 |

| | |
|---|---|
| 0,037451473 | 0,101468307 |
| 0,037451473 | 0,101468307 |
| 0,038000937 | 0,102734136 |
| 0,038627145 | 0,103976949 |
| 0,038627145 | 0,103976949 |
| 0,038993143 | 0,104736421 |
| 0,039237318 | 0,105034247 |
| 0,039272212 | 0,105034247 |
| 0,040048354 | 0,106881184 |
| 0,040908002 | 0,108480031 |
| 0,040908002 | 0,108480031 |
| 0,040908002 | 0,108480031 |
| 0,041074217 | 0,108690036 |
| 0,042094914 | 0,111155493 |
| 0,042465014 | 0,111426055 |
| 0,042465014 | 0,111426055 |
| 0,042465014 | 0,111426055 |
| 0,042962367 | 0,111667179 |
| 0,042962367 | 0,111667179 |
| 0,042962367 | 0,111667179 |
| 0,042962367 | 0,111667179 |
| 0,043003934 | 0,111667179 |
| 0,043227065 | 0,11178179 |
| 0,043227065 | 0,11178179 |
| 0,043939662 | 0,113389748 |
| 0,045385573 | 0,116879549 |
| 0,045694247 | 0,116940625 |
| 0,045694247 | 0,116940625 |
| 0,045783799 | 0,116940625 |
| 0,045783799 | 0,116940625 |
| 0,046929696 | 0,119622838 |
| 0,047826663 | 0,120192157 |
| 0,047826663 | 0,120192157 |
| 0,047826663 | 0,120192157 |
| 0,047826663 | 0,120192157 |
| 0,047826663 | 0,120192157 |
| 0,047826663 | 0,120192157 |
| 0,047826663 | 0,120192157 |
| 0,04819947 | 0,120402276 |
| 0,04819947 | 0,120402276 |
| 0,04819947 | 0,120402276 |
| 0,049227003 | 0,122723605 |
