## Supplemental Table S8 for "Temporal analysis of tear fluid proteome reveals critical corneal repair events after photorefractive surgery"

| term |
| --- |
| Cytoplasmic Translation (GO:0002181) |
| Macromolecule Biosynthetic Process (GO:0009059) |
| Translation (GO:0006412) |
| Gene Expression (GO:0010467) |
| Protein Metabolic Process (GO:0019538) |
| Protein Transport (GO:0015031) |
| RNA Processing (GO:0006396) |
| protein-RNA Complex Assembly (GO:0022618) |
| mRNA Splicing, via Spliceosome (GO:0000398) |
| Ribosomal Small Subunit Biogenesis (GO:0042274) |
| RNA Splicing, via Transesterification Reactions With Bulged Adenosine as Nucleophile (GO:0000377) |
| mRNA Processing (GO:0006397) |
| Ribonucleoprotein Complex Biogenesis (GO:0022613) |
| Ribosome Biogenesis (GO:0042254) |
| Regulation of Substrate Adhesion-Dependent Cell Spreading (GO:1900024) |
| Regulation of Translation (GO:0006417) |
| Negative Regulation of Blood Coagulation (GO:0030195) |
| Positive Regulation of Cell-Substrate Adhesion (GO:0010811) |
| Intracellular Protein Transport (GO:0006886) |
| Protein Localization (GO:0008104) |
| Positive Regulation of Substrate Adhesion-Dependent Cell Spreading (GO:1900026) |
| 7-Methylguanosine Cap Hypermethylation (GO:0036261) |
| RNA Capping (GO:0036260) |
| Positive Regulation of Protein Metabolic Process (GO:0051247) |
| Positive Regulation of Exocytosis (GO:0045921) |
| RNA Splicing (GO:0008380) |
| Formation of Cytoplasmic Translation Initiation Complex (GO:0001732) |
| Endoplasmic Reticulum to Golgi Vesicle-Mediated Transport (GO:0006888) |
| Ribosomal Small Subunit Assembly (GO:0000028) |
| Cytoplasmic Translational Initiation (GO:0002183) |
| Regulation of Nuclear-Transcribed mRNA Catabolic Process, Nonsense-Mediated Decay (GO:2000622 |
| rRNA Processing (GO:0006364) |
| Regulation of mRNA Splicing, via Spliceosome (GO:0048024) |
| Fibrinolysis (GO:0042730) |
| Phospholipid Efflux (GO:0033700) |
| Ribosome Assembly (GO:0042255) |
| Positive Regulation of Translation (GO:0045727) |
| Positive Regulation of Secretion by Cell (GO:1903532) |
| Protein Stabilization (GO:0050821) |
| Regulation of Exocytosis (GO:0017157) |
| Plasminogen Activation (GO:0031639) |
| rRNA Metabolic Process (GO:0016072) |
| Receptor-Mediated Endocytosis (GO:0006898) |
| Reverse Cholesterol Transport (GO:0043691) |
| mRNA Metabolic Process (GO:0016071) |

| |
|---|
| Zymogen Activation (GO:0031638) |
| Negative Regulation of Protein Metabolic Process (GO:0051248) |
| Regulation of Protein-Containing Complex Assembly (GO:0043254) |
| Regulation of Receptor-Mediated Endocytosis (GO:0048259) |
| Negative Regulation of Gene Expression (GO:0010629) |
| Retrograde Transport, Endosome to Golgi (GO:0042147) |
| High-Density Lipoprotein Particle Remodeling (GO:0034375) |
| Negative Regulation of RNA Splicing (GO:0033119) |
| Positive Regulation of Protein Processing (GO:0010954) |
| Homotypic Cell-Cell Adhesion (GO:0034109) |
| Negative Regulation of Cellular Component Organization (GO:0051129) |
| Cytosolic Transport (GO:0016482) |
| Bundle of His cell-Purkinje Myocyte Adhesion Involved in Cell Communication (GO:0086073) |
| Positive Regulation of CoA-transferase Activity (GO:1905920) |
| Endocytic Recycling (GO:0032456) |
| Positive Regulation of Cell-Cell Adhesion (GO:0022409) |
| U2-type Prespliceosome Assembly (GO:1903241) |
| Cholesterol Efflux (GO:0033344) |
| RNA Export From Nucleus (GO:0006405) |
| Regulation of RNA Splicing (GO:0043484) |
| Positive Regulation of Endocytosis (GO:0045807) |
| Supramolecular Fiber Organization (GO:0097435) |
| Spliceosomal snRNP Assembly (GO:0000387) |
| Cardiac Muscle Cell-Cardiac Muscle Cell Adhesion (GO:0086042) |
| - Reg of Nuclear-Transcribed mRNA Cat Proc, Nonsense-Mediated Decay (GO:2000623) |
| Blood Coagulation, Fibrin Clot Formation (GO:0072378) |
| Cellular Response to Superoxide (GO:0071451) |
| Protein Activation Cascade (GO:0072376) |
| Protein Localization to Organelle (GO:0033365) |
| Stress Granule Assembly (GO:0034063) |
| Regulation of Alternative mRNA Splicing, via Spliceosome (GO:0000381) |
| Regulation of Arp2/3 Complex-Mediated Actin Nucleation (GO:0034315) |
| mRNA Stabilization (GO:0048255) |
| DNA Geometric Change (GO:0032392) |
| Positive Regulation of Plasminogen Activation (GO:0010756) |
| Regulation of Cdc42 Protein Signal Transduction (GO:0032489) |
| Keratinocyte Differentiation (GO:0030216) |
| Early Endosome to Golgi Transport (GO:0034498) |
| Regulation of Fibrinolysis (GO:0051917) |
| Regulation of Plasminogen Activation (GO:0010755) |
| Regulation of Protein Localization (GO:0032880) |
| Negative Regulation of mRNA Catabolic Process (GO:1902373) |
| Negative Regulation of Ubiquitin-Dependent Protein Catabolic Process (GO:2000059) |
| Response to Reactive Oxygen Species (GO:0000302) |
| Protein Secretion (GO:0009306) |
| Plasma Membrane Bounded Cell Projection Organization (GO:0120036) |

| |
|---|
| Negative Regulation of Wound Healing (GO:0061045) |
| Positive Regulation of Cell Migration (GO:0030335) |
| Cholesterol Homeostasis (GO:0042632) |
| Clathrin-Dependent Endocytosis (GO:0072583) |
| Sterol Homeostasis (GO:0055092) |
| Negative Regulation of Ubiquitin Protein Ligase Activity (GO:1904667) |
| Bundle of His Cell to Purkinje Myocyte Communication (GO:0086069) |
| Spliceosomal Complex Assembly (GO:0000245) |
| Negative Regulation of Translation (GO:0017148) |
| Positive Regulation of Gene Expression (GO:0010628) |
| Regulation of mRNA Catabolic Process (GO:0061013) |
| Retrograde Vesicle-Mediated Transport, Golgi to Endoplasmic Reticulum (GO:0006890) |
| Positive Regulation of Hormone Secretion (GO:0046887) |
| Regulation of Translation in Response to Stress (GO:0043555) |
| High-Density Lipoprotein Particle Assembly (GO:0034380) |
| intra-Golgi Vesicle-Mediated Transport (GO:0006891) |
| Regulation of Cellular Localization (GO:0060341) |
| Positive Regulation of Blood Coagulation (GO:0030194) |
| Iron Ion Transport (GO:0006826) |
| Regulation of Proteasomal Ubiquitin-Dependent Protein Catabolic Process (GO:0032434) |
| Positive Regulation of Cellular Component Organization (GO:0051130) |
| Platelet Aggregation (GO:0070527) |
| Peptide Cross-Linking (GO:0018149) |
| Cholesterol Transport (GO:0030301) |
| Regulation of Cell Development (GO:0060284) |
| Negative Regulation of Cholesterol Transport (GO:0032375) |
| CRD-mediated mRNA Stabilization (GO:0070934) |
| Regulation of Low-Density Lipoprotein Particle Clearance (GO:0010988) |
| Cellular Response to Heat (GO:0034605) |
| Intermediate Filament Organization (GO:0045109) |
| Response to Calcium Ion (GO:0051592) |
| Protein-Containing Complex Assembly (GO:0065003) |
| Regulation of Ventricular Cardiac Muscle Cell Action Potential (GO:0098911) |
| - Reg of Nuclear-Transcribed mRNA Cat Proc, Deadenylation-Dep Decay (GO:1900152) |
| Negative Regulation of Ubiquitin-Protein Transferase Activity (GO:0051444) |
| Maturation of SSU-rRNA (GO:0030490) |
| Regulation of Heterotypic Cell-Cell Adhesion (GO:0034114) |
| Positive Regulation of mRNA Splicing, via Spliceosome (GO:0048026) |
| Regulation of Stress Fiber Assembly (GO:0051492) |
| Receptor Internalization (GO:0031623) |
| Epidermal Cell Differentiation (GO:0009913) |
| Positive Regulation of Transferase Activity (GO:0051347) |
| Negative Regulation of Cell-Substrate Adhesion (GO:0010812) |
| Regulation of Blood Vessel Endothelial Cell Migration (GO:0043535) |
| Negative Regulation of Apoptotic Process (GO:0043066) |
| Establishment of Protein Localization to Extracellular Region (GO:0035592) |

| |
|---|
| Regulation of Protein Secretion (GO:0050708) |
| Regulation of Protein Binding (GO:0043393) |
| Negative Regulation of Hemostasis (GO:1900047) |
| Chaperone Cofactor-Dependent Protein Refolding (GO:0051085) |
| Positive Regulation of Telomere Maintenance via Telomere Lengthening (GO:1904358) |
| Establishment of Protein Localization to Plasma Membrane (GO:0061951) |
| Regulation of mRNA Stability (GO:0043488) |
| Regulation of Apoptotic Process (GO:0042981) |
| Acylglycerol Homeostasis (GO:0055090) |
| Positive Regulation of mRNA Processing (GO:0050685) |
| Positive Regulation of Vasoconstriction (GO:0045907) |
| Positive Regulation of Heterotypic Cell-Cell Adhesion (GO:0034116) |
| Removal of Superoxide Radicals (GO:0019430) |
| Positive Regulation of Coagulation (GO:0050820) |
| Positive Regulation of Receptor-Mediated Endocytosis (GO:0048260) |
| Positive Regulation of Stress Fiber Assembly (GO:0051496) |
| RNA Biosynthetic Process (GO:0032774) |
| Negative Regulation of Protein Phosphorylation (GO:0001933) |
| Maturation of SSU-rRNA Frm Tricistronic rRNA Trnscpt (GO:0000462) |
| Negative Regulation of Proteolysis Involved in Protein Catabolic Process (GO:1903051) |
| Nuclear-Transcribed mRNA Catabolic Process, Nonsense-Mediated Decay (GO:0000184) |
| Acute-Phase Response (GO:0006953) |
| miRNA Metabolic Process (GO:0010586) |
| Negative Regulation of Coagulation (GO:0050819) |
| Phagolysosome Assembly (GO:0001845) |
| Positive Regulation of Phagocytosis (GO:0050766) |
| Hydrogen Peroxide Metabolic Process (GO:0042743) |
| Positive Regulation of Signal Transduction by P53 Class Mediator (GO:1901798) |
| Regulation of Nitric Oxide Biosynthetic Process (GO:0045428) |
| Protein Localization to Plasma Membrane (GO:0072659) |
| Regulation of Endothelial Cell Proliferation (GO:0001936) |
| Positive Regulation of Biosynthetic Process (GO:0009891) |
| Regulation of Blood Coagulation (GO:0030193) |
| Arp2/3 Complex-Mediated Actin Nucleation (GO:0034314) |
| Regulation of Vesicle Size (GO:0097494) |
| Clathrin Coat Assembly (GO:0048268) |
| Negative Regulation of mRNA Splicing, via Spliceosome (GO:0048025) |
| Positive Regulation of Cytoplasmic Translation (GO:2000767) |
| Positive Regulation of Nitric Oxide Biosynthetic Process (GO:0045429) |
| Regulation of Cell Population Proliferation (GO:0042127) |
| Regulation of Blood Vessel Branching (GO:1905553) |
| Protein Targeting to Vacuole Involved in Autophagy (GO:0071211) |
| Positive Regulation of Intrinsic Apoptotic Signaling Pathway by P53 Class Mediator (GO:1902255) |
| Peptide Antigen Assembly With MHC Class I Protein Complex (GO:0002502) |
| Pentose-Phosphate Shunt, Non-Oxidative Branch (GO:0009052) |
| Striated Muscle Hypertrophy (GO:0014897) |

| |
|---|
| Regulation of Very-Low-Density Lipoprotein Particle Remodeling (GO:0010901) |
| Regulation of Respiratory Burst Involved in Inflammatory Response (GO:0060264) |
| Regulation of Plasma Membrane Repair (GO:1905684) |
| MHC Class I Protein Complex Assembly (GO:0002397) |
| Negative Regulation of Morphogenesis of an Epithelium (GO:1905331) |
| Regulation of Protein Phosphorylation (GO:0001932) |
| Secondary Alcohol Metabolic Process (GO:1902652) |
| Positive Regulation of Peptide Hormone Secretion (GO:0090277) |
| Negative Regulation of Multicellular Organismal Process (GO:0051241) |
| Regulation of Primary Metabolic Process (GO:0080090) |
| Skin Development (GO:0043588) |
| Membrane Raft Organization (GO:0031579) |
| Regulation of Interleukin-1 Production (GO:0032652) |
| Cellular Response to Gamma Radiation (GO:0071480) |
| Negative Regulation of mRNA Processing (GO:0050686) |
| Positive Regulation of Nitric Oxide Metabolic Process (GO:1904407) |
| Positive Regulation of Lamellipodium Organization (GO:1902745) |
| Membraneless Organelle Assembly (GO:0140694) |
| RNA Methylation (GO:0001510) |
| Triglyceride Metabolic Process (GO:0006641) |
| Negative Regulation of Phosphorylation (GO:0042326) |
| Nuclear Export (GO:0051168) |
| Negative Regulation of Proteasomal Ubiquitin-Dependent Protein Catabolic Process (GO:0032435) |
| Positive Regulation of Cell Motility (GO:2000147) |
| Positive Regulation of Actin Filament Bundle Assembly (GO:0032233) |
| DNA Conformation Change (GO:0071103) |
| Regulation of Ubiquitin Protein Ligase Activity (GO:1904666) |
| Myeloid Cell Activation Involved in Immune Response (GO:0002275) |
| Positive Regulation of Cellular Process (GO:0048522) |
| Negative Regulation of Lipid Metabolic Process (GO:0045833) |
| Positive Regulation of Protein Transport (GO:0051222) |
| Heterochromatin Formation (GO:0031507) |
| Regulation of Endodeoxyribonuclease Activity (GO:0032071) |
| Humoral Immune Response Mediated by Circulating Immunoglobulin (GO:0002455) |
| Negative Regulation of Lipase Activity (GO:0060192) |
| Negative Regulation of Lipoprotein Particle Clearance (GO:0010985) |
| Positive Regulation of Membrane Depolarization (GO:1904181) |
| Positive Regulation of Respiratory Burst (GO:0060267) |
| Protein Oxidation (GO:0018158) |
| Antibacterial Humoral Response (GO:0019731) |
| Regulation of Organelle Organization (GO:0033043) |
| Regulation of Acute Inflammatory Response (GO:0002673) |
| Regulation of Amyloid Precursor Protein Catabolic Process (GO:1902991) |
| Regulation of Smooth Muscle Cell Proliferation (GO:0048660) |
| Actin Nucleation (GO:0045010) |
| Phosphatidylcholine Biosynthetic Process (GO:0006656) |

| |
|---|
| Positive Regulation of RNA Splicing (GO:0033120) |
| Negative Regulation of Smooth Muscle Cell Proliferation (GO:0048662) |
| tRNA Aminoacylation for Protein Translation (GO:0006418) |
| 'De Novo' Post-Translational Protein Folding (GO:0051084) |
| Positive Regulation of ERK1 and ERK2 Cascade (GO:0070374) |
| Protein Localization to Cytoplasmic Stress Granule (GO:1903608) |
| Positive Regulation of Leukocyte Differentiation (GO:1902107) |
| Positive Regulation of Apoptotic Cell Clearance (GO:2000427) |
| Regulation of Apoptotic Cell Clearance (GO:2000425) |
| Glyceraldehyde-3-Phosphate Metabolic Process (GO:0019682) |
| Triglyceride-Rich Lipoprotein Particle Remodeling (GO:0034370) |
| Positive Regulation of Arp2/3 Complex-Mediated Actin Nucleation (GO:2000601) |
| Superoxide Metabolic Process (GO:0006801) |
| Melanosome Organization (GO:0032438) |
| Negative Regulation of Endothelial Cell Migration (GO:0010596) |
| Regulation of Cellular Component Biogenesis (GO:0044087) |
| Myeloid Cell Differentiation (GO:0030099) |
| Negative Regulation of Response to Stimulus (GO:0048585) |
| Regulation of Cell-Cell Adhesion Mediated by Cadherin (GO:2000047) |
| Regulation of Actin Nucleation (GO:0051125) |
| Regulation of Nuclear-Transcribed mRNA Catabolic Process, Deadenylation-Dependent Decay (GO:1[illegible] |
| Hydrogen Peroxide Catabolic Process (GO:0042744) |
| Lysosomal Lumen Acidification (GO:0007042) |
| Regulation of Protein Catabolic Process (GO:0042176) |
| mRNA Transport (GO:0051028) |
| Negative Regulation of Protein-Containing Complex Assembly (GO:0031333) |
| Protein Polymerization (GO:0051258) |
| Response to Gamma Radiation (GO:0010332) |
| Positive Regulation of MAPK Cascade (GO:0043410) |
| Actin Filament Organization (GO:0007015) |
| Epithelial Cell Differentiation (GO:0030855) |
| Neuron Development (GO:0048666) |
| Protein Catabolic Process (GO:0030163) |
| Phosphatidylcholine Metabolic Process (GO:0046470) |
| Negative Regulation of Ferroptosis (GO:0110076) |
| Positive Regulation of Peptide Secretion (GO:0002793) |
| Malate-Aspartate Shuttle (GO:0043490) |
| Nucleic Acid Transport (GO:0050657) |
| Regulation of Epithelial Cell Apoptotic Process (GO:1904035) |
| Synapse Pruning (GO:0098883) |
| Regulation of Endoplasmic Reticulum Unfolded Protein Response (GO:1900101) |
| Chaperone-Mediated Protein Complex Assembly (GO:0051131) |
| Positive Regulation of Cholesterol Efflux (GO:0010875) |
| Positive Regulation of Telomere Maintenance via Telomerase (GO:0032212) |
| mRNA Export From Nucleus (GO:0006406) |
| Phospholipid Transport (GO:0015914) |

| |
|---|
| ATP Metabolic Process (GO:0046034) |
| Regulation of Phagocytosis (GO:0050764) |
| Protein Localization to Cell Periphery (GO:1990778) |
| Negative Regulation of DNA Metabolic Process (GO:0051053) |
| Positive Regulation of Amyloid Precursor Protein Catabolic Process (GO:1902993) |
| Positive Regulation of Lamellipodium Assembly (GO:0010592) |
| Acylglycerol Metabolic Process (GO:0006639) |
| Response to Hydrogen Peroxide (GO:0042542) |
| Regulation of ERK1 and ERK2 Cascade (GO:0070372) |
| Regulation of Programmed Cell Death (GO:0043067) |
| Regulation of Endosome Size (GO:0051036) |
| DNA Topological Change (GO:0006265) |
| Modification-Dependent Macromolecule Catabolic Process (GO:0043632) |
| Negative Regulation of Striated Muscle Cell Apoptotic Process (GO:0010664) |
| Positive Regulation of Cell-Cell Adhesion Mediated by Cadherin (GO:2000049) |
| Positive Regulation of Endothelial Cell Migration (GO:0010595) |
| Protein Targeting to Lysosome (GO:0006622) |
| Positive Regulation of Epidermal Growth Factor Receptor Signaling Pathway (GO:0045742) |
| Negative Regulation of Stress Fiber Assembly (GO:0051497) |
| Acute Inflammatory Response (GO:0002526) |
| Negative Regulation of Extrinsic Apoptotic Signaling Pathway via Death Domain Receptors (GO:1902[illegible] |
| ATP Biosynthetic Process (GO:0006754) |
| Negative Regulation of Receptor-Mediated Endocytosis (GO:0048261) |
| RNA Transport (GO:0050658) |
| Response to Endoplasmic Reticulum Stress (GO:0034976) |
| Protein-Containing Complex Disassembly (GO:0032984) |
| Negative Regulation of Binding (GO:0051100) |
| Regulation of Telomere Maintenance via Telomerase (GO:0032210) |
| Negative Regulation of Protein Localization (GO:1903828) |
| Regulation of Autophagosome Assembly (GO:2000785) |
| Negative Regulation of Programmed Cell Death (GO:0043069) |
| Protein Localization to Membrane (GO:0072657) |
| Positive Regulation of Cell-Matrix Adhesion (GO:0001954) |
| Positive Regulation of Response to Stimulus (GO:0048584) |
| Positive Regulation of Phospholipid Transport (GO:2001140) |
| Positive Regulation of Lipid Transport (GO:0032370) |
| Positive Regulation of Actin Nucleation (GO:0051127) |
| Oxygen Transport (GO:0015671) |
| Negative Regulation of Macromolecule Metabolic Process (GO:0010605) |
| Pinocytosis (GO:0006907) |
| Establishment of Endothelial Intestinal Barrier (GO:0090557) |
| Cytolysis by Host of Symbiont Cells (GO:0051838) |
| Regulation of Triglyceride Catabolic Process (GO:0010896) |
| Regulation of Toll-Like Receptor 9 Signaling Pathway (GO:0034163) |
| Golgi to Lysosome Transport (GO:0090160) |
| Very-Low-Density Lipoprotein Particle Assembly (GO:0034379) |

| |
|---|
| Positive Regulation of Cell Population Proliferation (GO:0008284) |
| Regulation of Cardiac Muscle Cell Contraction (GO:0086004) |
| Positive Regulation of ERBB Signaling Pathway (GO:1901186) |
| Negative Regulation of Endothelial Cell Apoptotic Process (GO:2000352) |
| Regulation of Cytoplasmic Translation (GO:2000765) |
| Sterol Transport (GO:0015918) |
| Positive Regulation of Cellular Component Biogenesis (GO:0044089) |
| Negative Regulation of Proteasomal Protein Catabolic Process (GO:1901799) |
| Regulation of Translational Initiation (GO:0006446) |
| Ribosomal Large Subunit Biogenesis (GO:0042273) |
| Positive Regulation of Endothelial Cell Proliferation (GO:0001938) |
| Negative Regulation of Inflammatory Response (GO:0050728) |
| Negative Regulation of Actin Filament Bundle Assembly (GO:0032232) |
| Striated Muscle Cell Development (GO:0055002) |
| Regulation of Lysosomal Lumen pH (GO:0035751) |
| Erythrocyte Differentiation (GO:0030218) |
| Purine Ribonucleoside Triphosphate Biosynthetic Process (GO:0009206) |
| Protein Localization to Endosome (GO:0036010) |
| Positive Regulation of Toll-Like Receptor 4 Signaling Pathway (GO:0034145) |
| Pentose-Phosphate Shunt (GO:0006098) |
| Negative Regulation of Intracellular Transport (GO:0032387) |
| Negative Regulation of Fibrinolysis (GO:0051918) |
| Negative Regulation of Epithelial Cell Apoptotic Process (GO:1904036) |
| tRNA Splicing, via Endonucleolytic Cleavage and Ligation (GO:0006388) |
| Carbon Dioxide Transport (GO:0015670) |
| Regulation of Mitochondrial Depolarization (GO:0051900) |
| RNA Splicing, via Endonucleolytic Cleavage and Ligation (GO:0000394) |
| Regulation of Ferroptosis (GO:0110075) |
| Regulation of Dendritic Cell Differentiation (GO:2001198) |
| Macrophage Activation Involved in Immune Response (GO:0002281) |
| Protein-Containing Complex Organization (GO:0043933) |
| Positive Regulation of Cholesterol Transport (GO:0032376) |
| Regulation of Cardiac Muscle Cell Action Potential (GO:0098901) |
| Establishment of Endothelial Barrier (GO:0061028) |
| Regulation of Toll-Like Receptor 4 Signaling Pathway (GO:0034143) |
| Regulation of Monocyte Chemotaxis (GO:0090025) |
| Regulation of Lipid Biosynthetic Process (GO:0046890) |
| Telomere Maintenance (GO:0000723) |
| DNA Metabolic Process (GO:0006259) |
| Translational Elongation (GO:0006414) |
| Positive Regulation of Protein Catabolic Process (GO:0045732) |
| Negative Regulation of Protein Binding (GO:0032091) |
| Positive Regulation of Blood Vessel Endothelial Cell Migration (GO:0043536) |
| tRNA Processing (GO:0008033) |
| Synaptic Vesicle Endocytosis (GO:0048488) |
| Maintenance of Blood-Brain Barrier (GO:0035633) |

| |
|---|
| Protein Methylation (GO:0006479) |
| Positive Regulation of Keratinocyte Proliferation (GO:0010838) |
| Regulation of T Cell Mediated Immune Response to Tumor Cell (GO:0002840) |
| Regulation of PERK-mediated Unfolded Protein Response (GO:1903897) |
| Primary miRNA Processing (GO:0031053) |
| Positive Regulation of Vascular Endothelial Cell Proliferation (GO:1905564) |
| Negative Regulation of Substrate Adhesion-Dependent Cell Spreading (GO:1900025) |
| NADH Oxidation (GO:0006116) |
| Low-Density Lipoprotein Particle Remodeling (GO:0034374) |
| Detection of Bacterium (GO:0016045) |
| Response to Misfolded Protein (GO:0051788) |
| Regulation of Transforming Growth Factor Beta Production (GO:0071634) |
| V(D)J Recombination (GO:0033151) |
| Morphogenesis of a Polarized Epithelium (GO:0001738) |
| Regulation of Epidermal Growth Factor Receptor Signaling Pathway (GO:0042058) |
| Regulation of Ubiquitin-Dependent Protein Catabolic Process (GO:2000058) |
| Regulation of Vasoconstriction (GO:0019229) |
| Heterotypic Cell-Cell Adhesion (GO:0034113) |
| Negative Regulation of Cell-Matrix Adhesion (GO:0001953) |
| Positive Regulation of Protein Secretion (GO:0050714) |
| Vacuolar Acidification (GO:0007035) |
| Chromosome Condensation (GO:0030261) |
| Golgi to Plasma Membrane Protein Transport (GO:0043001) |
| Regulation of Focal Adhesion Assembly (GO:0051893) |
| Cellular Response to Ionizing Radiation (GO:0071479) |
| Muscle Filament Sliding (GO:0030049) |
| ERBB2 Signaling Pathway (GO:0038128) |
| P-body Assembly (GO:0033962) |
| Ag Processing and Presentation of Endogenous Pep Ag via MHC Cls I via ER Pway (GO:0002484) |
| Ag Processing and Presentation of Endogenous Pep Ag via MHC Cls I via ER Pway, TAP-ind (GO:00024 |
| Antigen Processing and Presentation of Endogenous Peptide Antigen via MHC Class Ib (GO:0002476) |
| Antigen Processing and Presentation of Peptide Antigen via MHC Class Ib (GO:0002428) |
| Regulation of Protein Autophosphorylation (GO:0031952) |
| Cellular Response to Arsenic-Containing Substance (GO:0071243) |
| Protein Heterotetramerization (GO:0051290) |
| Negative Regulation of Cell Population Proliferation (GO:0008285) |
| Regulation of Cell Migration (GO:0030334) |
| Regulation of Extrinsic Apoptotic Signaling Pathway via Death Domain Receptors (GO:1902041) |
| Regulation of Lamellipodium Assembly (GO:0010591) |
| Purine Ribonucleoside Triphosphate Metabolic Process (GO:0009205) |
| Regulation of Cholesterol Efflux (GO:0010874) |
| Positive Regulation of Telomere Maintenance (GO:0032206) |
| Cell-Matrix Adhesion (GO:0007160) |
| Positive Regulation of Telomere Maintenance in Response to DNA Damage (GO:1904507) |
| Positive Regulation of Exosomal Secretion (GO:1903543) |
| Polarized Epithelial Cell Differentiation (GO:0030859) |

| |
|---|
| Negative Regulation of Focal Adhesion Assembly (GO:0051895) |
| Negative Regulation of Muscle Cell Apoptotic Process (GO:0010656) |
| Intracellular Potassium Ion Homeostasis (GO:0030007) |
| Telomere Maintenance in Response to DNA Damage (GO:0043247) |
| Cell-Cell Junction Maintenance (GO:0045217) |
| Regulation of Endocytic Recycling (GO:2001135) |
| Negative Regulation of Cell-Substrate Junction Organization (GO:0150118) |
| Regulation of Proteasomal Protein Catabolic Process (GO:0061136) |
| Positive Regulation of Leukocyte Chemotaxis (GO:0002690) |
| Regulation of Developmental Growth (GO:0048638) |
| Synaptic Vesicle Recycling (GO:0036465) |
| Triglyceride Homeostasis (GO:0070328) |
| Negative Regulation of Blood Vessel Endothelial Cell Migration (GO:0043537) |
| Negative Regulation of Endocytosis (GO:0045806) |
| Multicellular Organismal-Level Iron Ion Homeostasis (GO:0060586) |
| Regulation of Cholesterol Storage (GO:0010885) |
| Negative Regulation of Phagocytosis (GO:0050765) |
| mRNA Modification (GO:0016556) |
| Negative Regulation of NLRP3 Inflammasome Complex Assembly (GO:1900226) |
| Regulation of Telomere Maintenance in Response to DNA Damage (GO:1904505) |
| Actin-Myosin Filament Sliding (GO:0033275) |
| 2-Oxoglutarate Metabolic Process (GO:0006103) |
| Gas Transport (GO:0015669) |
| Sterol Metabolic Process (GO:0016125) |
| Macromolecule Modification (GO:0043412) |
| Positive Regulation of Mononuclear Cell Migration (GO:0071677) |
| Negative Regulation of Cellular Process (GO:0048523) |

| p-value | q-value |
|---|---|
| 1,0916E-57 | 2,35568E-54 |
| 1,82783E-44 | 1,97223E-41 |
| 1,56232E-41 | 1,12383E-38 |
| 7,10909E-37 | 3,83535E-34 |
| 5,09737E-26 | 2,20003E-23 |
| 1,08003E-17 | 3,88452E-15 |
| 4,29382E-16 | 1,32372E-13 |
| 1,91068E-14 | 5,12869E-12 |
| 2,13893E-14 | 5,12869E-12 |
| 8,92843E-14 | 1,92676E-11 |
| 3,32076E-13 | 6,51473E-11 |
| 8,33184E-13 | 1,49834E-10 |
| 6,68668E-12 | 1,10999E-09 |
| 3,48592E-11 | 5,3733E-09 |
| 1,68465E-10 | 2,42365E-08 |
| 4,70262E-10 | 6,34265E-08 |
| 1,41483E-09 | 1,75199E-07 |
| 1,46135E-09 | 1,75199E-07 |
| 4,64491E-09 | 5,27564E-07 |
| 5,30862E-09 | 5,52846E-07 |
| 5,37987E-09 | 5,52846E-07 |
| 4,26284E-08 | 3,99965E-06 |
| 4,26284E-08 | 3,99965E-06 |
| 7,55799E-08 | 6,79589E-06 |
| 1,90091E-07 | 1,64087E-05 |
| 2,50624E-07 | 2,01945E-05 |
| 2,52666E-07 | 2,01945E-05 |
| 4,09618E-07 | 3,15698E-05 |
| 1,10993E-06 | 8,25941E-05 |
| 1,55951E-06 | 0,00011218 |
| 1,655E-06 | 0,000115209 |
| 2,02304E-06 | 0,000136429 |
| 3,07651E-06 | 0,000201185 |
| 5,00115E-06 | 0,000308356 |
| 5,00115E-06 | 0,000308356 |
| 5,37113E-06 | 0,000321969 |
| 9,51916E-06 | 0,000555199 |
| 1,043E-05 | 0,000592313 |
| 1,26761E-05 | 0,000701414 |
| 1,51655E-05 | 0,000818179 |
| 2,00511E-05 | 0,001041632 |
| 2,02727E-05 | 0,001041632 |
| 2,11598E-05 | 0,001061927 |
| 2,5502E-05 | 0,001250755 |
| 2,6988E-05 | 0,001294226 |

| | |
|---|---|
| 3,28897E-05 | 0,001542957 |
| 4,4821E-05 | 0,002037202 |
| 4,53131E-05 | 0,002037202 |
| 5,21558E-05 | 0,002296985 |
| 6,11829E-05 | 0,002640652 |
| 8,34923E-05 | 0,003345122 |
| 8,37055E-05 | 0,003345122 |
| 8,37055E-05 | 0,003345122 |
| 8,37055E-05 | 0,003345122 |
| 9,538E-05 | 0,003742364 |
| 0,000112376 | 0,0043305 |
| 0,000118362 | 0,004481149 |
| 0,000126865 | 0,004598014 |
| 0,000126865 | 0,004598014 |
| 0,000127841 | 0,004598014 |
| 0,000164083 | 0,005804786 |
| 0,000171335 | 0,005820239 |
| 0,000171335 | 0,005820239 |
| 0,000177099 | 0,005820239 |
| 0,000178568 | 0,005820239 |
| 0,000179813 | 0,005820239 |
| 0,000180703 | 0,005820239 |
| 0,00021146 | 0,006710738 |
| 0,00024927 | 0,007479497 |
| 0,00024927 | 0,007479497 |
| 0,000256078 | 0,007479497 |
| 0,000256078 | 0,007479497 |
| 0,000256078 | 0,007479497 |
| 0,00025648 | 0,007479497 |
| 0,000260185 | 0,007486382 |
| 0,000301167 | 0,00855156 |
| 0,00031592 | 0,008853958 |
| 0,000376778 | 0,010424201 |
| 0,000428561 | 0,01136532 |
| 0,000428561 | 0,01136532 |
| 0,000428561 | 0,01136532 |
| 0,000431861 | 0,01136532 |
| 0,000448429 | 0,011384821 |
| 0,000448429 | 0,011384821 |
| 0,000448429 | 0,011384821 |
| 0,000486524 | 0,012065046 |
| 0,000491994 | 0,012065046 |
| 0,000491994 | 0,012065046 |
| 0,000517922 | 0,012418614 |
| 0,000517922 | 0,012418614 |
| 0,00053697 | 0,012619746 |

| | |
|---|---|
| 0,000538006 | 0,012619746 |
| 0,000587391 | 0,013630007 |
| 0,000602599 | 0,013834143 |
| 0,000633183 | 0,014383243 |
| 0,000655069 | 0,014725396 |
| 0,000673669 | 0,014834477 |
| 0,000673669 | 0,014834477 |
| 0,000698244 | 0,01517103 |
| 0,000703013 | 0,01517103 |
| 0,000757994 | 0,01619555 |
| 0,000800251 | 0,016766427 |
| 0,000800251 | 0,016766427 |
| 0,000902776 | 0,01873261 |
| 0,000992798 | 0,020065015 |
| 0,000992798 | 0,020065015 |
| 0,000994883 | 0,020065015 |
| 0,001012262 | 0,020226492 |
| 0,001107545 | 0,021728022 |
| 0,001107545 | 0,021728022 |
| 0,00113223 | 0,022012183 |
| 0,001287616 | 0,024809599 |
| 0,001309081 | 0,024999973 |
| 0,001342795 | 0,025269633 |
| 0,001373895 | 0,025269633 |
| 0,001373895 | 0,025269633 |
| 0,00139346 | 0,025269633 |
| 0,00139346 | 0,025269633 |
| 0,00139346 | 0,025269633 |
| 0,001491072 | 0,026814442 |
| 0,001550097 | 0,027645534 |
| 0,001859111 | 0,032016343 |
| 0,001879115 | 0,032016343 |
| 0,001882513 | 0,032016343 |
| 0,001882513 | 0,032016343 |
| 0,001882513 | 0,032016343 |
| 0,001909932 | 0,032016343 |
| 0,001913859 | 0,032016343 |
| 0,001913859 | 0,032016343 |
| 0,001966041 | 0,032636275 |
| 0,002024196 | 0,033092536 |
| 0,002024196 | 0,033092536 |
| 0,002148871 | 0,03460645 |
| 0,002148871 | 0,03460645 |
| 0,00221801 | 0,035455306 |
| 0,002244774 | 0,035619279 |
| 0,002408876 | 0,037739366 |

| | |
|---|---|
| 0,002413361 | 0,037739366 |
| 0,002463555 | 0,038014567 |
| 0,002466191 | 0,038014567 |
| 0,002633995 | 0,040029297 |
| 0,002633995 | 0,040029297 |
| 0,002690996 | 0,040609575 |
| 0,002845503 | 0,042354282 |
| 0,002845862 | 0,042354282 |
| 0,003055464 | 0,044855047 |
| 0,003055464 | 0,044855047 |
| 0,003150135 | 0,045019813 |
| 0,003150135 | 0,045019813 |
| 0,003150135 | 0,045019813 |
| 0,003150135 | 0,045019813 |
| 0,003325799 | 0,046908975 |
| 0,003325799 | 0,046908975 |
| 0,003422186 | 0,047955047 |
| 0,003501533 | 0,048086817 |
| 0,003520722 | 0,048086817 |
| 0,003520722 | 0,048086817 |
| 0,003520722 | 0,048086817 |
| 0,003939427 | 0,052477055 |
| 0,003939427 | 0,052477055 |
| 0,003939427 | 0,052477055 |
| 0,003939427 | 0,052477055 |
| 0,003990925 | 0,052731653 |
| 0,004031846 | 0,052731653 |
| 0,004031846 | 0,052731653 |
| 0,004061705 | 0,05280216 |
| 0,004195318 | 0,054212551 |
| 0,004548067 | 0,058075311 |
| 0,004548067 | 0,058075311 |
| 0,004590862 | 0,05827694 |
| 0,004838613 | 0,059667008 |
| 0,004838613 | 0,059667008 |
| 0,004838613 | 0,059667008 |
| 0,004838613 | 0,059667008 |
| 0,004838613 | 0,059667008 |
| 0,005199737 | 0,060586237 |
| 0,005281579 | 0,060586237 |
| 0,005323288 | 0,060586237 |
| 0,005323288 | 0,060586237 |
| 0,005323288 | 0,060586237 |
| 0,005323288 | 0,060586237 |
| 0,005323288 | 0,060586237 |
| 0,005323288 | 0,060586237 |

| | |
|---|---|
| 0,005323288 | 0,060586237 |
| 0,005323288 | 0,060586237 |
| 0,005323288 | 0,060586237 |
| 0,005323288 | 0,060586237 |
| 0,005323288 | 0,060586237 |
| 0,005346268 | 0,060586237 |
| 0,005373432 | 0,060586237 |
| 0,005373432 | 0,060586237 |
| 0,005390434 | 0,060586237 |
| 0,005426431 | 0,060674809 |
| 0,005782886 | 0,063018263 |
| 0,005851736 | 0,063018263 |
| 0,005851736 | 0,063018263 |
| 0,005851736 | 0,063018263 |
| 0,005851736 | 0,063018263 |
| 0,005860378 | 0,063018263 |
| 0,005860378 | 0,063018263 |
| 0,005869634 | 0,063018263 |
| 0,006398507 | 0,067355993 |
| 0,006398507 | 0,067355993 |
| 0,006398507 | 0,067355993 |
| 0,006398507 | 0,067355993 |
| 0,006574623 | 0,068562837 |
| 0,006576695 | 0,068562837 |
| 0,006959247 | 0,071412006 |
| 0,00698236 | 0,071412006 |
| 0,00698236 | 0,071412006 |
| 0,00698236 | 0,071412006 |
| 0,007289807 | 0,074204731 |
| 0,007344245 | 0,074407887 |
| 0,007591383 | 0,076196304 |
| 0,007591383 | 0,076196304 |
| 0,007860308 | 0,076407861 |
| 0,007860308 | 0,076407861 |
| 0,007860308 | 0,076407861 |
| 0,007860308 | 0,076407861 |
| 0,007860308 | 0,076407861 |
| 0,007860308 | 0,076407861 |
| 0,007860308 | 0,076407861 |
| 0,008182036 | 0,078619925 |
| 0,008186226 | 0,078619925 |
| 0,008233597 | 0,078619925 |
| 0,008233597 | 0,078619925 |
| 0,008846 | 0,084095457 |
| 0,00960813 | 0,090542993 |
| 0,00960813 | 0,090542993 |

| | |
|---|---|
| 0,010002009 | 0,092636636 |
| 0,010002009 | 0,092636636 |
| 0,010002009 | 0,092636636 |
| 0,010002009 | 0,092636636 |
| 0,010718014 | 0,095125308 |
| 0,010833018 | 0,095125308 |
| 0,010833018 | 0,095125308 |
| 0,010833018 | 0,095125308 |
| 0,010833018 | 0,095125308 |
| 0,010833018 | 0,095125308 |
| 0,010833018 | 0,095125308 |
| 0,010833018 | 0,095125308 |
| 0,011009405 | 0,095125308 |
| 0,011009405 | 0,095125308 |
| 0,011009405 | 0,095125308 |
| 0,011009405 | 0,095125308 |
| 0,011059265 | 0,095125308 |
| 0,011059265 | 0,095125308 |
| 0,011108238 | 0,095125308 |
| 0,011108238 | 0,095125308 |
| 0,011108238 | 0,095125308 |
| 0,011108238 | 0,095125308 |
| 0,011108238 | 0,095125308 |
| 0,011694586 | 0,099750658 |
| 0,012728087 | 0,106941212 |
| 0,012728087 | 0,106941212 |
| 0,012728087 | 0,106941212 |
| 0,012735816 | 0,106941212 |
| 0,012830057 | 0,107314974 |
| 0,013028776 | 0,108556368 |
| 0,013299241 | 0,110383703 |
| 0,013532141 | 0,11178131 |
| 0,013616606 | 0,11178131 |
| 0,013623023 | 0,11178131 |
| 0,014219479 | 0,113650506 |
| 0,014219479 | 0,113650506 |
| 0,014219479 | 0,113650506 |
| 0,014219479 | 0,113650506 |
| 0,014219479 | 0,113650506 |
| 0,014219479 | 0,113650506 |
| 0,014219479 | 0,113650506 |
| 0,014492401 | 0,114251419 |
| 0,014492401 | 0,114251419 |
| 0,014492401 | 0,114251419 |
| 0,014559379 | 0,114251419 |
| 0,014559379 | 0,114251419 |

| | |
|---|---|
| 0,015537924 | 0,121049965 |
| 0,015537924 | 0,121049965 |
| 0,015681785 | 0,121305244 |
| 0,015683115 | 0,121305244 |
| 0,016379186 | 0,125787488 |
| 0,016379186 | 0,125787488 |
| 0,017018574 | 0,129774143 |
| 0,017018574 | 0,129774143 |
| 0,017481837 | 0,13283734 |
| 0,017965986 | 0,13296574 |
| 0,017998543 | 0,13296574 |
| 0,017998543 | 0,13296574 |
| 0,017998543 | 0,13296574 |
| 0,017998543 | 0,13296574 |
| 0,017998543 | 0,13296574 |
| 0,018026732 | 0,13296574 |
| 0,018397049 | 0,13296574 |
| 0,018397049 | 0,13296574 |
| 0,018397049 | 0,13296574 |
| 0,018397049 | 0,13296574 |
| 0,018397049 | 0,13296574 |
| 0,018397049 | 0,13296574 |
| 0,018397049 | 0,13296574 |
| 0,018422964 | 0,13296574 |
| 0,01854806 | 0,133422376 |
| 0,019888457 | 0,140780875 |
| 0,019888457 | 0,140780875 |
| 0,019897204 | 0,140780875 |
| 0,019897204 | 0,140780875 |
| 0,019897204 | 0,140780875 |
| 0,020227826 | 0,142652447 |
| 0,020377485 | 0,143239778 |
| 0,021442134 | 0,148907544 |
| 0,021944913 | 0,148907544 |
| 0,022149825 | 0,148907544 |
| 0,022149825 | 0,148907544 |
| 0,022149825 | 0,148907544 |
| 0,022149825 | 0,148907544 |
| 0,022149825 | 0,148907544 |
| 0,022149825 | 0,148907544 |
| 0,022149825 | 0,148907544 |
| 0,022149825 | 0,148907544 |
| 0,022149825 | 0,148907544 |
| 0,022149825 | 0,148907544 |
| 0,022149825 | 0,148907544 |
| 0,022149825 | 0,148907544 |

| | |
|---|---|
| 0,022224739 | 0,148947164 |
| 0,022828013 | 0,150650921 |
| 0,022828013 | 0,150650921 |
| 0,022828013 | 0,150650921 |
| 0,022828013 | 0,150650921 |
| 0,022828013 | 0,150650921 |
| 0,023007381 | 0,151247018 |
| 0,023058512 | 0,151247018 |
| 0,023627997 | 0,154512782 |
| 0,024747019 | 0,161341593 |
| 0,024968468 | 0,161933086 |
| 0,024987821 | 0,161933086 |
| 0,025241429 | 0,162116081 |
| 0,025241429 | 0,162116081 |
| 0,025241429 | 0,162116081 |
| 0,026508257 | 0,16387079 |
| 0,026653683 | 0,16387079 |
| 0,026653683 | 0,16387079 |
| 0,026653683 | 0,16387079 |
| 0,026653683 | 0,16387079 |
| 0,026653683 | 0,16387079 |
| 0,026653683 | 0,16387079 |
| 0,026653683 | 0,16387079 |
| 0,026653683 | 0,16387079 |
| 0,026653683 | 0,16387079 |
| 0,026653683 | 0,16387079 |
| 0,026653683 | 0,16387079 |
| 0,026653683 | 0,16387079 |
| 0,026653683 | 0,16387079 |
| 0,026653683 | 0,16387079 |
| 0,027021041 | 0,165657402 |
| 0,027786584 | 0,16707042 |
| 0,027786584 | 0,16707042 |
| 0,027786584 | 0,16707042 |
| 0,027786584 | 0,16707042 |
| 0,027786584 | 0,16707042 |
| 0,027786584 | 0,16707042 |
| 0,027793457 | 0,16707042 |
| 0,028236485 | 0,169262043 |
| 0,028342751 | 0,169428411 |
| 0,030110408 | 0,178367433 |
| 0,030250949 | 0,178367433 |
| 0,030250949 | 0,178367433 |
| 0,030250949 | 0,178367433 |
| 0,030463018 | 0,178367433 |
| 0,030463018 | 0,178367433 |

| | |
|---|---|
| 0,030463018 | 0,178367433 |
| 0,031491192 | 0,178367433 |
| 0,031491192 | 0,178367433 |
| 0,031491192 | 0,178367433 |
| 0,031491192 | 0,178367433 |
| 0,031491192 | 0,178367433 |
| 0,031491192 | 0,178367433 |
| 0,031491192 | 0,178367433 |
| 0,031491192 | 0,178367433 |
| 0,031491192 | 0,178367433 |
| 0,031491192 | 0,178367433 |
| 0,031491192 | 0,178367433 |
| 0,031491192 | 0,178367433 |
| 0,031491192 | 0,178367433 |
| 0,032233224 | 0,181616967 |
| 0,032233224 | 0,181616967 |
| 0,033270054 | 0,186002009 |
| 0,033270054 | 0,186002009 |
| 0,033270054 | 0,186002009 |
| 0,034031991 | 0,189770122 |
| 0,036206809 | 0,196711479 |
| 0,036206809 | 0,196711479 |
| 0,036206809 | 0,196711479 |
| 0,036421132 | 0,196711479 |
| 0,036421132 | 0,196711479 |
| 0,036644122 | 0,196711479 |
| 0,036644122 | 0,196711479 |
| 0,036644122 | 0,196711479 |
| 0,036644122 | 0,196711479 |
| 0,036644122 | 0,196711479 |
| 0,036644122 | 0,196711479 |
| 0,036644122 | 0,196711479 |
| 0,036644122 | 0,196711479 |
| 0,036644122 | 0,196711479 |
| 0,036644122 | 0,196711479 |
| 0,036915124 | 0,1972945 |
| 0,036935578 | 0,1972945 |
| 0,039272212 | 0,207719202 |
| 0,039272212 | 0,207719202 |
| 0,039272212 | 0,207719202 |
| 0,039272212 | 0,207719202 |
| 0,040908002 | 0,215842221 |
| 0,041618558 | 0,216287679 |
| 0,042094914 | 0,216287679 |
| 0,042094914 | 0,216287679 |
| 0,042094914 | 0,216287679 |

| | |
|---|---|
| 0,042094914 | 0,216287679 |
| 0,042094914 | 0,216287679 |
| 0,042094914 | 0,216287679 |
| 0,042094914 | 0,216287679 |
| 0,042094914 | 0,216287679 |
| 0,042094914 | 0,216287679 |
| 0,042094914 | 0,216287679 |
| 0,043263719 | 0,221239588 |
| 0,043263719 | 0,221239588 |
| 0,045783799 | 0,231385103 |
| 0,045783799 | 0,231385103 |
| 0,045783799 | 0,231385103 |
| 0,045783799 | 0,231385103 |
| 0,045783799 | 0,231385103 |
| 0,047826663 | 0,236720042 |
| 0,047826663 | 0,236720042 |
| 0,047826663 | 0,236720042 |
| 0,047826663 | 0,236720042 |
| 0,047826663 | 0,236720042 |
| 0,047826663 | 0,236720042 |
| 0,047826663 | 0,236720042 |
| 0,047826663 | 0,236720042 |
| 0,047826663 | 0,236720042 |
| 0,04819947 | 0,238019351 |
| 0,048935848 | 0,241104017 |
| 0,049227003 | 0,241519062 |
| 0,049243924 | 0,241519062 |
