## Supplemental Table S9 for "Temporal analysis of tear fluid proteome reveals critical corneal repair events after photorefractive surgery"

| term |
| --- |
| mRNA Binding (GO:0003729) |
| Cadherin Binding (GO:0045296) |
| Endopeptidase Inhibitor Activity (GO:0004866) |
| GDP Binding (GO:0019003) |
| Serine-Type Endopeptidase Inhibitor Activity (GO:0004867) |
| Myosin V Binding (GO:0031489) |
| Guanyl Ribonucleotide Binding (GO:0032561) |
| Myosin Binding (GO:0017022) |
| mRNA 3'-UTR Binding (GO:0003730) |
| Poly-Pyrimidine Tract Binding (GO:0008187) |
| GTPase Activity (GO:0003924) |
| Ribonucleoside Triphosphate Phosphatase Activity (GO:0017111) |
| Lipoprotein Particle Receptor Binding (GO:0070325) |
| rRNA Binding (GO:0019843) |
| GTP Binding (GO:0005525) |
| mRNA 5'-UTR Binding (GO:0048027) |
| poly(U) RNA Binding (GO:0008266) |
| Calcium Ion Binding (GO:0005509) |
| Cadherin Binding Involved in Cell-Cell Adhesion (GO:0098641) |
| Metal Ion Binding (GO:0046872) |
| Ubiquitin Ligase Inhibitor Activity (GO:1990948) |
| Poly-Purine Tract Binding (GO:0070717) |
| Supercoiled DNA Binding (GO:0097100) |
| Phosphatidylcholine-Sterol O-acyltransferase Activator Activity (GO:0060228) |
| Cholesterol Transfer Activity (GO:0120020) |
| Cell-Cell Adhesion Mediator Activity (GO:0098632) |
| Ubiquitin-Protein Transferase Inhibitor Activity (GO:0055105) |
| Ubiquitin Protein Ligase Binding (GO:0031625) |
| Peptidase Inhibitor Activity (GO:0030414) |
| Ubiquitin-Like Protein Ligase Binding (GO:0044389) |
| Protease Binding (GO:0002020) |
| Sterol Transfer Activity (GO:0120015) |
| Protein Binding Involved in Heterotypic Cell-Cell Adhesion (GO:0086080) |
| pre-mRNA Binding (GO:0036002) |
| Endopeptidase Regulator Activity (GO:0061135) |
| DNA Polymerase Binding (GO:0070182) |
| Nucleosomal DNA Binding (GO:0031492) |
| Telomeric DNA Binding (GO:0042162) |
| Channel Inhibitor Activity (GO:0016248) |
| Ribosome Binding (GO:0043022) |
| Single-Stranded DNA Binding (GO:0003697) |
| Low-Density Lipoprotein Particle Receptor Binding (GO:0050750) |
| poly(A) Binding (GO:0008143) |
| MHC Class II Protein Complex Binding (GO:0023026) |
| Anion Binding (GO:0043168) |

| |
|---|
| Insulin-Like Growth Factor Binding (GO:0005520) |
| Apolipoprotein Receptor Binding (GO:0034190) |
| Protein Homodimerization Activity (GO:0042803) |
| GTPase Binding (GO:0051020) |
| Insulin-Like Growth Factor II Binding (GO:0031995) |
| Disordered Domain Specific Binding (GO:0097718) |
| Phosphorylation-Dependent Protein Binding (GO:0140031) |
| Small Ribosomal Subunit rRNA Binding (GO:0070181) |
| poly(G) Binding (GO:0034046) |
| aminoacyl-tRNA Ligase Activity (GO:0004812) |
| Telomerase RNA Binding (GO:0070034) |
| Double-Stranded Telomeric DNA Binding (GO:0003691) |
| G-rich Strand Telomeric DNA Binding (GO:0098505) |
| Lipase Inhibitor Activity (GO:0055102) |
| RAGE Receptor Binding (GO:0050786) |
| Myosin Heavy Chain Binding (GO:0032036) |
| Clathrin Heavy Chain Binding (GO:0032050) |
| Complement Component C3b Binding (GO:0001851) |
| Damaged DNA Binding (GO:0003684) |
| Cysteine-Type Endopeptidase Activator Activity Involved in Apoptotic Process (GO:0008656) |
| N6-methyladenosine-containing RNA Reader Activity (GO:1990247) |
| Single-Stranded Telomeric DNA Binding (GO:0043047) |
| DNA-(apurinic or Apyrimidinic Site) Endonuclease Activity (GO:0003906) |
| Phosphatidylcholine Binding (GO:0031210) |
| Histone Deacetylase Binding (GO:0042826) |
| Insulin-Like Growth Factor I Binding (GO:0031994) |
| Actin Binding (GO:0003779) |
| Hyaluronic Acid Binding (GO:0005540) |
| U6 snRNA Binding (GO:0017070) |
| Peptidase Activator Activity Involved in Apoptotic Process (GO:0016505) |
| Cholesterol Binding (GO:0015485) |
| Retinoid Binding (GO:0005501) |
| Aldehyde Dehydrogenase [NAD(P)+] Activity (GO:0004030) |
| Clathrin Adaptor Activity (GO:0035615) |
| Small GTPase Binding (GO:0031267) |

| p-value | q-value |
|---|---|
| 1,35503E-23 | 4,7426E-21 |
| 3,53626E-23 | 6,18845E-21 |
| 3,02022E-09 | 3,51174E-07 |
| 4,01342E-09 | 3,51174E-07 |
| 1,33204E-08 | 9,3243E-07 |
| 4,3334E-07 | 2,52782E-05 |
| 7,14144E-07 | 3,47534E-05 |
| 7,94364E-07 | 3,47534E-05 |
| 1,18575E-06 | 4,60397E-05 |
| 1,31542E-06 | 4,60397E-05 |
| 1,58601E-06 | 5,04638E-05 |
| 2,54375E-06 | 7,41927E-05 |
| 3,93507E-06 | 0,000105944 |
| 6,46349E-06 | 0,000161587 |
| 8,68549E-06 | 0,000188703 |
| 9,16559E-06 | 0,000188703 |
| 9,16559E-06 | 0,000188703 |
| 3,27945E-05 | 0,000637671 |
| 3,54274E-05 | 0,00065261 |
| 4,39538E-05 | 0,000769192 |
| 5,79252E-05 | 0,00096542 |
| 6,21247E-05 | 0,000988347 |
| 0,000126865 | 0,001850122 |
| 0,000126865 | 0,001850122 |
| 0,000136731 | 0,001914227 |
| 0,00014401 | 0,001938596 |
| 0,000186386 | 0,002416111 |
| 0,000234358 | 0,002929481 |
| 0,000328847 | 0,003968848 |
| 0,000458581 | 0,005350112 |
| 0,000531856 | 0,005884439 |
| 0,000538006 | 0,005884439 |
| 0,000992798 | 0,010015868 |
| 0,000994883 | 0,010015868 |
| 0,001001587 | 0,010015868 |
| 0,001107545 | 0,0107678 |
| 0,00114402 | 0,010821813 |
| 0,001309081 | 0,012057326 |
| 0,00139346 | 0,012505409 |
| 0,001622574 | 0,01419752 |
| 0,002122045 | 0,017931007 |
| 0,002254184 | 0,017931007 |
| 0,002254184 | 0,017931007 |
| 0,002254184 | 0,017931007 |
| 0,003503582 | 0,027250085 |

| | |
|---|---|
| 0,004838613 | 0,036815532 |
| 0,005323288 | 0,039641506 |
| 0,006334399 | 0,046188326 |
| 0,007523308 | 0,053737911 |
| 0,007860308 | 0,055022157 |
| 0,008233597 | 0,056505079 |
| 0,010833018 | 0,070214008 |
| 0,010833018 | 0,070214008 |
| 0,010833018 | 0,070214008 |
| 0,012079934 | 0,076872309 |
| 0,012735816 | 0,079598852 |
| 0,014219479 | 0,087312593 |
| 0,017998543 | 0,104991499 |
| 0,017998543 | 0,104991499 |
| 0,017998543 | 0,104991499 |
| 0,022149825 | 0,123054581 |
| 0,022149825 | 0,123054581 |
| 0,022149825 | 0,123054581 |
| 0,026508257 | 0,137188073 |
| 0,026653683 | 0,137188073 |
| 0,026653683 | 0,137188073 |
| 0,026653683 | 0,137188073 |
| 0,026653683 | 0,137188073 |
| 0,030463018 | 0,154522553 |
| 0,031419052 | 0,15523827 |
| 0,031491192 | 0,15523827 |
| 0,034215072 | 0,166323268 |
| 0,036644122 | 0,171005901 |
| 0,036644122 | 0,171005901 |
| 0,036644122 | 0,171005901 |
| 0,038627145 | 0,177888168 |
| 0,039272212 | 0,178510057 |
| 0,042094914 | 0,186496456 |
| 0,042094914 | 0,186496456 |
| 0,047609156 | 0,208290056 |
