## Supplemental Table S10 for "Temporal analysis of tear fluid proteome reveals critical corneal repair events after photorefractive surgery"

| term |
| --- |
| Scavenging of Heme From Plasma |
| Classical Antibody-Mediated Complement Activation |
| FCGR Activation |
| Role of LAT2 NTAL LAB on Calcium Mobilization |
| Creation of C4 and C2 Activators |
| Binding and Uptake of Ligands by Scavenger Receptors |
| Initial Triggering of Complement |
| Role of Phospholipids in Phagocytosis |
| FCERI Mediated Ca+2 Mobilization |
| FCERI Mediated MAPK Activation |
| Cell Surface Interactions at the Vascular Wall |
| FCGR3A-mediated IL10 Synthesis |
| Regulation of Complement Cascade |
| Complement Cascade |
| Parasite Infection |
| Leishmania Phagocytosis |
| FCGR3A-mediated Phagocytosis |
| Regulation of Actin Dynamics for Phagocytic Cup Formation |
| FCERI Mediated NF-kB Activation |
| Anti-inflammatory Response Favouring Leishmania Parasite Infection |
| Leishmania Parasite Growth and Survival |
| Fcgamma Receptor (FCGR) Dependent Phagocytosis |
| Immunoregulatory Interactions Between a Lymphoid and a non-Lymphoid Cell |
| Fc Epsilon Receptor (FCERI) Signaling |
| Parasitic Infection Pathways |
| Leishmania Infection |
| CD22 Mediated BCR Regulation |
| Antigen Activates B Cell Receptor (BCR) Leading to Generation of Second Messengers |
| Hemostasis |
| Signaling by the B Cell Receptor (BCR) |
| Potential Therapeutics for SARS |
| Innate Immune System |
| Vesicle-mediated Transport |
| Adaptive Immune System |
| SARS-CoV Infections |
| Immune System |
| Infectious Disease |
| Metabolism of Nucleotides |
| Interconversion of Nucleotide Di- and Triphosphates |
| Synthesis of UDP-N-acetyl-glucosamine |
| Disease |
| TP53 Regulates Transcription of Cell Death Genes With Uncertain Role in P53-Dependent Apoptosis |
| Diseases Associated With Glycosylation Precursor Biosynthesis |
| Synthesis of Substrates in N-glycan Biosythesis |
| Viral Infection Pathways |

| |
|---|
| Cytosolic Sulfonation of Small Molecules |
| Diseases of Glycosylation |
| Biosynthesis of the N-glycan Precursor (Dolichol LLO) and Transfer to a Nascent Protein |
| EPHA-mediated Growth Cone Collapse |
| Post-translational Modification Synthesis of GPI-anchored Proteins |
| Neutrophil Degranulation |
| Drug ADME |
| Phase II - Conjugation of Compounds |
| TP53 Regulates Transcription of Cell Death Genes |
| Metabolism of Vitamins and Cofactors |
| Signaling by MST1 |
| Intestinal Absorption |
| Interleukin-10 Signaling |
| Methionine Salvage Pathway |
| Metal Sequestration by Antimicrobial Proteins |
| MET Receptor Activation |
| Synthesis of GDP-mannose |
| Activation of Caspases Through Apoptosome-Mediated Cleavage |
| Molybdenum Cofactor Biosynthesis |
| EPH-ephrin Mediated Repulsion of Cells |
| Metabolism of Water-Soluble Vitamins and Cofactors |
| SMAC (DIABLO) Binds to IAPs |
| SMAC(DIABLO)-mediated Dissociation of IAP Caspase Complexes |
| SMAC, XIAP-regulated Apoptotic Response |

| p-value | q-value |
|---|---|
| 4,87281E-23 | 1,36439E-20 |
| 1,04951E-21 | 1,46931E-19 |
| 3,50512E-21 | 2,87826E-19 |
| 4,24928E-21 | 2,87826E-19 |
| 5,13974E-21 | 2,87826E-19 |
| 1,08187E-20 | 5,04875E-19 |
| 2,18292E-20 | 8,73168E-19 |
| 3,63701E-20 | 1,27295E-18 |
| 5,96511E-20 | 1,85581E-18 |
| 7,01122E-20 | 1,96314E-18 |
| 1,99017E-19 | 5,0659E-18 |
| 3,24424E-19 | 7,56989E-18 |
| 8,73983E-19 | 1,88243E-17 |
| 3,69648E-18 | 7,39296E-17 |
| 5,35945E-18 | 8,82733E-17 |
| 5,35945E-18 | 8,82733E-17 |
| 5,35945E-18 | 8,82733E-17 |
| 6,05429E-18 | 9,41779E-17 |
| 9,76847E-18 | 1,43956E-16 |
| 3,78098E-17 | 5,04131E-16 |
| 3,78098E-17 | 5,04131E-16 |
| 9,71055E-17 | 1,23589E-15 |
| 6,49771E-16 | 7,91025E-15 |
| 1,44309E-15 | 1,6836E-14 |
| 3,4048E-15 | 3,66671E-14 |
| 3,4048E-15 | 3,66671E-14 |
| 2,26838E-13 | 2,3524E-12 |
| 7,2634E-12 | 7,2634E-11 |
| 1,22485E-11 | 1,18261E-10 |
| 2,41056E-09 | 2,24985E-08 |
| 2,74445E-09 | 2,47886E-08 |
| 6,43682E-09 | 5,63222E-08 |
| 1,00184E-08 | 8,50043E-08 |
| 1,59625E-08 | 1,31456E-07 |
| 5,85275E-06 | 4,6822E-05 |
| 8,84754E-06 | 6,88142E-05 |
| 5,42878E-05 | 0,000410826 |
| 0,000255518 | 0,001882765 |
| 0,000671378 | 0,004820153 |
| 0,000897198 | 0,006280384 |
| 0,000938081 | 0,006406406 |
| 0,002850864 | 0,019005762 |
| 0,003277135 | 0,021339482 |
| 0,005980307 | 0,038056502 |
| 0,006214651 | 0,038668943 |

| | |
|---|---|
| 0,008328646 | 0,050696106 |
| 0,010083919 | 0,060074411 |
| 0,010667763 | 0,06222862 |
| 0,01202495 | 0,068714002 |
| 0,016524099 | 0,092534953 |
| 0,020543346 | 0,112786999 |
| 0,025033608 | 0,134796351 |
| 0,026234697 | 0,137403582 |
| 0,026499262 | 0,137403582 |
| 0,027534034 | 0,138930295 |
| 0,028423899 | 0,138930295 |
| 0,028423899 | 0,138930295 |
| 0,028778418 | 0,138930295 |
| 0,034011841 | 0,148801805 |
| 0,034011841 | 0,148801805 |
| 0,034011841 | 0,148801805 |
| 0,034011841 | 0,148801805 |
| 0,034011841 | 0,148801805 |
| 0,034011841 | 0,148801805 |
| 0,034802431 | 0,149918165 |
| 0,036935758 | 0,156697154 |
| 0,039567924 | 0,162926744 |
| 0,039567924 | 0,162926744 |
| 0,045092327 | 0,182983355 |
