## Supplemental Table S11 for "Temporal analysis of tear fluid proteome reveals critical corneal repair events after photorefractive surgery"

| term |
|---|
| Nucleotide Metabolic Process (GO:0009117) |
| Protein Geranylgeranylation (GO:0018344) |
| Purine Ribonucleotide Metabolic Process (GO:0009150) |
| Nucleobase-Containing Small Molecule Interconversion (GO:0015949) |
| Leukocyte Cell-Cell Adhesion (GO:0007159) |
| UDP-N-acetylglucosamine Biosynthetic Process (GO:0006048) |
| Protein Prenylation (GO:0018342) |
| UDP-N-acetylglucosamine Metabolic Process (GO:0006047) |
| Amino Sugar Biosynthetic Process (GO:0046349) |
| Podocyte Differentiation (GO:0072112) |
| Phosphate-Containing Compound Metabolic Process (GO:0006796) |
| Intestinal Absorption (GO:0050892) |
| Cellular Response to Type II Interferon (GO:0071346) |
| Negative Regulation of Phagocytosis (GO:0050765) |
| Nucleobase-Containing Small Molecule Metabolic Process (GO:0055086) |
| Response to Type II Interferon (GO:0034341) |
| Nucleotide-Sugar Biosynthetic Process (GO:0009226) |
| Regulation of Cell Adhesion (GO:0030155) |
| Regulation of Rho Protein Signal Transduction (GO:0035023) |
| Positive Regulation of Rho Protein Signal Transduction (GO:0035025) |
| Proteolysis (GO:0006508) |
| Establishment of Endothelial Barrier (GO:0061028) |
| Regulation of Interleukin-8 Production (GO:0032677) |
| Macromolecule Modification (GO:0043412) |
| Purine Ribonucleoside Triphosphate Metabolic Process (GO:0009205) |
| Negative Regulation of Endocytosis (GO:0045806) |
| Positive Regulation of DNA-binding Transcription Factor Activity (GO:0051091) |
| Detection of Chemical Stimulus Involved in Sensory Perception of Bitter Taste (GO:0001580) |
| Purine Nucleotide Metabolic Process (GO:0006163) |
| Sensory Perception of Bitter Taste (GO:0050913) |
| Positive Regulation of Neuron Apoptotic Process (GO:0043525) |
| Negative Regulation of Cell-Cell Adhesion (GO:0022408) |
| Positive Regulation of NF-kappaB Transcription Factor Activity (GO:0051092) |
| Detection of Chemical Stimulus Involved in Sensory Perception of Taste (GO:0050912) |
| Positive Regulation of Neuron Differentiation (GO:0045666) |
| Ephrin Receptor Signaling Pathway (GO:0048013) |
| Regulation of Peptidyl-Tyrosine Phosphorylation (GO:0050730) |
| Protein Metabolic Process (GO:0019538) |
| Regulation of Interleukin-6 Production (GO:0032675) |
| Positive Regulation of Establishment of Endothelial Barrier (GO:1903142) |
| Positive Regulation of Epithelial Cell Differentiation Involved in Kidney Development (GO:2000698) |
| Positive Regulation of Endothelial Cell Development (GO:1901552) |
| Positive Regulation of Cell-Cell Adhesion Mediated by Integrin (GO:0033634) |
| Regulation of Barbed-End Actin Filament Capping (GO:2000812) |
| Receptor-Mediated Virion Attachment to Host Cell (GO:0046813) |

| |
|---|
| Renal Filtration Cell Differentiation (GO:0061318) |
| Pyrimidine Nucleotide Metabolic Process (GO:0006220) |
| Inner Ear Auditory Receptor Cell Differentiation (GO:0042491) |
| Pyrimidine Ribonucleotide Metabolic Process (GO:0009218) |
| Negative Regulation of Synapse Assembly (GO:0051964) |
| Negative Regulation of Synapse Organization (GO:1905809) |
| Toll-Like Receptor 2 Signaling Pathway (GO:0034134) |
| Valine Metabolic Process (GO:0006573) |
| Negative Regulation of Mast Cell Activation (GO:0033004) |
| Epithelial Fluid Transport (GO:0042045) |
| UDP Metabolic Process (GO:0046048) |
| Hexose Import Across Plasma Membrane (GO:0140271) |
| Glomerular Epithelial Cell Differentiation (GO:0072311) |
| Regulation of Cell-Cell Adhesion (GO:0022407) |
| Protein Deglutamylation (GO:0035608) |
| Negative Regulation of Axon Regeneration (GO:0048681) |
| Cellular Response to Lipoteichoic Acid (GO:0071223) |
| Podocyte Development (GO:0072015) |
| Cellular Response to Bacterial Lipopeptide (GO:0071221) |
| Macromolecule Depalmitoylation (GO:0098734) |
| Protein Depalmitoylation (GO:0002084) |
| Inner Ear Receptor Cell Differentiation (GO:0060113) |
| Response to Lipoteichoic Acid (GO:0070391) |
| Glomerular Epithelial Cell Development (GO:0072310) |
| L-methionine Salvage (GO:0071267) |
| L-leucine Metabolic Process (GO:0006551) |
| I-kappaB Phosphorylation (GO:0007252) |
| Positive Regulation of Interleukin-18 Production (GO:0032741) |
| Epithelial Tube Formation (GO:0072175) |
| Regulation of Muscle Organ Development (GO:0048634) |
| Negative Regulation of Dendritic Spine Development (GO:0061000) |
| Regulation of Interferon-Beta Production (GO:0032648) |
| Positive Regulation of Small GTPase Mediated Signal Transduction (GO:0051057) |
| Negative Regulation of Neuron Projection Development (GO:0010977) |
| Positive Regulation of Extracellular Matrix Disassembly (GO:0090091) |
| Regulation of Cellular Extravasation (GO:0002691) |
| Pyrimidine Ribonucleotide Biosynthetic Process (GO:0009220) |
| Purine Ribonucleoside Monophosphate Catabolic Process (GO:0009169) |
| Malate Metabolic Process (GO:0006108) |
| Lymphocyte Apoptotic Process (GO:0070227) |
| Regulation of Striated Muscle Tissue Development (GO:0016202) |
| Regulation of Microvillus Organization (GO:0032530) |
| Hair Cell Differentiation (GO:0035315) |
| Positive Regulation of Plasminogen Activation (GO:0010756) |
| Negative Regulation of Neuron Projection Regeneration (GO:0070571) |
| C-terminal Protein Amino Acid Modification (GO:0018410) |

| |
|---|
| Leukotriene D4 Biosynthetic Process (GO:1901750) |
| Regulation of ERK1 and ERK2 Cascade (GO:0070372) |
| Regulation of Cytokine Production Involved in Inflammatory Response (GO:1900015) |
| Neuron Projection Guidance (GO:0097485) |
| Negative Regulation of T Cell Apoptotic Process (GO:0070233) |
| Mitral Valve Development (GO:0003174) |
| Regulation of Tumor Necrosis Factor Superfamily Cytokine Production (GO:1903555) |
| Nucleoside Triphosphate Metabolic Process (GO:0009141) |
| Integrin Activation (GO:0033622) |
| Negative Regulation of Nitric Oxide Biosynthetic Process (GO:0045019) |
| Amino Sugar Metabolic Process (GO:0006040) |
| Negative Regulation of Nitric Oxide Metabolic Process (GO:1904406) |
| Negative Regulation of Leukocyte Proliferation (GO:0070664) |
| Regulation of Cell Killing (GO:0031341) |
| Regulation of Nephron Tubule Epithelial Cell Differentiation (GO:0072182) |
| Regulation of Chemokine (C-C Motif) Ligand 5 Production (GO:0071649) |
| Regulation of Phagocytosis (GO:0050764) |
| ATP Metabolic Process (GO:0046034) |

| p-value | q-value |
|---|---|
| 5,07855E-05 | 0,030369758 |
| 0,000324068 | 0,069047613 |
| 0,000346393 | 0,069047613 |
| 0,000675436 | 0,088529193 |
| 0,000740211 | 0,088529193 |
| 0,000897198 | 0,089420711 |
| 0,001149208 | 0,095556548 |
| 0,001431117 | 0,095556548 |
| 0,001742579 | 0,095556548 |
| 0,001742579 | 0,095556548 |
| 0,001757729 | 0,095556548 |
| 0,00208325 | 0,103815305 |
| 0,002497769 | 0,114897354 |
| 0,003277135 | 0,13998047 |
| 0,004721828 | 0,186650278 |
| 0,004993987 | 0,186650278 |
| 0,007663191 | 0,23941992 |
| 0,009171057 | 0,23941992 |
| 0,009601718 | 0,23941992 |
| 0,009734268 | 0,23941992 |
| 0,010664204 | 0,23941992 |
| 0,011237488 | 0,23941992 |
| 0,012191846 | 0,23941992 |
| 0,012591366 | 0,23941992 |
| 0,01452723 | 0,23941992 |
| 0,016309109 | 0,23941992 |
| 0,018137195 | 0,23941992 |
| 0,019146811 | 0,23941992 |
| 0,020135572 | 0,23941992 |
| 0,020135572 | 0,23941992 |
| 0,021145307 | 0,23941992 |
| 0,022175749 | 0,23941992 |
| 0,022723036 | 0,23941992 |
| 0,023226628 | 0,23941992 |
| 0,024297682 | 0,23941992 |
| 0,025388646 | 0,23941992 |
| 0,025388646 | 0,23941992 |
| 0,025433836 | 0,23941992 |
| 0,028093491 | 0,23941992 |
| 0,028423899 | 0,23941992 |
| 0,028423899 | 0,23941992 |
| 0,028423899 | 0,23941992 |
| 0,028423899 | 0,23941992 |
| 0,028423899 | 0,23941992 |
| 0,028423899 | 0,23941992 |

| | |
|---|---|
| 0,028423899 | 0,23941992 |
| 0,028423899 | 0,23941992 |
| 0,028423899 | 0,23941992 |
| 0,028423899 | 0,23941992 |
| 0,028423899 | 0,23941992 |
| 0,028423899 | 0,23941992 |
| 0,028423899 | 0,23941992 |
| 0,028423899 | 0,23941992 |
| 0,028423899 | 0,23941992 |
| 0,028423899 | 0,23941992 |
| 0,028423899 | 0,23941992 |
| 0,028423899 | 0,23941992 |
| 0,028423899 | 0,23941992 |
| 0,028778418 | 0,23941992 |
| 0,034011841 | 0,23941992 |
| 0,034011841 | 0,23941992 |
| 0,034011841 | 0,23941992 |
| 0,034011841 | 0,23941992 |
| 0,034011841 | 0,23941992 |
| 0,034011841 | 0,23941992 |
| 0,034011841 | 0,23941992 |
| 0,034011841 | 0,23941992 |
| 0,034011841 | 0,23941992 |
| 0,034011841 | 0,23941992 |
| 0,034011841 | 0,23941992 |
| 0,034011841 | 0,23941992 |
| 0,034011841 | 0,23941992 |
| 0,034011841 | 0,23941992 |
| 0,034011841 | 0,23941992 |
| 0,034011841 | 0,23941992 |
| 0,034011841 | 0,23941992 |
| 0,034802431 | 0,23941992 |
| 0,037337305 | 0,23941992 |
| 0,03863062 | 0,23941992 |
| 0,039567924 | 0,23941992 |
| 0,039567924 | 0,23941992 |
| 0,039567924 | 0,23941992 |
| 0,039567924 | 0,23941992 |
| 0,039567924 | 0,23941992 |
| 0,039567924 | 0,23941992 |
| 0,039567924 | 0,23941992 |
| 0,039567924 | 0,23941992 |
| 0,039567924 | 0,23941992 |
| 0,039567924 | 0,23941992 |
| 0,039567924 | 0,23941992 |
| 0,039567924 | 0,23941992 |

| | |
|---|---|
| 0,039567924 | 0,23941992 |
| 0,039926984 | 0,23941992 |
| 0,039940873 | 0,23941992 |
| 0,04067356 | 0,23941992 |
| 0,045092327 | 0,23941992 |
| 0,045092327 | 0,23941992 |
| 0,045092327 | 0,23941992 |
| 0,045092327 | 0,23941992 |
| 0,045092327 | 0,23941992 |
| 0,045092327 | 0,23941992 |
| 0,045092327 | 0,23941992 |
| 0,045092327 | 0,23941992 |
| 0,045092327 | 0,23941992 |
| 0,045092327 | 0,23941992 |
| 0,045092327 | 0,23941992 |
| 0,045092327 | 0,23941992 |
| 0,049567497 | 0,23941992 |
| 0,049567497 | 0,23941992 |
