## Supplemental Table S12 for "Temporal analysis of tear fluid proteome reveals critical corneal repair events after photorefractive surgery"

| term |
| --- |
| Nucleotidase Activity (GO:0008252) |
| Protein Geranylgeranyltransferase Activity (GO:0004661) |
| Phospholipase Inhibitor Activity (GO:0004859) |
| Carboxypeptidase Activity (GO:0004180) |
| Lipase Inhibitor Activity (GO:0055102) |
| Transition Metal Ion Binding (GO:0046914) |
| 5'-Nucleotidase Activity (GO:0008253) |
| Calcium-Dependent Phospholipid Binding (GO:0005544) |
| Nucleoside Monophosphate Kinase Activity (GO:0050145) |
| Serine-Type Endopeptidase Inhibitor Activity (GO:0004867) |
| Metallocarboxypeptidase Activity (GO:0004181) |
| ADP Binding (GO:0043531) |
| NF-kappaB Binding (GO:0051059) |
| Actin Binding (GO:0003779) |
| Malate Dehydrogenase Activity (GO:0016615) |
| Intramolecular Oxidoreductase Activity, Interconverting Aldoses and Ketoses (GO:0016861) |
| Phosphatase Binding (GO:0019902) |
| Endopeptidase Inhibitor Activity (GO:0004866) |
| Endopeptidase Activity (GO:0004175) |
| Nucleoside Kinase Activity (GO:0019206) |
| Serine-Type Exopeptidase Activity (GO:0070008) |
| Metalloexopeptidase Activity (GO:0008235) |
| Manganese Ion Binding (GO:0030145) |
| Exopeptidase Activity (GO:0008238) |
| Sulfurtransferase Activity (GO:0016783) |
| G Protein-Coupled Glutamate Receptor Binding (GO:0035256) |
| Interleukin-1 Binding (GO:0019966) |
| Iron Ion Binding (GO:0005506) |
| PDZ Domain Binding (GO:0030165) |
| Intramolecular Phosphotransferase Activity (GO:0016868) |
| Interleukin-1 Receptor Binding (GO:0005149) |
| Guanylate Kinase Activity (GO:0004385) |
| Pyrophosphatase Activity (GO:0016462) |

| p-value | q-value |
| --- | --- |
| 6,43654E-05 | 0,007981314 |
| 0,000324068 | 0,020092198 |
| 0,000897198 | 0,027329608 |
| 0,000973134 | 0,027329608 |
| 0,001149208 | 0,027329608 |
| 0,0013224 | 0,027329608 |
| 0,00208325 | 0,03690329 |
| 0,003154939 | 0,048901557 |
| 0,004721828 | 0,061925443 |
| 0,004993987 | 0,061925443 |
| 0,008328646 | 0,093886555 |
| 0,011237488 | 0,107188343 |
| 0,011237488 | 0,107188343 |
| 0,022929379 | 0,169528877 |
| 0,028423899 | 0,169528877 |
| 0,028423899 | 0,169528877 |
| 0,029370721 | 0,169528877 |
| 0,031343454 | 0,169528877 |
| 0,033884652 | 0,169528877 |
| 0,034011841 | 0,169528877 |
| 0,034011841 | 0,169528877 |
| 0,034802431 | 0,169528877 |
| 0,036061163 | 0,169528877 |
| 0,037337305 | 0,169528877 |
| 0,039567924 | 0,169528877 |
| 0,039567924 | 0,169528877 |
| 0,039567924 | 0,169528877 |
| 0,042611255 | 0,169528877 |
| 0,043970924 | 0,169528877 |
| 0,045092327 | 0,169528877 |
| 0,045092327 | 0,169528877 |
| 0,045092327 | 0,169528877 |
| 0,045346607 | 0,169528877 |
