## Supplemental Table S13 for "Temporal analysis of tear fluid proteome reveals critical corneal repair events after photorefractive surgery"

| term |
| --- |
| Neutrophil Degranulation |
| Post-translational Protein Phosphorylation |
| Regulation of IGF Transport and Uptake by Insulin-like Growth Factor Binding Proteins (IGFBPs) |
| Innate Immune System |
| Peptide Hormone Metabolism |
| Immune System |
| Extracellular Matrix Organization |
| Platelet Degranulation |
| Platelet Activation, Signaling and Aggregation |
| Response to Elevated Platelet Cytosolic Ca2+ |
| Metabolism of Proteins |
| Synthesis, Secretion, and Inactivation of Glucagon-like Peptide-1 (GLP-1) |
| Integration of Energy Metabolism |
| Incretin Synthesis, Secretion, and Inactivation |
| Activation of Kainate Receptors Upon Glutamate Binding |
| ChREBP Activates Metabolic Gene Expression |
| TGFBR3 Regulates TGF-beta Signaling |
| Hemostasis |
| Transport of Small Molecules |
| Synthesis, Secretion, and Inactivation of Glucose-dependent Insulinotropic Polypeptide (GIP) |
| XBP1(S) Activates Chaperone Genes |
| IRE1alpha Activates Chaperones |
| Aquaporin-mediated Transport |
| Post-translational Protein Modification |
| Metabolism of Angiotensinogen to Angiotensins |
| Non-integrin membrane-ECM Interactions |
| Prostacyclin Signalling Through Prostacyclin Receptor |
| G Beta Gamma Signalling Through PLC Beta |
| Asparagine N-linked Glycosylation |
| Collagen Biosynthesis and Modifying Enzymes |
| Presynaptic Function of Kainate Receptors |
| Other Interleukin Signaling |
| Thromboxane Signalling Through TP Receptor |
| ECM Proteoglycans |
| Regulation of Insulin Secretion |
| Calnexin Calreticulin Cycle |
| Surfactant Metabolism |
| Disease |
| Collagen Formation |
| G-protein Beta Gamma Signalling |
| Thrombin Signalling Through Proteinase Activated Receptors (PARs) |
| Glucagon-type Ligand Receptors |
| Signal Amplification |
| Glucagon Signaling in Metabolic Regulation |
| Class B 2 (Secretin Family Receptors) |

| |
|---|
| Unfolded Protein Response (UPR) |
| N-glycan Trimming in the ER and Calnexin Calreticulin Cycle |
| Nucleotide Catabolism |
| Transport to the Golgi and Subsequent Modification |
| Molecules Associated With Elastic Fibres |
| Fatty acyl-CoA Biosynthesis |
| Infectious Disease |
| Glucagon-like Peptide-1 (GLP1) Regulates Insulin Secretion |
| ADORA2B Mediated Anti-Inflammatory Cytokines Production |
| Vasopressin Regulates Renal Water Homeostasis via Aquaporins |
| Axon Guidance |
| Elastic Fibre Formation |
| GPER1 Signaling |
| SRP-dependent Cotranslational Protein Targeting to Membrane |
| L1CAM Interactions |
| G Alpha (Z) Signalling Events |
| Sodium-coupled Sulphate, Di- and Tri-Carboxylate Transporters |
| Signaling by TGFBR3 |
| Nervous System Development |
| Mineralocorticoid Biosynthesis |
| Maturation of Spike Protein 9683686 |
| Signaling by LRP5 Mutants |
| Uptake and Function of Diphtheria Toxin |
| SARS-CoV-2 Targets PDZ Proteins in Cell-Cell Junction |
| Acrosome Reaction and Sperm Oocyte Membrane Binding |
| Fibronectin Matrix Formation |
| Sensory Processing of Sound by Outer Hair Cells of the Cochlea |
| Neurofascin Interactions |
| Phosphate Bond Hydrolysis by NUDT Proteins |
| Parasitic Infection Pathways |
| Leishmania Infection |
| Cellular Responses to Stimuli |
| Diseases of Glycosylation |
| Signal Transduction |
| FGFR1b Ligand Binding and Activation |
| Phosphate Bond Hydrolysis by NTPDase Proteins |
| Erythrocytes Take up Oxygen and Release Carbon Dioxide |
| Fatty Acids Bound to GPR40 (FFAR1) Regulate Insulin Secretion |
| Ca2+ Pathway |
| ER to Golgi Anterograde Transport |
| NR1H2 & NR1H3 Regulate Gene Expression Linked to Lipogenesis |
| CD163 Mediating an Anti-Inflammatory Response |
| Serine Biosynthesis |
| G Alpha (S) Signalling Events |
| Anti-inflammatory Response Favouring Leishmania Parasite Infection |
| Leishmania Parasite Growth and Survival |

| |
|---|
| Sensory Processing of Sound by Inner Hair Cells of the Cochlea |
| High Laminar Flow Shear Stress Activates Signaling by PIEZO1 and PECAM1 CDH5 KDR in Endothelial [illegible] |
| Thyroxine Biosynthesis |
| Glycoprotein Hormones |
| Ionotropic Activity of Kainate Receptors |
| Activation of Ca-permeable Kainate Receptor |
| Acetylcholine Regulates Insulin Secretion |
| Reactions Specific to the Complex N-glycan Synthesis Pathway |

| p-value | q-value |
|---|---|
| 2,53678E-09 | 8,67524E-07 |
| 3,86425E-09 | 8,67524E-07 |
| 1,42401E-08 | 2,13127E-06 |
| 5,99158E-08 | 6,72555E-06 |
| 5,12719E-06 | 0,000460421 |
| 6,78279E-06 | 0,000507578 |
| 2,30495E-05 | 0,001478463 |
| 4,97813E-05 | 0,002768411 |
| 5,90998E-05 | 0,002768411 |
| 6,16573E-05 | 0,002768411 |
| 8,93086E-05 | 0,003645416 |
| 0,000160215 | 0,005994722 |
| 0,000229879 | 0,007733129 |
| 0,000241122 | 0,007733129 |
| 0,000473053 | 0,014160044 |
| 0,000706866 | 0,018669584 |
| 0,000706866 | 0,018669584 |
| 0,000957613 | 0,023887121 |
| 0,001159977 | 0,027412085 |
| 0,001936634 | 0,042789641 |
| 0,002001297 | 0,042789641 |
| 0,002245505 | 0,045828723 |
| 0,002374229 | 0,046349078 |
| 0,002818249 | 0,052724733 |
| 0,003332117 | 0,058779445 |
| 0,00340371 | 0,058779445 |
| 0,00416193 | 0,069211347 |
| 0,004609053 | 0,072894953 |
| 0,00482032 | 0,072894953 |
| 0,004870487 | 0,072894953 |
| 0,005077354 | 0,073539735 |
| 0,006607095 | 0,089896538 |
| 0,006607095 | 0,089896538 |
| 0,006915482 | 0,091325046 |
| 0,00742909 | 0,095304615 |
| 0,007728761 | 0,096394819 |
| 0,010208747 | 0,123193774 |
| 0,010816641 | 0,123193774 |
| 0,010980135 | 0,123193774 |
| 0,011563694 | 0,123193774 |
| 0,011563694 | 0,123193774 |
| 0,01226918 | 0,123193774 |
| 0,01226918 | 0,123193774 |
| 0,01226918 | 0,123193774 |
| 0,012346815 | 0,123193774 |

| | |
|---|---|
| 0,012703038 | 0,123992695 |
| 0,013735158 | 0,128480957 |
| 0,013735158 | 0,128480957 |
| 0,015189106 | 0,134463508 |
| 0,015273138 | 0,134463508 |
| 0,015273138 | 0,134463508 |
| 0,015803112 | 0,136453798 |
| 0,01942274 | 0,164543589 |
| 0,020303339 | 0,165749074 |
| 0,020303339 | 0,165749074 |
| 0,020777656 | 0,16659228 |
| 0,021200387 | 0,166999537 |
| 0,022113697 | 0,171190514 |
| 0,023030662 | 0,17526724 |
| 0,023535757 | 0,176125913 |
| 0,024949349 | 0,182457317 |
| 0,025243551 | 0,182457317 |
| 0,025925864 | 0,182457317 |
| 0,026007279 | 0,182457317 |
| 0,030216035 | 0,191084504 |
| 0,030216035 | 0,191084504 |
| 0,030216035 | 0,191084504 |
| 0,030216035 | 0,191084504 |
| 0,030216035 | 0,191084504 |
| 0,030216035 | 0,191084504 |
| 0,030216035 | 0,191084504 |
| 0,03210105 | 0,200185714 |
| 0,035163402 | 0,213356318 |
| 0,035163402 | 0,213356318 |
| 0,036804559 | 0,216849567 |
| 0,036804559 | 0,216849567 |
| 0,037292063 | 0,216849567 |
| 0,038763518 | 0,216849567 |
| 0,039876647 | 0,216849567 |
| 0,040085777 | 0,216849567 |
| 0,040085777 | 0,216849567 |
| 0,040085777 | 0,216849567 |
| 0,040085777 | 0,216849567 |
| 0,041129164 | 0,219845174 |
| 0,044960401 | 0,22611482 |
| 0,044983286 | 0,22611482 |
| 0,044983286 | 0,22611482 |
| 0,044983286 | 0,22611482 |
| 0,04712923 | 0,22611482 |
| 0,047863512 | 0,22611482 |
| 0,047863512 | 0,22611482 |

| | |
|---|---|
| 0,048449712 | 0,22611482 |
| 0,048449712 | 0,22611482 |
| 0,049856052 | 0,22611482 |
| 0,049856052 | 0,22611482 |
| 0,049856052 | 0,22611482 |
| 0,049856052 | 0,22611482 |
| 0,049856052 | 0,22611482 |
| 0,049856052 | 0,22611482 |
