## Supplemental Table S14 for "Temporal analysis of tear fluid proteome reveals critical corneal repair events after photorefractive surgery"

| term |
| --- |
| Positive Regulation of Cell Adhesion (GO:0045785) |
| Negative Regulation of Platelet Aggregation (GO:0090331) |
| Negative Regulation of Homotypic Cell-Cell Adhesion (GO:0034111) |
| Establishment or Maintenance of Epithelial Cell Apical/Basal Polarity (GO:0045197) |
| Establishment of Apical/Basal Cell Polarity (GO:0035089) |
| Negative Regulation of Platelet Activation (GO:0010544) |
| Negative Regulation of Protein Processing (GO:0010955) |
| Regulation of Plasminogen Activation (GO:0010755) |
| + Reg of Transmembrane Receptor Prot Serine/Threonine Kinase Sgnlng Pway (GO:0090100) |
| Endothelial Cell Migration (GO:0043542) |
| Cellular Component Assembly (GO:0022607) |
| Regulation of Phosphatidylinositol 3-Kinase/Protein Kinase B Signal Transduction (GO:0051896) |
| Regulation of ERK1 and ERK2 Cascade (GO:0070372) |
| Peptide Cross-Linking (GO:0018149) |
| Regulation of Cell Migration (GO:0030334) |
| Negative Regulation of T Cell Receptor Signaling Pathway (GO:0050860) |
| Positive Regulation of Transforming Growth Factor Beta Receptor Signaling Pathway (GO:0030511) |
| Response to Ketone (GO:1901654) |
| Positive Regulation of Cellular Response to Transforming Growth Factor Beta Stimulus (GO:1903846) |
| Glial Cell Migration (GO:0008347) |
| Regulation of Platelet Aggregation (GO:0090330) |
| Extracellular Matrix Organization (GO:0030198) |
| Negative Regulation of Plasminogen Activation (GO:0010757) |
| Response to Catecholamine (GO:0071869) |
| Gland Morphogenesis (GO:0022612) |
| Extrinsic Apoptotic Signaling Pathway (GO:0097191) |
| Protein-Containing Complex Organization (GO:0043933) |
| Negative Regulation of Antigen Receptor-Mediated Signaling Pathway (GO:0050858) |
| Positive Regulation of Protein Polymerization (GO:0032273) |
| Negative Regulation of Epithelial Cell Proliferation (GO:0050680) |
| Cellular Response to Prostaglandin E Stimulus (GO:0071380) |
| Adenylate Cyclase-Activating Dopamine Receptor Signaling Pathway (GO:0007191) |
| Cellular Response to Monoamine Stimulus (GO:0071868) |
| Positive Regulation of MAPK Cascade (GO:0043410) |
| Regulation of Cell Adhesion Mediated by Integrin (GO:0033628) |
| Positive Regulation of Growth (GO:0045927) |
| Positive Regulation of Cell Differentiation (GO:0045597) |
| Regulation of Sprouting Angiogenesis (GO:1903670) |
| Positive Regulation of Protein Kinase Activity (GO:0045860) |
| Regulation of T Cell Receptor Signaling Pathway (GO:0050856) |
| Cellular Response to Prostaglandin Stimulus (GO:0071379) |
| Establishment of Epithelial Cell Apical/Basal Polarity (GO:0045198) |
| Positive Regulation of Actin Filament Polymerization (GO:0030838) |
| Positive Regulation of Cellular Process (GO:0048522) |
| Regulation of Epithelial Cell Migration (GO:0010632) |

| |
|---|
| Neutrophil Chemotaxis (GO:0030593) |
| Negative Regulation of Platelet-Derived Growth Factor Receptor Signaling Pathway (GO:0010642) |
| Collagen Fibril Organization (GO:0030199) |
| Positive Regulation of Supramolecular Fiber Organization (GO:1902905) |
| Negative Regulation of Vascular Permeability (GO:0043116) |
| Cellular Response to Catecholamine Stimulus (GO:0071870) |
| Regulation of Protein Secretion (GO:0050708) |
| - Reg of Phosphatidylinositol 3-Kinase/Prot Kinase B Signal Transduction (GO:0051898) |
| Positive Regulation of Protein Phosphorylation (GO:0001934) |
| Regulation of Sensory Perception (GO:0051931) |
| Response to Progesterone (GO:0032570) |
| Substrate-Dependent Cell Migration (GO:0006929) |
| Positive Regulation of Signal Transduction (GO:0009967) |
| Positive Regulation of Developmental Growth (GO:0048639) |
| Negative Regulation of ERK1 and ERK2 Cascade (GO:0070373) |
| Neutrophil Migration (GO:1990266) |
| Microvillus Assembly (GO:0030033) |
| Acidic Amino Acid Transport (GO:0015800) |
| acyl-CoA Biosynthetic Process (GO:0071616) |
| Regulation of Platelet-Derived Growth Factor Receptor Signaling Pathway (GO:0010640) |
| Polarized Epithelial Cell Differentiation (GO:0030859) |
| Response to Prostaglandin E (GO:0034695) |
| Cotranslational Protein Targeting to Membrane (GO:0006613) |
| Negative Regulation of Cell Migration (GO:0030336) |
| Regulation of Sensory Perception of Pain (GO:0051930) |
| Regulation of Interleukin-1 Production (GO:0032652) |
| Regulation of Multicellular Organism Growth (GO:0040014) |
| One-Carbon Compound Transport (GO:0019755) |
| Peptide Catabolic Process (GO:0043171) |
| Establishment of Protein Localization to Endoplasmic Reticulum (GO:0072599) |
| Negative Regulation of Cell Population Proliferation (GO:0008285) |
| Regulation of Transforming Growth Factor Beta Receptor Signaling Pathway (GO:0017015) |
| G Protein-Coupled Acetylcholine Receptor Signaling Pathway (GO:0007213) |
| Positive Regulation of Cell Adhesion Mediated by Integrin (GO:0033630) |
| Gliogenesis (GO:0042063) |
| Positive Regulation of Cell Population Proliferation (GO:0008284) |
| Protein Localization to Membrane (GO:0072657) |
| G Protein-Coupled Dopamine Receptor Signaling Pathway (GO:0007212) |
| Establishment of Epithelial Cell Polarity (GO:0090162) |
| Positive Regulation of Angiogenesis (GO:0045766) |
| Negative Regulation of MAPK Cascade (GO:0043409) |
| Granulocyte Chemotaxis (GO:0071621) |
| Protein Targeting to Membrane (GO:0006612) |
| Regulation of Angiogenesis (GO:0045765) |
| Protein Stabilization (GO:0050821) |
| Protein Transport (GO:0015031) |

| |
|---|
| Protein Targeting (GO:0006605) |
| Receptor-Mediated Endocytosis (GO:0006898) |
| Positive Regulation of Apoptotic Signaling Pathway (GO:2001235) |
| Modulation by Host of Viral Process (GO:0044788) |
| Regulation of Cell Growth (GO:0001558) |
| Regulation of Actin Filament Polymerization (GO:0030833) |
| Negative Regulation of Cellular Process (GO:0048523) |
| Regulation of Cell Population Proliferation (GO:0042127) |
| Monocarboxylic Acid Transport (GO:0015718) |
| Regulation of Cell Adhesion (GO:0030155) |
| Positive Regulation of Phosphate Metabolic Process (GO:0045937) |
| Protein Targeting to ER (GO:0045047) |
| Positive Regulation of Cell-Substrate Adhesion (GO:0010811) |
| Protein-Containing Complex Assembly (GO:0065003) |
| Positive Regulation of Cytoskeleton Organization (GO:0051495) |
| Intracellular Protein Transport (GO:0006886) |
| Plasma Membrane Organization (GO:0007009) |
| Gland Development (GO:0048732) |
| Positive Regulation of Protein Serine/Threonine Kinase Activity (GO:0071902) |
| Negative Regulation of Supramolecular Fiber Organization (GO:1902904) |
| Establishment or Maintenance of Apical/Basal Cell Polarity (GO:0035088) |
| Regulation of Cell Size (GO:0008361) |
| Supramolecular Fiber Organization (GO:0097435) |
| Response to Tumor Necrosis Factor (GO:0034612) |
| Phagocytosis, Engulfment (GO:0006911) |
| Plasma Membrane Invagination (GO:0099024) |
| Dicarboxylic Acid Transport (GO:0006835) |
| Positive Regulation of Cell Migration (GO:0030335) |
| Negative Regulation of Protein Phosphorylation (GO:0001933) |
| Regulation of Protein Kinase Activity (GO:0045859) |
| Membrane Fusion (GO:0061025) |
| Negative Regulation of Protein Polymerization (GO:0032272) |
| Positive Regulation of Cell Motility (GO:2000147) |
| Skeletal System Development (GO:0001501) |
| + Reg of Phosphatidylinositol 3-Kinase/Prot Kinase B Signal Transduction (GO:0051897) |
| Regulation of Blood Coagulation (GO:0030193) |
| Cellular Response to Tumor Necrosis Factor (GO:0071356) |
| Peptide Hormone Processing (GO:0016486) |
| Regulation of Chemotaxis (GO:0050920) |
| Regulation of Stress-Activated MAPK Cascade (GO:0032872) |
| Morphogenesis of an Epithelium (GO:0002009) |
| Negative Regulation of Apoptotic Process (GO:0043066) |
| Regulation of Protein Localization to Plasma Membrane (GO:1903076) |
| Regulation of Extrinsic Apoptotic Signaling Pathway via Death Domain Receptors (GO:1902041) |
| Response to Steroid Hormone (GO:0048545) |
| Positive Regulation of Locomotion (GO:0040017) |

| |
|---|
| Regulation of Cell Migration Involved in Sprouting Angiogenesis (GO:0090049) |
| Protein Localization (GO:0008104) |
| Regulation of Epithelial Cell Proliferation (GO:0050678) |
| Negative Regulation of Developmental Growth (GO:0048640) |
| Actin Filament Bundle Assembly (GO:0051017) |
| Positive Regulation of Phosphorylation (GO:0042327) |
| Negative Regulation of Microtubule Polymerization or Depolymerization (GO:0031111) |
| Actin Filament Bundle Organization (GO:0061572) |
| Membrane Protein Proteolysis (GO:0033619) |
| Negative Regulation of Programmed Cell Death (GO:0043069) |
| Acetylcholine Receptor Signaling Pathway (GO:0095500) |
| Platelet Aggregation (GO:0070527) |
| Positive Regulation of Substrate Adhesion-Dependent Cell Spreading (GO:1900026) |
| Negative Regulation of Signal Transduction (GO:0009968) |
| Plasma Membrane Bounded Cell Projection Assembly (GO:0120031) |
| Positive Regulation of Hydrolase Activity (GO:0051345) |
| Cellular Response to Heat (GO:0034605) |
| Positive Regulation of Epithelial Cell Migration (GO:0010634) |
| Small GTPase-mediated Signal Transduction (GO:0007264) |
| Positive Regulation of Reproductive Process (GO:2000243) |
| Positive Regulation of SMAD Protein Signal Transduction (GO:0060391) |
| Positive Regulation of Cell Growth (GO:0030307) |
| Protein Catabolic Process (GO:0030163) |
| Regulation of Apoptotic Signaling Pathway (GO:2001233) |
| Positive Regulation of Multicellular Organismal Process (GO:0051240) |
| Negative Regulation of Intracellular Signal Transduction (GO:1902532) |
| Positive Regulation of Vasculature Development (GO:1904018) |
| Odontogenesis (GO:0042476) |
| Keratinocyte Differentiation (GO:0030216) |
| Regulation of Cellular Component Size (GO:0032535) |
| Negative Regulation of Protein Modification Process (GO:0031400) |
| Bone Development (GO:0060348) |
| Negative Regulation of Canonical Wnt Signaling Pathway (GO:0090090) |
| Cellular Response to Ketone (GO:1901655) |
| Positive Regulation of Cell-Matrix Adhesion (GO:0001954) |
| Positive Regulation of Apoptotic Process (GO:0043065) |
| Regulation of Signal Transduction (GO:0009966) |
| Positive Regulation of Chemotaxis (GO:0050921) |
| Homotypic Cell-Cell Adhesion (GO:0034109) |
| Regulation of Cell-Cell Adhesion (GO:0022407) |
| Monoatomic Anion Transport (GO:0006820) |
| Negative Regulation of Cell Growth (GO:0030308) |
| Proteolysis (GO:0006508) |
| Regulation of Canonical Wnt Signaling Pathway (GO:0060828) |
| Epithelial Tube Branching Involved in Lung Morphogenesis (GO:0060441) |
| Iodide Transport (GO:0015705) |

| |
|---|
| Prevention of Polyspermy (GO:0060468) |
| Receptor-Mediated Virion Attachment to Host Cell (GO:0046813) |
| Regulation of Fc Receptor Mediated Stimulatory Signaling Pathway (GO:0060368) |
| Regulation of Blood Vessel Remodeling (GO:0060312) |
| Regulation of Basement Membrane Organization (GO:0110011) |
| Dopamine Biosynthetic Process (GO:0042416) |
| Regulation of cGMP-mediated Signaling (GO:0010752) |
| Negative Regulation of Cellular Extravasation (GO:0002692) |
| Negative Regulation of IRE1-mediated Unfolded Protein Response (GO:1903895) |
| Chronic Inflammatory Response (GO:0002544) |
| Regulation of Transforming Growth Factor Beta2 Production (GO:0032909) |
| Response to Glucagon (GO:0033762) |
| Cell-Cell Adhesion Mediated by Integrin (GO:0033631) |
| Peptidyl-Proline Modification (GO:0018208) |
| Bradykinin Catabolic Process (GO:0010815) |
| Neutrophil Degranulation (GO:0043312) |
| Regulation of Plasma Membrane Repair (GO:1905684) |
| Regulation of Monoatomic Anion Transport (GO:0044070) |
| Regulation of Long-Chain Fatty Acid Import Across Plasma Membrane (GO:0010746) |
| Regulation of Extracellular Exosome Assembly (GO:1903551) |
| Reg of Epithelial to Mesenchymal Transition Inv in Endocardial Cushion Formation (GO:1905005) |
| Response to Lithium Ion (GO:0010226) |
| Negative Regulation of Small GTPase Mediated Signal Transduction (GO:0051058) |
| Activation of Protein Kinase Activity (GO:0032147) |
| Positive Regulation of Epithelial to Mesenchymal Transition (GO:0010718) |
| Positive Regulation of Blood Vessel Endothelial Cell Migration (GO:0043536) |
| Regulation of Interleukin-12 Production (GO:0032655) |
| Regulation of Epidermal Growth Factor Receptor Signaling Pathway (GO:0042058) |
| Organic Hydroxy Compound Transport (GO:0015850) |
| Inorganic Anion Transport (GO:0015698) |
| Regulation of Substrate Adhesion-Dependent Cell Spreading (GO:1900024) |
| Positive Regulation of Proteolysis (GO:0045862) |
| Positive Regulation of Reactive Oxygen Species Metabolic Process (GO:2000379) |
| Antibacterial Humoral Response (GO:0019731) |
| Cytoplasmic Microtubule Organization (GO:0031122) |
| B Cell Differentiation (GO:0030183) |
| Regulation of Focal Adhesion Assembly (GO:0051893) |
| Protein Localization to Plasma Membrane (GO:0072659) |
| Positive Regulation of Blood Circulation (GO:1903524) |
| - Reg of Blood Vessel Endothelial Cell Proliferation Inv in Sprouting Angiogenesis (GO:1903588) |
| Lysosomal Protein Catabolic Process (GO:1905146) |
| Positive Regulation of Secondary Metabolite Biosynthetic Process (GO:1900378) |
| Positive Regulation of Multicellular Organism Growth (GO:0040018) |
| Positive Regulation of Melanin Biosynthetic Process (GO:0048023) |
| Engulfment of Apoptotic Cell (GO:0043652) |
| Substrate-Dependent Cell Migration, Cell Extension (GO:0006930) |

| |
|---|
| Cortical Microtubule Organization (GO:0043622) |
| Nucleoside Diphosphate Metabolic Process (GO:0009132) |
| Regulation of Transforming Growth Factor Beta1 Production (GO:0032908) |
| Pharyngeal Arch Artery Morphogenesis (GO:0061626) |
| Cellular Hypotonic Response (GO:0071476) |
| Calcium Ion-Regulated Exocytosis of Neurotransmitter (GO:0048791) |
| Neurotransmitter Receptor Transport to Postsynaptic Membrane (GO:0098969) |
| Negative Regulation of Ruffle Assembly (GO:1900028) |
| Regulation of Leukocyte Degranulation (GO:0043300) |
| Negative Regulation of Granulocyte Differentiation (GO:0030853) |
| Cleavage Furrow Formation (GO:0036089) |
| Positive Regulation of Protein Localization to Plasma Membrane (GO:1903078) |
| Protein Localization to Cell Periphery (GO:1990778) |
| Receptor Internalization (GO:0031623) |
| Epidermal Cell Differentiation (GO:0009913) |
| Cellular Response to Alcohol (GO:0097306) |
| Regulation of Cytoskeleton Organization (GO:0051493) |
| Negative Regulation of Cell Motility (GO:2000146) |
| Cartilage Development (GO:0051216) |
| Positive Regulation of Protein Localization to Cell Periphery (GO:1904377) |
| Positive Regulation of Fibroblast Migration (GO:0010763) |
| Regulation of Angiotensin Levels in Blood (GO:0002002) |
| Regulation of CD4-positive, Alpha-Beta T Cell Activation (GO:2000514) |
| Protein Retention in ER Lumen (GO:0006621) |
| Hypotonic Response (GO:0006971) |
| Tight Junction Organization (GO:0120193) |
| Establishment of Centrosome Localization (GO:0051660) |
| Establishment of Golgi Localization (GO:0051683) |
| Positive Regulation of Astrocyte Differentiation (GO:0048711) |
| Cellular Response to Thyroid Hormone Stimulus (GO:0097067) |
| Peptidyl-Lysine Hydroxylation (GO:0017185) |
| Angiotensin Maturation (GO:0002003) |
| Reg of Natural Killer Cell Mediated Cytotoxicity Directed Against Tumor Cell Target (GO:0002858) |
| Regulation of Microvillus Organization (GO:0032530) |
| Negative Regulation of p38MAPK Cascade (GO:1903753) |
| Positive Regulation of Cardiac Epithelial to Mesenchymal Transition (GO:0062043) |
| Regulation of SMAD Protein Signal Transduction (GO:0060390) |
| Negative Regulation of Wnt Signaling Pathway (GO:0030178) |
| Vascular Endothelial Cell Response to Laminar Fluid Shear Stress (GO:0097700) |
| Ventricular Trabecula Myocardium Morphogenesis (GO:0003222) |
| Sphingosine-1-Phosphate Receptor Signaling Pathway (GO:0003376) |
| Positive Regulation of Early Endosome to Late Endosome Transport (GO:2000643) |
| Interleukin-12-Mediated Signaling Pathway (GO:0035722) |
| Response to Thyroid Hormone (GO:0097066) |
| Protein Catabolic Process in the Vacuole (GO:0007039) |
| Regulation of CD4-positive, Alpha-Beta T Cell Proliferation (GO:2000561) |

| |
|---|
| Multi-Pass Transmembrane Protein Insertion Into ER Membrane (GO:0160063) |
| Regulation of Cell Killing (GO:0031341) |
| Positive Regulation of Transforming Growth Factor Beta Production (GO:0071636) |
| Response to Magnesium Ion (GO:0032026) |
| Regulation of Response to Wounding (GO:1903034) |
| Cardioblast Differentiation (GO:0010002) |
| Positive Regulation by Host of Viral Genome Replication (GO:0044829) |
| Regulation of Myeloid Leukocyte Mediated Immunity (GO:0002886) |
| Negative Regulation of Retrograde Protein Transport, ER to Cytosol (GO:1904153) |
| Acid Secretion (GO:0046717) |
| T Cell Mediated Cytotoxicity (GO:0001913) |
| T Cell Apoptotic Process (GO:0070231) |
| R-loop Processing (GO:0062176) |
| Negative Regulation of Leukocyte Degranulation (GO:0043301) |
| Regulation of Fertilization (GO:0080154) |
| Regulation of Endothelial Cell Differentiation (GO:0045601) |
| Regulation of Dendritic Cell Antigen Processing and Presentation (GO:0002604) |
| Monocarboxylic Acid Biosynthetic Process (GO:0072330) |
| Regulation of Cell-Matrix Adhesion (GO:0001952) |
| Regulation of Phosphorylation (GO:0042325) |
| Adenylate Cyclase-Activating G Protein-Coupled Receptor Signaling Pathway (GO:0007189) |
| Positive Regulation of ERK1 and ERK2 Cascade (GO:0070374) |
| Regulation of MAPK Cascade (GO:0043408) |
| Glucosamine-Containing Compound Catabolic Process (GO:1901072) |
| Regulation of Cell-Cell Adhesion Mediated by Integrin (GO:0033632) |
| Morphogenesis of an Epithelial Sheet (GO:0002011) |
| Protein Localization to Microtubule Cytoskeleton (GO:0072698) |
| Protein Localization to Microtubule (GO:0035372) |
| Positive Regulation of Vasculogenesis (GO:2001214) |
| Positive Regulation of Tissue Remodeling (GO:0034105) |
| Interleukin-15-Mediated Signaling Pathway (GO:0035723) |
| Positive Regulation of Sodium Ion Transmembrane Transporter Activity (GO:2000651) |
| Positive Regulation of Protein Localization to Endosome (GO:1905668) |
| Positive Regulation of Protein Localization to Early Endosome (GO:1902966) |
| Positive Regulation of Cardiocyte Differentiation (GO:1905209) |
| Response to Prostaglandin (GO:0034694) |
| Positive Regulation of Acrosome Reaction (GO:2000344) |
| Cellular Response to Interleukin-15 (GO:0071350) |
| Regulation of Translation in Response to Endoplasmic Reticulum Stress (GO:0036490) |
| Regulation of Protein Localization to Early Endosome (GO:1902965) |
| Nitric oxide-cGMP-mediated Signaling (GO:0038060) |
| Astral Microtubule Organization (GO:0030953) |
| Activation of NF-kappaB-inducing Kinase Activity (GO:0007250) |
| Regulation of Melanin Biosynthetic Process (GO:0048021) |
| Negative Regulation of Protein Exit From Endoplasmic Reticulum (GO:0070862) |
| Negative Regulation of Potassium Ion Transmembrane Transporter Activity (GO:1901017) |

| |
|---|
| NAD Biosynthesis via Nicotinamide Riboside Salvage Pathway (GO:0034356) |
| Regulation of Complement-Dependent Cytotoxicity (GO:1903659) |
| Neurogenesis (GO:0022008) |
| Regulation of Protein Serine/Threonine Kinase Activity (GO:0071900) |
| Cell-Cell Junction Assembly (GO:0007043) |
| Actin Filament Organization (GO:0007015) |
| Ras Protein Signal Transduction (GO:0007265) |
| Regulation of Cell Communication (GO:0010646) |
| Neuron Development (GO:0048666) |
| Epithelium Development (GO:0060429) |
| Skin Development (GO:0043588) |
| Positive Regulation of Intracellular Signal Transduction (GO:1902533) |
| Regulation of Heart Contraction (GO:0008016) |
| Fatty Acid Biosynthetic Process (GO:0006633) |
| Myelin Assembly (GO:0032288) |
| Receptor-Mediated Endocytosis of Virus by Host Cell (GO:0019065) |
| Positive Regulation of Nitric-Oxide Synthase Activity (GO:0051000) |
| Surfactant Homeostasis (GO:0043129) |
| Embryonic Appendage Morphogenesis (GO:0035113) |
| Positive Regulation of Chondrocyte Differentiation (GO:0032332) |
| Cellular Response to Interleukin-12 (GO:0071349) |
| Positive Regulation of Hemostasis (GO:1900048) |
| Atrial Septum Morphogenesis (GO:0060413) |
| Angiotensin-Activated Signaling Pathway (GO:0038166) |
| Negative Regulation of Protein Maturation (GO:1903318) |
| Cellular Response to Increased Oxygen Levels (GO:0036295) |
| NAD Biosynthetic Process via the Salvage Pathway (GO:0034355) |
| Regulation of Granulocyte Differentiation (GO:0030852) |
| Regulation of Complement Activation (GO:0030449) |

| p-value | q-value |
|---|---|
| 4,47619E-07 | 0,000499095 |
| 1,05794E-05 | 0,005898005 |
| 2,06272E-05 | 0,007666459 |
| 3,31288E-05 | 0,008686776 |
| 4,50012E-05 | 0,008686776 |
| 6,8721E-05 | 0,008686776 |
| 6,8721E-05 | 0,008686776 |
| 6,8721E-05 | 0,008686776 |
| 7,01175E-05 | 0,008686776 |
| 8,0899E-05 | 0,008836341 |
| 8,71747E-05 | 0,008836341 |
| 0,00013126 | 0,012196242 |
| 0,0001468 | 0,012509866 |
| 0,000160215 | 0,012509866 |
| 0,000168294 | 0,012509866 |
| 0,000184826 | 0,012880092 |
| 0,000241122 | 0,014150058 |
| 0,000241122 | 0,014150058 |
| 0,000241122 | 0,014150058 |
| 0,000254988 | 0,014215589 |
| 0,000307457 | 0,016324495 |
| 0,000341749 | 0,017001955 |
| 0,00038121 | 0,017001955 |
| 0,00038121 | 0,017001955 |
| 0,00038121 | 0,017001955 |
| 0,000404227 | 0,017325943 |
| 0,000419552 | 0,017325943 |
| 0,000473053 | 0,017483034 |
| 0,000476676 | 0,017483034 |
| 0,000529832 | 0,017483034 |
| 0,000531918 | 0,017483034 |
| 0,000531918 | 0,017483034 |
| 0,000531918 | 0,017483034 |
| 0,000533115 | 0,017483034 |
| 0,000628696 | 0,020028465 |
| 0,000714515 | 0,022130111 |
| 0,000822961 | 0,024800043 |
| 0,000954292 | 0,028000947 |
| 0,001070798 | 0,029958867 |
| 0,001109419 | 0,029958867 |
| 0,001128495 | 0,029958867 |
| 0,001128495 | 0,029958867 |
| 0,001192596 | 0,030924283 |
| 0,001286366 | 0,032597696 |
| 0,00137051 | 0,032612324 |

| | |
|---|---|
| 0,00137051 | 0,032612324 |
| 0,00137469 | 0,032612324 |
| 0,001564269 | 0,035075103 |
| 0,001595575 | 0,035075103 |
| 0,001644149 | 0,035075103 |
| 0,001644149 | 0,035075103 |
| 0,001656319 | 0,035075103 |
| 0,001667247 | 0,035075103 |
| 0,00184036 | 0,036924674 |
| 0,001936634 | 0,036924674 |
| 0,001936634 | 0,036924674 |
| 0,001936634 | 0,036924674 |
| 0,002005288 | 0,036924674 |
| 0,002121213 | 0,036924674 |
| 0,002121213 | 0,036924674 |
| 0,002121213 | 0,036924674 |
| 0,002251908 | 0,036924674 |
| 0,002251908 | 0,036924674 |
| 0,002251908 | 0,036924674 |
| 0,002251908 | 0,036924674 |
| 0,002251908 | 0,036924674 |
| 0,002251908 | 0,036924674 |
| 0,002251908 | 0,036924674 |
| 0,002531026 | 0,040899917 |
| 0,002589736 | 0,041250787 |
| 0,002949882 | 0,043854916 |
| 0,002949882 | 0,043854916 |
| 0,002949882 | 0,043854916 |
| 0,002949882 | 0,043854916 |
| 0,002949882 | 0,043854916 |
| 0,003284987 | 0,046029184 |
| 0,003331995 | 0,046029184 |
| 0,003332117 | 0,046029184 |
| 0,003332117 | 0,046029184 |
| 0,003332117 | 0,046029184 |
| 0,003343824 | 0,046029184 |
| 0,003542484 | 0,048169142 |
| 0,003736209 | 0,049593721 |
| 0,003736209 | 0,049593721 |
| 0,003966802 | 0,052035106 |
| 0,004080006 | 0,052897749 |
| 0,004282859 | 0,054889518 |
| 0,004473643 | 0,056683092 |
| 0,004539041 | 0,05686552 |
| 0,00463022 | 0,057363278 |
| 0,004973588 | 0,060940117 |

| | |
|---|---|
| 0,005198643 | 0,062327819 |
| 0,005198643 | 0,062327819 |
| 0,005504505 | 0,065216214 |
| 0,005566608 | 0,065216214 |
| 0,005615028 | 0,065216214 |
| 0,005726322 | 0,065780622 |
| 0,005805885 | 0,065780622 |
| 0,005876904 | 0,065780622 |
| 0,005953434 | 0,065780622 |
| 0,006048668 | 0,065780622 |
| 0,006076596 | 0,065780622 |
| 0,006076596 | 0,065780622 |
| 0,006185875 | 0,066261842 |
| 0,006239904 | 0,066261842 |
| 0,006423676 | 0,067569799 |
| 0,00651899 | 0,067577803 |
| 0,006666868 | 0,067577803 |
| 0,006666868 | 0,067577803 |
| 0,006666868 | 0,067577803 |
| 0,006915482 | 0,069466331 |
| 0,007728761 | 0,076261664 |
| 0,007728761 | 0,076261664 |
| 0,007834288 | 0,076624837 |
| 0,007964723 | 0,077223183 |
| 0,008319495 | 0,078592382 |
| 0,008319495 | 0,078592382 |
| 0,008319495 | 0,078592382 |
| 0,008387887 | 0,078592382 |
| 0,008809923 | 0,080917684 |
| 0,008809923 | 0,080917684 |
| 0,008929878 | 0,080917684 |
| 0,008929878 | 0,080917684 |
| 0,008998917 | 0,080917684 |
| 0,00914078 | 0,081535758 |
| 0,009537357 | 0,08392964 |
| 0,009559699 | 0,08392964 |
| 0,01001579 | 0,087246918 |
| 0,010208747 | 0,08755964 |
| 0,010208747 | 0,08755964 |
| 0,010876815 | 0,091241622 |
| 0,010876815 | 0,091241622 |
| 0,01088353 | 0,091241622 |
| 0,010980135 | 0,09136456 |
| 0,011563694 | 0,093431294 |
| 0,011563694 | 0,093431294 |
| 0,011563694 | 0,093431294 |

| | |
|---|---|
| 0,011563694 | 0,093431294 |
| 0,011873388 | 0,095243365 |
| 0,011996428 | 0,095542982 |
| 0,01226918 | 0,096270618 |
| 0,01226918 | 0,096270618 |
| 0,012346815 | 0,096270618 |
| 0,012993069 | 0,099227886 |
| 0,012993069 | 0,099227886 |
| 0,012993069 | 0,099227886 |
| 0,013550485 | 0,100618666 |
| 0,013735158 | 0,100618666 |
| 0,013735158 | 0,100618666 |
| 0,013735158 | 0,100618666 |
| 0,013780074 | 0,100618666 |
| 0,013780074 | 0,100618666 |
| 0,013806866 | 0,100618666 |
| 0,014495248 | 0,104272264 |
| 0,014495248 | 0,104272264 |
| 0,015737544 | 0,112483088 |
| 0,016068632 | 0,114117988 |
| 0,016881532 | 0,119132333 |
| 0,017437492 | 0,12151752 |
| 0,017437492 | 0,12151752 |
| 0,017711646 | 0,122661399 |
| 0,017966924 | 0,123661234 |
| 0,018089181 | 0,123738874 |
| 0,018309776 | 0,124484151 |
| 0,018558779 | 0,125412354 |
| 0,01942274 | 0,129678772 |
| 0,01942274 | 0,129678772 |
| 0,021200387 | 0,137300291 |
| 0,021200387 | 0,137300291 |
| 0,022038483 | 0,137300291 |
| 0,022113697 | 0,137300291 |
| 0,022113697 | 0,137300291 |
| 0,022117871 | 0,137300291 |
| 0,022950624 | 0,137300291 |
| 0,023043083 | 0,137300291 |
| 0,023043083 | 0,137300291 |
| 0,023043083 | 0,137300291 |
| 0,023988361 | 0,137300291 |
| 0,024046855 | 0,137300291 |
| 0,02433493 | 0,137300291 |
| 0,02474514 | 0,137300291 |
| 0,025243551 | 0,137300291 |
| 0,025243551 | 0,137300291 |

| | |
|---|---|
| 0,025243551 | 0,137300291 |
| 0,025243551 | 0,137300291 |
| 0,025243551 | 0,137300291 |
| 0,025243551 | 0,137300291 |
| 0,025243551 | 0,137300291 |
| 0,025243551 | 0,137300291 |
| 0,025243551 | 0,137300291 |
| 0,025243551 | 0,137300291 |
| 0,025243551 | 0,137300291 |
| 0,025243551 | 0,137300291 |
| 0,025243551 | 0,137300291 |
| 0,025243551 | 0,137300291 |
| 0,025243551 | 0,137300291 |
| 0,025243551 | 0,137300291 |
| 0,025243551 | 0,137300291 |
| 0,025243551 | 0,137300291 |
| 0,025243551 | 0,137300291 |
| 0,025243551 | 0,137300291 |
| 0,025243551 | 0,137300291 |
| 0,025243551 | 0,137300291 |
| 0,025243551 | 0,137300291 |
| 0,025243551 | 0,137300291 |
| 0,025925864 | 0,138977587 |
| 0,025925864 | 0,138977587 |
| 0,025925864 | 0,138977587 |
| 0,026917726 | 0,140378663 |
| 0,026917726 | 0,140378663 |
| 0,027924758 | 0,140378663 |
| 0,027924758 | 0,140378663 |
| 0,027924758 | 0,140378663 |
| 0,028946781 | 0,140378663 |
| 0,028946781 | 0,140378663 |
| 0,028946781 | 0,140378663 |
| 0,029983621 | 0,140378663 |
| 0,029983621 | 0,140378663 |
| 0,029983621 | 0,140378663 |
| 0,029983621 | 0,140378663 |
| 0,030064437 | 0,140378663 |
| 0,030216035 | 0,140378663 |
| 0,030216035 | 0,140378663 |
| 0,030216035 | 0,140378663 |
| 0,030216035 | 0,140378663 |
| 0,030216035 | 0,140378663 |
| 0,030216035 | 0,140378663 |
| 0,030216035 | 0,140378663 |
| 0,030216035 | 0,140378663 |

| | |
|---|---|
| 0,030216035 | 0,140378663 |
| 0,030216035 | 0,140378663 |
| 0,030216035 | 0,140378663 |
| 0,030216035 | 0,140378663 |
| 0,030216035 | 0,140378663 |
| 0,030216035 | 0,140378663 |
| 0,030216035 | 0,140378663 |
| 0,030216035 | 0,140378663 |
| 0,030216035 | 0,140378663 |
| 0,030216035 | 0,140378663 |
| 0,030216035 | 0,140378663 |
| 0,03210105 | 0,147951673 |
| 0,032431716 | 0,147951673 |
| 0,033181295 | 0,147951673 |
| 0,033181295 | 0,147951673 |
| 0,033181295 | 0,147951673 |
| 0,034269438 | 0,147951673 |
| 0,034269438 | 0,147951673 |
| 0,034275667 | 0,147951673 |
| 0,034275667 | 0,147951673 |
| 0,035163402 | 0,147951673 |
| 0,035163402 | 0,147951673 |
| 0,035163402 | 0,147951673 |
| 0,035163402 | 0,147951673 |
| 0,035163402 | 0,147951673 |
| 0,035163402 | 0,147951673 |
| 0,035163402 | 0,147951673 |
| 0,035163402 | 0,147951673 |
| 0,035163402 | 0,147951673 |
| 0,035163402 | 0,147951673 |
| 0,035163402 | 0,147951673 |
| 0,035163402 | 0,147951673 |
| 0,035163402 | 0,147951673 |
| 0,035163402 | 0,147951673 |
| 0,035163402 | 0,147951673 |
| 0,035163402 | 0,147951673 |
| 0,037641852 | 0,153067267 |
| 0,038763518 | 0,153067267 |
| 0,040085777 | 0,153067267 |
| 0,040085777 | 0,153067267 |
| 0,040085777 | 0,153067267 |
| 0,040085777 | 0,153067267 |
| 0,040085777 | 0,153067267 |
| 0,040085777 | 0,153067267 |
| 0,040085777 | 0,153067267 |
| 0,040085777 | 0,153067267 |

| | |
|---|---|
| 0,040085777 | 0,153067267 |
| 0,040085777 | 0,153067267 |
| 0,040085777 | 0,153067267 |
| 0,040085777 | 0,153067267 |
| 0,040085777 | 0,153067267 |
| 0,040085777 | 0,153067267 |
| 0,040085777 | 0,153067267 |
| 0,040085777 | 0,153067267 |
| 0,040085777 | 0,153067267 |
| 0,040085777 | 0,153067267 |
| 0,040085777 | 0,153067267 |
| 0,040085777 | 0,153067267 |
| 0,040085777 | 0,153067267 |
| 0,040085777 | 0,153067267 |
| 0,040085777 | 0,153067267 |
| 0,040085777 | 0,153067267 |
| 0,040085777 | 0,153067267 |
| 0,042317756 | 0,15528286 |
| 0,043519159 | 0,15528286 |
| 0,043519159 | 0,15528286 |
| 0,043543016 | 0,15528286 |
| 0,043543016 | 0,15528286 |
| 0,044164943 | 0,15528286 |
| 0,044983286 | 0,15528286 |
| 0,044983286 | 0,15528286 |
| 0,044983286 | 0,15528286 |
| 0,044983286 | 0,15528286 |
| 0,044983286 | 0,15528286 |
| 0,044983286 | 0,15528286 |
| 0,044983286 | 0,15528286 |
| 0,044983286 | 0,15528286 |
| 0,044983286 | 0,15528286 |
| 0,044983286 | 0,15528286 |
| 0,044983286 | 0,15528286 |
| 0,044983286 | 0,15528286 |
| 0,044983286 | 0,15528286 |
| 0,044983286 | 0,15528286 |
| 0,044983286 | 0,15528286 |
| 0,044983286 | 0,15528286 |
| 0,044983286 | 0,15528286 |
| 0,044983286 | 0,15528286 |
| 0,044983286 | 0,15528286 |
| 0,044983286 | 0,15528286 |
| 0,044983286 | 0,15528286 |
| 0,044983286 | 0,15528286 |
| 0,044983286 | 0,15528286 |
| 0,044983286 | 0,15528286 |
| 0,044983286 | 0,15528286 |

| | |
|---|---|
| 0,044983286 | 0,15528286 |
| 0,044983286 | 0,15528286 |
| 0,04595976 | 0,157193658 |
| 0,04595976 | 0,157193658 |
| 0,04595976 | 0,157193658 |
| 0,04712923 | 0,158827136 |
| 0,047198645 | 0,158827136 |
| 0,047198645 | 0,158827136 |
| 0,047863512 | 0,158827136 |
| 0,047863512 | 0,158827136 |
| 0,048449712 | 0,158827136 |
| 0,049216687 | 0,158827136 |
| 0,049712809 | 0,158827136 |
| 0,049712809 | 0,158827136 |
| 0,049856052 | 0,158827136 |
| 0,049856052 | 0,158827136 |
| 0,049856052 | 0,158827136 |
| 0,049856052 | 0,158827136 |
| 0,049856052 | 0,158827136 |
| 0,049856052 | 0,158827136 |
| 0,049856052 | 0,158827136 |
| 0,049856052 | 0,158827136 |
| 0,049856052 | 0,158827136 |
| 0,049856052 | 0,158827136 |
| 0,049856052 | 0,158827136 |
| 0,049856052 | 0,158827136 |
| 0,049856052 | 0,158827136 |
| 0,049856052 | 0,158827136 |
| 0,049856052 | 0,158827136 |
