## Supplemental Table S15 for "Temporal analysis of tear fluid proteome reveals critical corneal repair events after photorefractive surgery"

| term |
| --- |
| Cadherin Binding (GO:0045296) |
| Endopeptidase Inhibitor Activity (GO:0004866) |
| Exopeptidase Activity (GO:0008238) |
| Peptidase Inhibitor Activity (GO:0030414) |
| Endopeptidase Regulator Activity (GO:0061135) |
| Calcium Ion Binding (GO:0005509) |
| Metal Ion Binding (GO:0046872) |
| Actin Binding (GO:0003779) |
| Virus Receptor Activity (GO:0001618) |
| Dopamine Receptor Binding (GO:0050780) |
| Monocarboxylic Acid Transmembrane Transporter Activity (GO:0008028) |
| Adrenergic Receptor Binding (GO:0031690) |
| Ribonucleoside Triphosphate Phosphatase Activity (GO:0017111) |
| Cysteine-Type Endopeptidase Inhibitor Activity (GO:0004869) |
| Chemorepellent Activity (GO:0045499) |
| Phosphatidylinositol-4,5-Bisphosphate Binding (GO:0005546) |
| Aminopeptidase Activity (GO:0004177) |
| GTPase Activity (GO:0003924) |
| Phosphatidylinositol Bisphosphate Binding (GO:1902936) |
| Growth Factor Receptor Binding (GO:0070851) |
| ATP Binding (GO:0005524) |
| Phosphoric Diester Hydrolase Activity (GO:0008081) |
| Protein Homodimerization Activity (GO:0042803) |
| Adenyl Ribonucleotide Binding (GO:0032559) |
| Arylesterase Activity (GO:0004064) |
| Phospholipase D Activity (GO:0004630) |
| Iodide Transmembrane Transporter Activity (GO:0015111) |
| Cyclase Activator Activity (GO:0010853) |
| Secondary Active Monocarboxylate Transmembrane Transporter Activity (GO:0015355) |
| Icosatetraenoic Acid Binding (GO:0050543) |
| Arachidonate Binding (GO:0050544) |
| Kinase Binding (GO:0019900) |
| Omega Peptidase Activity (GO:0008242) |
| G Protein-Coupled Glutamate Receptor Binding (GO:0035256) |
| Icosanoid Binding (GO:0050542) |
| Signal Recognition Particle Binding (GO:0005047) |
| Protein Kinase Inhibitor Activity (GO:0004860) |
| Pyrophosphatase Activity (GO:0016462) |
| Serine-Type Endopeptidase Inhibitor Activity (GO:0004867) |
| Cysteine-Type Peptidase Activity (GO:0008234) |
| UDP Phosphatase Activity (GO:0045134) |
| Adenylate Cyclase Regulator Activity (GO:0010854) |
| Tumor Necrosis Factor Receptor Activity (GO:0005031) |
| Type II Transforming Growth Factor Beta Receptor Binding (GO:0005114) |
| GDP Phosphatase Activity (GO:0004382) |

| |
|---|
| Channel Activator Activity (GO:0099103) |
| Receptor Ligand Activity (GO:0048018) |
| Protein Serine/Threonine Kinase Activator Activity (GO:0043539) |
| Dipeptidyl-Peptidase Activity (GO:0008239) |

| p-value | q-value |
|---|---|
| 7,71738E-08 | 1,29652E-05 |
| 1,70419E-07 | 1,43152E-05 |
| 7,38513E-06 | 0,000413568 |
| 5,06009E-05 | 0,002125238 |
| 0,000113257 | 0,00380545 |
| 0,00025484 | 0,007135516 |
| 0,000305271 | 0,007326508 |
| 0,000394849 | 0,007976735 |
| 0,000427325 | 0,007976735 |
| 0,000905807 | 0,015217552 |
| 0,001667247 | 0,02546341 |
| 0,002589736 | 0,036256297 |
| 0,002812856 | 0,036350757 |
| 0,005077354 | 0,060928243 |
| 0,006607095 | 0,072612564 |
| 0,006915482 | 0,072612564 |
| 0,008929878 | 0,088248206 |
| 0,011735286 | 0,109529335 |
| 0,013065112 | 0,115523094 |
| 0,014572179 | 0,116637207 |
| 0,014579651 | 0,116637207 |
| 0,021200387 | 0,146238504 |
| 0,022964583 | 0,146238504 |
| 0,023487395 | 0,146238504 |
| 0,025243551 | 0,146238504 |
| 0,025243551 | 0,146238504 |
| 0,025243551 | 0,146238504 |
| 0,025243551 | 0,146238504 |
| 0,025243551 | 0,146238504 |
| 0,030216035 | 0,1566142 |
| 0,030216035 | 0,1566142 |
| 0,033848256 | 0,1566142 |
| 0,035163402 | 0,1566142 |
| 0,035163402 | 0,1566142 |
| 0,035163402 | 0,1566142 |
| 0,035163402 | 0,1566142 |
| 0,035383995 | 0,1566142 |
| 0,036506112 | 0,1566142 |
| 0,037641852 | 0,1566142 |
| 0,039428925 | 0,1566142 |
| 0,040085777 | 0,1566142 |
| 0,040085777 | 0,1566142 |
| 0,040085777 | 0,1566142 |
| 0,044983286 | 0,161278952 |
| 0,044983286 | 0,161278952 |

| | |
|---|---|
| 0,044983286 | 0,161278952 |
| 0,045119707 | 0,161278952 |
| 0,049712809 | 0,1707454 |
| 0,049856052 | 0,1707454 |
