## Supplemental Table S16 for "Temporal analysis of tear fluid proteome reveals critical corneal repair events after photorefractive surgery"

| term |
| --- |
| Lewis Blood Group Biosynthesis |
| Blood Group Systems Biosynthesis |
| Reactions Specific to the Complex N-glycan Synthesis Pathway |
| Metabolism |
| Metabolism of Carbohydrates |
| Asparagine N-linked Glycosylation |
| N-glycan Antennae Elongation in the Medial trans-Golgi |
| Nucleotide Catabolism |
| Maturation of Spike Protein 9694548 |
| Intra-Golgi Traffic |
| Endosomal Vacuolar Pathway |
| Translation of Structural Proteins 9694635 |
| Transport to the Golgi and Subsequent Modification |
| Purine Catabolism |
| Late SARS-CoV-2 Infection Events |
| Pre-NOTCH Processing in Golgi |
| Intra-Golgi and Retrograde Golgi-to-ER Traffic |
| Metabolism of Nucleotides |
| Metabolism of Proteins |
| Keratan Sulfate Biosynthesis |
| Post-translational Protein Modification |
| Diseases of Metabolism |
| Activation of Matrix Metalloproteinases |
| Sialic Acid Metabolism |
| Glycosaminoglycan Metabolism |
| Translation of Structural Proteins 9683701 |
| Keratan Sulfate Keratin Metabolism |
| Diseases of Glycosylation |
| SARS-CoV-2 Infection |
| Innate Immune System |
| Synthesis of Substrates in N-glycan Biosythesis |
| O-linked Glycosylation of Mucins |
| Defective LFNG Causes SCDO3 |
| N-glycan Trimming and Elongation in the cis-Golgi |
| Arachidonate Production From DAG |
| Biosynthesis of the N-glycan Precursor (Dolichol LLO) and Transfer to a Nascent Protein |
| Maturation of Spike Protein 9683686 |
| Pre-NOTCH Expression and Processing |
| Neutrophil Degranulation |
| Acyl Chain Remodeling of DAG and TAG |
| Antigen processing-Cross Presentation |
| SARS-CoV Infections |
| RUNX1 Regulates Transcription of Genes Involved in Differentiation of Keratinocytes |
| Defective CHST14 Causes EDS, Musculocontractural Type |
| TRAIL Signaling |

| |
|---|
| Synthesis of PG |
| Abacavir ADME |
| Immune System |
| Sphingolipid Metabolism |
| Drug ADME |
| Maturation of Protein 3A 9683673 |
| Maturation of Protein 3A 9694719 |
| Uptake of Dietary Cobalamins Into Enterocytes |
| O-linked Glycosylation |
| CASP8 Activity Is Inhibited |
| Dimerization of Procaspase-8 |
| Dermatan Sulfate Biosynthesis |
| Regulation by c-FLIP |
| Modulation by Mtb of Host Immune System |
| Pyrimidine Salvage |
| Ribavirin ADME |
| Nef Mediated Downregulation of MHC Class I Complex Cell Surface Expression |
| Pyrimidine Catabolism |
| Butyrophilin (BTN) Family Interactions |
| MHC Class II Antigen Presentation |
| Trafficking and Processing of Endosomal TLR |
| Nephron Development |
| Glycerophospholipid Biosynthesis |
| Glycogen Breakdown (Glycogenolysis) |
| Tryptophan Catabolism |
| Degradation of the Extracellular Matrix |
| Glycogen Storage Diseases |
| Membrane Trafficking |
| Regulation of MITF-M-dependent Genes Involved in Cell Cycle and Proliferation |
| Caspase Activation via Death Receptors in the Presence of Ligand |
| Regulation of MITF-M-dependent Genes Involved in Apoptosis |
| SARS-CoV-1 Infection |
| Diseases Associated With N-glycosylation of Proteins |
| Receptor-type Tyrosine-Protein Phosphatases |
| Synaptic Adhesion-Like Molecules |
| Cobalamin (Cbl, Vitamin B12) Transport and Metabolism |
| Nef-mediates Down Modulation of Cell Surface Receptors by Recruiting Them to Clathrin Adapters |
| Nucleotide Salvage |
| Azathioprine ADME |
| Disease |
| Triglyceride Catabolism |
| Basigin Interactions |
| Glycogen Metabolism |
| Termination of O-glycan Biosynthesis |

| p-value | q-value |
|---|---|
| 3,32469E-08 | 6,05093E-06 |
| 7,9047E-08 | 7,19328E-06 |
| 7,52333E-07 | 3,65217E-05 |
| 8,02675E-07 | 3,65217E-05 |
| 1,07643E-06 | 3,91821E-05 |
| 1,53967E-06 | 4,67033E-05 |
| 1,59624E-05 | 0,000415021 |
| 3,97109E-05 | 0,000903423 |
| 5,09834E-05 | 0,001030998 |
| 7,94139E-05 | 0,001445333 |
| 0,000229226 | 0,003522651 |
| 0,000232263 | 0,003522651 |
| 0,000417509 | 0,005702243 |
| 0,000469523 | 0,005702243 |
| 0,000469965 | 0,005702243 |
| 0,000527581 | 0,006001235 |
| 0,000591264 | 0,006330006 |
| 0,000849766 | 0,008592083 |
| 0,00119796 | 0,011475198 |
| 0,001287921 | 0,011532623 |
| 0,001330687 | 0,011532623 |
| 0,001539182 | 0,012733229 |
| 0,001788269 | 0,013055528 |
| 0,001788269 | 0,013055528 |
| 0,001793342 | 0,013055528 |
| 0,002010374 | 0,014072617 |
| 0,002126087 | 0,0143314 |
| 0,002666224 | 0,017330459 |
| 0,002856268 | 0,017925542 |
| 0,005340097 | 0,032396588 |
| 0,006579762 | 0,038564405 |
| 0,006780555 | 0,038564405 |
| 0,009464812 | 0,049217022 |
| 0,009464812 | 0,049217022 |
| 0,009464812 | 0,049217022 |
| 0,009877597 | 0,049936742 |
| 0,011347288 | 0,05581639 |
| 0,012406134 | 0,058407737 |
| 0,012515944 | 0,058407737 |
| 0,013226282 | 0,059750594 |
| 0,014326701 | 0,059750594 |
| 0,014361174 | 0,059750594 |
| 0,015101799 | 0,059750594 |
| 0,015101799 | 0,059750594 |
| 0,015101799 | 0,059750594 |

| | |
|---|---|
| 0,015101799 | 0,059750594 |
| 0,016973845 | 0,060786683 |
| 0,017159386 | 0,060786683 |
| 0,017586133 | 0,060786683 |
| 0,018209613 | 0,060786683 |
| 0,018842427 | 0,060786683 |
| 0,018842427 | 0,060786683 |
| 0,018842427 | 0,060786683 |
| 0,019808851 | 0,060786683 |
| 0,020707551 | 0,060786683 |
| 0,020707551 | 0,060786683 |
| 0,020707551 | 0,060786683 |
| 0,020707551 | 0,060786683 |
| 0,020707551 | 0,060786683 |
| 0,020707551 | 0,060786683 |
| 0,020707551 | 0,060786683 |
| 0,020707551 | 0,060786683 |
| 0,022569223 | 0,064181229 |
| 0,022569223 | 0,064181229 |
| 0,023877239 | 0,065803931 |
| 0,02442745 | 0,065803931 |
| 0,02442745 | 0,065803931 |
| 0,024586084 | 0,065803931 |
| 0,026282236 | 0,068333815 |
| 0,026282236 | 0,068333815 |
| 0,02901727 | 0,074382298 |
| 0,029981516 | 0,075786609 |
| 0,031348913 | 0,078157564 |
| 0,03182602 | 0,078274807 |
| 0,03366711 | 0,081698854 |
| 0,035504791 | 0,08460669 |
| 0,035795138 | 0,08460669 |
| 0,037339069 | 0,086021652 |
| 0,037339069 | 0,086021652 |
| 0,03916995 | 0,089111637 |
| 0,040997441 | 0,090994321 |
| 0,040997441 | 0,090994321 |
| 0,042821548 | 0,09278002 |
| 0,042821548 | 0,09278002 |
| 0,04385867 | 0,093909151 |
| 0,044642276 | 0,094475514 |
| 0,046459632 | 0,096086965 |
| 0,046459632 | 0,096086965 |
| 0,048273621 | 0,098716843 |
