## Supplemental Table S17 for "Temporal analysis of tear fluid proteome reveals critical corneal repair events after photorefractive surgery"

| term |
| --- |
| Protein Glycosylation (GO:0006486) |
| Macromolecule Glycosylation (GO:0043413) |
| Oligosaccharide Biosynthetic Process (GO:0009312) |
| Carbohydrate Biosynthetic Process (GO:0016051) |
| Glycoprotein Biosynthetic Process (GO:0009101) |
| Protein O-linked Glycosylation (GO:0006493) |
| Oligosaccharide Metabolic Process (GO:0009311) |
| Purine Ribonucleotide Catabolic Process (GO:0009154) |
| Protein N-linked Glycosylation (GO:0006487) |
| N-glycan Processing (GO:0006491) |
| Purine Ribonucleoside Catabolic Process (GO:0046130) |
| Purine Ribonucleoside Metabolic Process (GO:0046128) |
| Purine Ribonucleoside Monophosphate Catabolic Process (GO:0009169) |
| Proteoglycan Biosynthetic Process (GO:0030166) |
| Regulation of Receptor Binding (GO:1900120) |
| IMP Metabolic Process (GO:0046040) |
| Negative Regulation of Receptor Binding (GO:1900121) |
| Ceramide Metabolic Process (GO:0006672) |
| Fucosylation (GO:0036065) |
| Keratan Sulfate Proteoglycan Biosynthetic Process (GO:0018146) |
| Sphingolipid Metabolic Process (GO:0006665) |
| Keratan Sulfate Proteoglycan Metabolic Process (GO:0042339) |
| Sphingolipid Biosynthetic Process (GO:0030148) |
| Antigen Processing and Presentation of Exogenous Peptide Antigen via MHC Class II (GO:0019886) |
| Proteoglycan Metabolic Process (GO:0006029) |
| Antigen Processing and Presentation of Peptide Antigen via MHC Class II (GO:0002495) |
| Antigen Processing and Presentation of Exogenous Peptide Antigen (GO:0002478) |
| Lytic Vacuole Organization (GO:0080171) |
| O-glycan Processing (GO:0016266) |
| Lysosome Organization (GO:0007040) |
| Membrane Lipid Biosynthetic Process (GO:0046467) |
| Negative Regulation of Protein Binding (GO:0032091) |
| Glycoprotein Metabolic Process (GO:0009100) |
| Glycolipid Biosynthetic Process (GO:0009247) |
| Purine Ribonucleotide Metabolic Process (GO:0009150) |
| Protein Metabolic Process (GO:0019538) |
| Defense Response to Bacterium (GO:0042742) |
| Cell Surface Receptor Protein Tyrosine Phosphatase Signaling Pathway (GO:0007185) |
| Glycosylceramide Catabolic Process (GO:0046477) |
| Dermatan Sulfate Proteoglycan Metabolic Process (GO:0050655) |
| Proteolysis Involved in Protein Catabolic Process (GO:0051603) |
| Macromolecule Modification (GO:0043412) |
| Marginal Zone B Cell Differentiation (GO:0002315) |
| Glucan Catabolic Process (GO:0009251) |
| Monoacylglycerol Catabolic Process (GO:0052651) |

| |
|---|
| Adenosine Metabolic Process (GO:0046085) |
| Regulation of Protein Lipidation (GO:1903059) |
| Thyroid Hormone Transport (GO:0070327) |
| Purine Nucleotide Catabolic Process (GO:0006195) |
| Sulfur Amino Acid Transport (GO:0000101) |
| Regulation of Epithelial Cell Proliferation (GO:0050678) |
| Lipid Glycosylation (GO:0030259) |
| Proline Transport (GO:0015824) |
| Positive Regulation of Epithelial Cell Proliferation Involved in Wound Healing (GO:0060054) |
| GMP Metabolic Process (GO:0046037) |
| Amino Sugar Metabolic Process (GO:0006040) |
| TRAIL-activated Apoptotic Signaling Pathway (GO:0036462) |
| Aromatic Amino Acid Transport (GO:0015801) |
| Regulation of Lysosomal Protein Catabolic Process (GO:1905165) |
| Sialylation (GO:0097503) |
| Hormone Transport (GO:0009914) |
| Fucose Catabolic Process (GO:0019317) |
| L-fucose Metabolic Process (GO:0042354) |
| L-fucose Catabolic Process (GO:0042355) |
| Protein O-linked Glycosylation via Threonine (GO:0018243) |
| Protein O-linked Glycosylation via Serine (GO:0018242) |
| L-leucine Transport (GO:0015820) |
| Regulation of Macroautophagy (GO:0016241) |
| Protein Catabolic Process (GO:0030163) |
| Branched-Chain Amino Acid Transport (GO:0015803) |
| Positive Regulation of Hemostasis (GO:1900048) |
| N-acetylneuraminate Metabolic Process (GO:0006054) |
| Positive Regulation of Cellular Senescence (GO:2000774) |
| Protein Modification Process (GO:0036211) |
| AMP Metabolic Process (GO:0046033) |
| Negative Regulation of Lipopolysaccharide-Mediated Signaling Pathway (GO:0031665) |
| Purine Ribonucleoside Monophosphate Metabolic Process (GO:0009167) |
| Proteolysis (GO:0006508) |
| Response to Iron Ion (GO:0010039) |
| Monoacylglycerol Metabolic Process (GO:0046462) |
| Positive Regulation of Coagulation (GO:0050820) |
| Mature B Cell Differentiation Involved in Immune Response (GO:0002313) |
| calcineurin-NFAT Signaling Cascade (GO:0033173) |
| Negative Regulation of Defense Response to Virus (GO:0050687) |
| Regulation of Sensory Perception (GO:0051931) |
| Protein Alpha-1,2-Demannosylation (GO:0036508) |
| Glycogen Catabolic Process (GO:0005980) |
| Response to Interleukin-6 (GO:0070741) |
| Positive Regulation of Protein-Containing Complex Disassembly (GO:0043243) |
| Sphingoid Biosynthetic Process (GO:0046520) |
| Sphingosine Biosynthetic Process (GO:0046512) |

| |
|---|
| Regulation of Sensory Perception of Pain (GO:0051930) |
| Ganglioside Biosynthetic Process (GO:0001574) |
| Positive Regulation of Secretion (GO:0051047) |
| Establishment of Protein Localization to Endoplasmic Reticulum (GO:0072599) |
| Glycolipid Metabolic Process (GO:0006664) |
| Calcineurin-Mediated Signaling (GO:0097720) |
| Synaptic Membrane Adhesion (GO:0099560) |
| Regulation of Secretion (GO:0051046) |
| Diol Biosynthetic Process (GO:0034312) |
| Regulation of Protein-Containing Complex Disassembly (GO:0043244) |
| Glycerolipid Catabolic Process (GO:0046503) |
| Negative Regulation of Nervous System Development (GO:0051961) |
| Negative Regulation of Insulin Secretion (GO:0046676) |
| Negative Regulation of Response to External Stimulus (GO:0032102) |
| Negative Regulation of Peptide Hormone Secretion (GO:0090278) |
| Positive Regulation of calcineurin-NFAT Signaling Cascade (GO:0070886) |
| Positive Regulation of Calcineurin-Mediated Signaling (GO:0106058) |
| Antigen Processing and Presentation of Peptide Antigen via MHC Class I (GO:0002474) |
| Striated Muscle Cell Differentiation (GO:0051146) |
| Sphingosine Metabolic Process (GO:0006670) |
| Positive Regulation of Release of Cytochrome C From Mitochondria (GO:0090200) |
| Glycogen Metabolic Process (GO:0005977) |
| Positive Regulation of Blood Coagulation (GO:0030194) |
| Regulation of Membrane Depolarization (GO:0003254) |
| Antigen Processing and Presentation of Endogenous Peptide Antigen (GO:0002483) |
| Negative Regulation of Cell Development (GO:0010721) |
| Epithelial Structure Maintenance (GO:0010669) |
| Peptidyl-Tyrosine Dephosphorylation (GO:0035335) |
| Negative Regulation of Neuron Differentiation (GO:0045665) |
| Cellular Response to Interleukin-6 (GO:0071354) |
| Protein Targeting to ER (GO:0045047) |
| Regulation of Lipopolysaccharide-Mediated Signaling Pathway (GO:0031664) |
| Response to dsRNA (GO:0043331) |
| Myotube Differentiation (GO:0014902) |
| Amyloid Fibril Formation (GO:1990000) |

| p-value | q-value |
|---|---|
| 9,51625E-09 | 3,0452E-06 |
| 4,15182E-08 | 6,64292E-06 |
| 7,9047E-08 | 8,43168E-06 |
| 2,1946E-07 | 1,75568E-05 |
| 3,15495E-07 | 2,01917E-05 |
| 7,77937E-07 | 4,149E-05 |
| 2,02967E-06 | 9,2785E-05 |
| 5,06253E-06 | 0,000202501 |
| 6,48505E-06 | 0,00023058 |
| 8,21905E-06 | 0,00026301 |
| 3,50236E-05 | 0,001018868 |
| 5,24726E-05 | 0,001399269 |
| 7,33737E-05 | 0,001806122 |
| 0,000109753 | 0,002362024 |
| 0,000125483 | 0,002362024 |
| 0,000125483 | 0,002362024 |
| 0,000125483 | 0,002362024 |
| 0,000138821 | 0,002467925 |
| 0,000315297 | 0,005310273 |
| 0,000363369 | 0,005813906 |
| 0,000452934 | 0,006829427 |
| 0,000469523 | 0,006829427 |
| 0,000523512 | 0,007283641 |
| 0,001025837 | 0,013677823 |
| 0,001109993 | 0,014207916 |
| 0,001197359 | 0,014736722 |
| 0,001578669 | 0,018710149 |
| 0,002126087 | 0,024298134 |
| 0,002366776 | 0,026116144 |
| 0,003162119 | 0,033729267 |
| 0,003752501 | 0,038735493 |
| 0,004065467 | 0,040654673 |
| 0,004390165 | 0,042571294 |
| 0,004899012 | 0,046108352 |
| 0,005433758 | 0,04968007 |
| 0,005827128 | 0,051796696 |
| 0,008446659 | 0,073052187 |
| 0,009464812 | 0,075718495 |
| 0,009464812 | 0,075718495 |
| 0,009464812 | 0,075718495 |
| 0,010118819 | 0,077258131 |
| 0,011109726 | 0,077258131 |
| 0,011347288 | 0,077258131 |
| 0,011347288 | 0,077258131 |
| 0,011347288 | 0,077258131 |

| | |
|---|---|
| 0,011347288 | 0,077258131 |
| 0,011347288 | 0,077258131 |
| 0,013226282 | 0,08106911 |
| 0,013226282 | 0,08106911 |
| 0,013226282 | 0,08106911 |
| 0,013488696 | 0,08106911 |
| 0,015101799 | 0,08106911 |
| 0,015101799 | 0,08106911 |
| 0,015101799 | 0,08106911 |
| 0,015101799 | 0,08106911 |
| 0,015101799 | 0,08106911 |
| 0,015101799 | 0,08106911 |
| 0,015101799 | 0,08106911 |
| 0,016973845 | 0,08106911 |
| 0,016973845 | 0,08106911 |
| 0,016973845 | 0,08106911 |
| 0,016973845 | 0,08106911 |
| 0,016973845 | 0,08106911 |
| 0,016973845 | 0,08106911 |
| 0,016973845 | 0,08106911 |
| 0,016973845 | 0,08106911 |
| 0,016973845 | 0,08106911 |
| 0,017277905 | 0,081307786 |
| 0,017586133 | 0,081558876 |
| 0,018842427 | 0,082596941 |
| 0,018842427 | 0,082596941 |
| 0,018842427 | 0,082596941 |
| 0,018842427 | 0,082596941 |
| 0,019357322 | 0,083707338 |
| 0,020707551 | 0,088352219 |
| 0,022569223 | 0,090892836 |
| 0,022569223 | 0,090892836 |
| 0,022836924 | 0,090892836 |
| 0,02442745 | 0,090892836 |
| 0,02442745 | 0,090892836 |
| 0,02442745 | 0,090892836 |
| 0,02442745 | 0,090892836 |
| 0,02442745 | 0,090892836 |
| 0,02442745 | 0,090892836 |
| 0,02442745 | 0,090892836 |
| 0,02442745 | 0,090892836 |
| 0,026282236 | 0,092421051 |
| 0,026282236 | 0,092421051 |
| 0,026282236 | 0,092421051 |
| 0,026282236 | 0,092421051 |
| 0,026282236 | 0,092421051 |

| | |
|---|---|
| 0,02813359 | 0,095773922 |
| 0,02813359 | 0,095773922 |
| 0,02813359 | 0,095773922 |
| 0,029981516 | 0,099938385 |
| 0,029981516 | 0,099938385 |
| 0,03182602 | 0,102871985 |
| 0,03182602 | 0,102871985 |
| 0,03182602 | 0,102871985 |
| 0,03366711 | 0,103591108 |
| 0,03366711 | 0,103591108 |
| 0,03366711 | 0,103591108 |
| 0,03366711 | 0,103591108 |
| 0,03366711 | 0,103591108 |
| 0,034557981 | 0,104234248 |
| 0,035504791 | 0,104234248 |
| 0,035504791 | 0,104234248 |
| 0,035504791 | 0,104234248 |
| 0,035504791 | 0,104234248 |
| 0,037339069 | 0,104811421 |
| 0,037339069 | 0,104811421 |
| 0,037339069 | 0,104811421 |
| 0,037339069 | 0,104811421 |
| 0,037339069 | 0,104811421 |
| 0,03916995 | 0,108055035 |
| 0,03916995 | 0,108055035 |
| 0,040997441 | 0,11024522 |
| 0,040997441 | 0,11024522 |
| 0,040997441 | 0,11024522 |
| 0,042821548 | 0,112318814 |
| 0,042821548 | 0,112318814 |
| 0,042821548 | 0,112318814 |
| 0,044642276 | 0,114284226 |
| 0,044642276 | 0,114284226 |
| 0,044642276 | 0,114284226 |
| 0,048273621 | 0,122599673 |
