## Supplemental Table S18 for "Temporal analysis of tear fluid proteome reveals critical corneal repair events after photorefractive surgery"

| term |
| --- |
| Beta-Galactoside (CMP) Alpha-2,3-Sialyltransferase Activity (GO:0003836) |
| Alpha-(1->3)-Fucosyltransferase Activity (GO:0046920) |
| Alpha-Glucosidase Activity (GO:0090599) |
| Mannosidase Activity (GO:0015923) |
| Fucosyltransferase Activity (GO:0008417) |
| Sialyltransferase Activity (GO:0008373) |
| Hydrolase Activity, Hydrolyzing O-glycosyl Compounds (GO:0004553) |
| Hexosyltransferase Activity (GO:0016758) |
| Cysteine-Type Peptidase Activity (GO:0008234) |
| Cysteine-Type Endopeptidase Activity (GO:0004197) |
| L-leucine Transmembrane Transporter Activity (GO:0015190) |
| 4-galactosyl-N-acetylglucosaminide 3-alpha-L-fucosyltransferase Activity (GO:0017083) |
| Dermatan Sulfotransferase Activity (GO:0120534) |
| Phospholipase D Activity (GO:0004630) |
| Phospholipase Activity (GO:0004620) |
| Glucosidase Activity (GO:0015926) |
| Alpha-Mannosidase Activity (GO:0004559) |
| Mannosyl-Oligosaccharide 1,2-Alpha-Mannosidase Activity (GO:0004571) |
| Mannosyl-Oligosaccharide Mannosidase Activity (GO:0015924) |
| Glucosyltransferase Activity (GO:0046527) |
| L-alanine Transmembrane Transporter Activity (GO:0015180) |
| 5'-3' DNA Exonuclease Activity (GO:0035312) |
| Single-Stranded DNA Exodeoxyribonuclease Activity (GO:0008297) |
| Phosphate Ion Binding (GO:0042301) |
| Branched-Chain Amino Acid Transmembrane Transporter Activity (GO:0015658) |
| Aromatic Amino Acid Transmembrane Transporter Activity (GO:0015173) |
| N-acetyl-beta-D-glucosaminide Beta-(1,3)-Galactosyltransferase Activity (GO:0008499) |
| 5'-Nucleotidase Activity (GO:0008253) |
| Monoacylglycerol Lipase Activity (GO:0047372) |
| Beta-1,3-Galactosyltransferase Activity (GO:0048531) |
| Nucleotidase Activity (GO:0008252) |
| 2 Iron, 2 Sulfur Cluster Binding (GO:0051537) |
| Alanine Transmembrane Transporter Activity (GO:0022858) |
| Transmembrane Receptor Protein Phosphatase Activity (GO:0019198) |
| Transmembrane Receptor Protein Tyrosine Phosphatase Activity (GO:0005001) |
| Receptor Ligand Activity (GO:0048018) |
| Nucleobase-Containing Compound Kinase Activity (GO:0019205) |
| UDP-galactosyltransferase Activity (GO:0035250) |
| Lysophospholipase Activity (GO:0004622) |
| Protein Homodimerization Activity (GO:0042803) |
| MHC Class II Protein Complex Binding (GO:0023026) |
| Cytokine Activity (GO:0005125) |

| p-value | q-value |
|---|---|
| 5,24726E-05 | 0,002003148 |
| 7,33737E-05 | 0,002003148 |
| 7,33737E-05 | 0,002003148 |
| 9,77145E-05 | 0,002003148 |
| 0,000191251 | 0,00313651 |
| 0,000653598 | 0,008932511 |
| 0,000792737 | 0,008970563 |
| 0,000875177 | 0,008970563 |
| 0,002718224 | 0,024766041 |
| 0,008711027 | 0,048972506 |
| 0,009464812 | 0,048972506 |
| 0,009464812 | 0,048972506 |
| 0,009464812 | 0,048972506 |
| 0,009464812 | 0,048972506 |
| 0,010858117 | 0,048972506 |
| 0,011347288 | 0,048972506 |
| 0,011347288 | 0,048972506 |
| 0,011347288 | 0,048972506 |
| 0,011347288 | 0,048972506 |
| 0,013226282 | 0,054227754 |
| 0,015101799 | 0,055674212 |
| 0,015101799 | 0,055674212 |
| 0,016973845 | 0,055674212 |
| 0,016973845 | 0,055674212 |
| 0,016973845 | 0,055674212 |
| 0,018842427 | 0,059426117 |
| 0,020707551 | 0,0628896 |
| 0,022569223 | 0,065307375 |
| 0,026282236 | 0,065307375 |
| 0,026282236 | 0,065307375 |
| 0,026282236 | 0,065307375 |
| 0,026282236 | 0,065307375 |
| 0,026282236 | 0,065307375 |
| 0,02813359 | 0,065912981 |
| 0,02813359 | 0,065912981 |
| 0,035163277 | 0,080094131 |
| 0,037339069 | 0,080414602 |
| 0,03916995 | 0,080414602 |
| 0,03916995 | 0,080414602 |
| 0,039226635 | 0,080414602 |
| 0,044642276 | 0,089284552 |
| 0,048602873 | 0,094891323 |
